# Genome-wide genealogies reveal deep admixtures forming modern humans

**DOI:** 10.64898/2026.04.17.719197

**Authors:** Hrushikesh Loya, Anjali Gupta Hinch, Pier Francesco Palamara, Leo Speidel, Simon Robert Myers

## Abstract

Over the past decade, genomic modelling has revealed a rich tapestry of admixtures shaping present-day human populations. These have largely focused on the past few thousand years, when ancestral populations are either well characterised by present-day genomic diversity or directly observed through ancient DNA. Genomic modelling and fossil evidence have so far only provided a fragmented picture of the coexistence and mixing of human groups in the deeper past. Here, we propose a new method, GhostBuster, that leverages inferred genome-wide genealogies to detect admixture events of unsampled ghost populations, while simultaneously inferring accurate local ancestry. Local ancestry enables us to identify ancestry-specific genomic signatures that independently corroborate the events. We identify at least three waves of “back-to-Africa” migrations starting ∼14,000 years ago. Applying GhostBuster to deeper timescales reveals that modern humans were shaped by repeated episodes of mixture. Around 50,000 years ago, we identify a human lineage that expanded to form present-day non-Africans, while also expanding within Africa, mixing with the other local African group in varying proportions. These ancient groups help explain polygenic score portability differences within Africa, and exhibit differences in population size and recombination landscapes. Extending our analysis further back to between 300,000 and 1 million years ago reveals two deeply diverged ancestral lineages. These lineages evolved profoundly different recombination landscapes, with different PRDM9 alleles (PRDM9-A and C) and recombination hotspots. We demonstrate that both Neanderthals and ancestral modern humans are formed through a mixture of these two lineages, with no evidence of gene flow from the PRDM9-A-carrying group into Denisovans.

## Main

Modern population genomic methods have produced increasingly detailed reconstructions of recent human population history, yet leveraging genomic variation to reconstruct the deeper human past, where admixing groups are often unsampled, has remained challenging. Distinguishing among scenarios based on allele sharing patterns alone can be difficult, while linkage-disequilibrium–based signatures decay over time, and short, ancient ancestry tracts are difficult to detect. Ancient DNA has revealed numerous previously hidden mixture events over the past millennia^1^. However, ancient DNA is often challenging to sequence at deeper time depths and in warmer climates, due to high degradation rates.

Consequently, many questions about the deep population history of sub-Saharan Africa, a region of central importance for understanding human evolution, remain incompletely resolved^2^. A recent study argues for deep population structure within humans, involving two ancient lineages diverging more than 1.5 million years ago and admixing around 300k years ago, followed by diversification to form present-day populations^3^. Other models, though, suggest a complex structured metapopulation, connected by episodes of fragmentation and gene flow^4–6^. Additionally, several studies propose introgression from unsampled archaic hominins, or ghost populations^7–11^ and provide evidence for gene flow from early *Homo sapiens* into Neanderthals^12–14^.

Here, we introduce GhostBuster, a new approach that leverages genome-wide genealogies, which can now be inferred from genetic variation at scale^15–19^ to identify shared ancestral lineages up to millions of years into the past. GhostBuster decomposes the genome of a target individual into segments of distinct ancestries, based on local coalescence patterns. It fits a mixture model, whereby regions from each distinct ancestry coalesce with reference populations at different rates, which are directly identified from the data. This flexibility allows GhostBuster to identify source ancestries corresponding to new, unsampled ‘ghost’ groups, because reference populations are not required to exactly match these underlying sources. It additionally identifies local ancestry tracts deriving from each source component within target individuals. Through simulations and real-data examples, we demonstrate that GhostBuster can successfully recover signatures of both ancient and recent admixtures, accurately identifying source groups, admixture times, and the local ancestry of target individuals simultaneously.

Alongside this, we show that by identifying ancestry segments, GhostBuster allows us to identify mutational “signatures”, specific to particular ancestry backgrounds, to provide independent support for the detected events. These signatures are induced, for example, by ancestry-specific mutation- or recombination-rate landscapes, or population size differences. Elevated TCC to TTC substitutions have been previously observed in West Eurasian populations^20–22^. Others are connected to the recombination rate. In humans and other species^23–25^, recombination events induce overtransmission of weak-to-strong (A/T to G/C) mutations to offspring, a phenomenon known as GC-biased gene conversion (GCbGC), which can drive such mutations to higher frequencies, especially in larger populations^26^. Because most hominin recombination occurs in hotspots positioned by PRDM9, a rapidly-evolving zinc-finger protein that binds specific DNA motifs to initiate recombination^27,28^, GCbGC^26^ occurs nearby hotspots that have been active in the recent past. Therefore, two groups with different ancestral hotspots, or different ancestral sizes, can show different GCbGC patterns at hotspots, genome-wide. Similarly, mutations within target motifs disrupting the binding of PRDM9 are also overtransmitted, a phenomenon known as “hotspot death” and generating analogous signatures in humans and other species^27,29^.

We apply GhostBuster to both modern human and archaic hominin samples, uncovering three significant sets of events. First, we characterise Holocene-era back-to-Africa migrations that introduced detectable levels of Eurasian ancestry into all sub-Saharan African populations we analysed. Several recent studies have identified such gene flow impacting particular African populations^12,30–32^. Our analyses reveal that these migrations occurred between 3-14,000 years ago at multiple times and involved at least three different source populations. Moreover, they fully explain the presence of small amounts of Neanderthal ancestry in modern Africans, and introgressing segments also carry a clear signature of elevated TCC to TTC substitutions.

Second, we identify an admixture occurring around 50,000 years ago in ancestors of African populations between two groups that diverged at least 300,000 years ago, one of which is closely related to the population that gave rise to present-day populations outside of Africa. We hypothesise that this “out-of-Africa”-related population subsequently expanded both within and outside Africa, producing a gradient of admixture proportions across present-day African groups. Using GCbGC mutational signatures, we demonstrate that the out-of-Africa related population had a smaller population size, with differences in PRDM9-driven hotspot activity between the two groups. Our analysis also reveals differing levels of polygenic score (PGS) portability between the two ancient population backgrounds, for genome-wide association studies (GWAS) conducted in European cohorts.

Finally, we characterise an ancient event involving two populations that likely diverged more than 1M years ago. Remarkably, we find, using mutational signatures, that one population (Population HC) evolves the PRDM9-C hotspot motif and entirely lacks activity at PRDM9-A hotspots, while for the other (larger) population (Population HA), this pattern reverses. We demonstrate that ancestors of Neanderthals and Denisovans diverged from the HC hominin lineage; however, Neanderthals subsequently received gene-flow from the HA hominin lineage, importing PRDM9-A associated signatures, which are not observed in Denisovans. This event corresponds to the previously proposed “human-to-Neanderthal” gene-flow. The HA and HC populations also admixed in a separate event, approximately 300,000 years ago, to form ancestors of modern humans. Overall, our results provide a detailed picture of early human evolutionary history and its associated genomic signatures, revealing how deep population structure and ancient gene flow have shaped patterns of genomic diversity in modern and archaic humans.

### Overview of the GhostBuster algorithm

GhostBuster fits ancestral admixing components to target individuals using an Expectation-Maximisation algorithm. The method uses observed coalescences between a target individual and reference populations as input and iterates between two steps until convergence (Figure 1a): First, conditional on the current local ancestry assignment, it updates each ancestry component’s coalescence rates to reference populations. Second, it updates local ancestry by computing a posterior probability that observed coalescence events were drawn from a given ancestry component, conditional on that component’s coalescence rates (see Methods and Supplementary Note for full details of the theoretical derivations and algorithmic details). We implement a number of strategies to identify the number of components to be fit, such as cross-validating on held-out chromosomes (Methods). GhostBuster can focus on a specific time period by only considering coalescence events occurring within a prespecified window. We can run GhostBuster by fixing ancestry proportions to constrain the model to fit rare ancestries, and we can also run the method in ‘supervised’ mode by fixing coalescence rates for each component. We use supervised GhostBuster if admixing sources are known and included in the genealogy. We infer genealogies using Relate^15^, given its scalability and accuracy, but GhostBuster is, in principle, compatible with any genealogy inference method.

**Figure 1:**
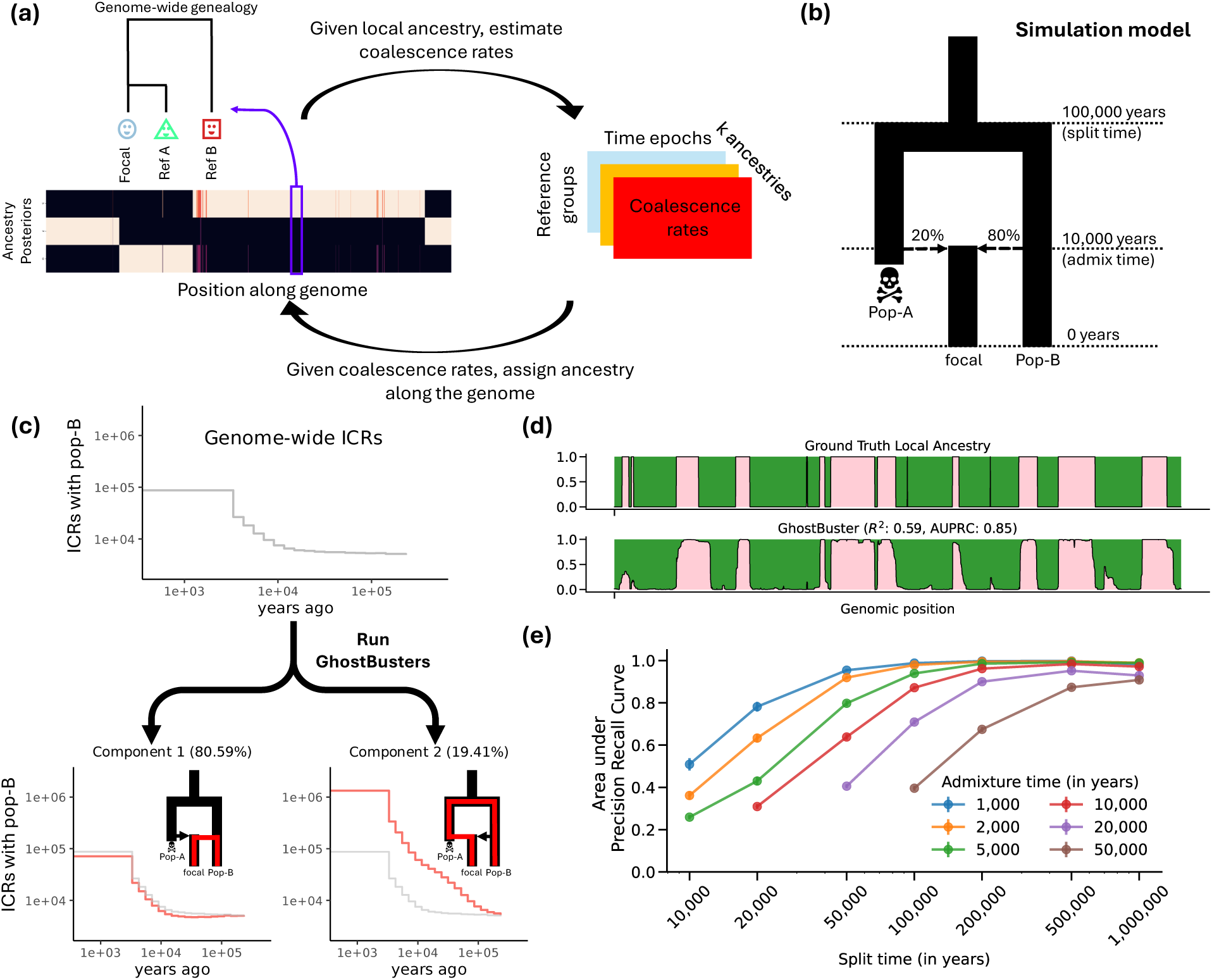
Overview of the GhostBuster framework and validation using simulated data. (a) Schematic overview of GhostBuster. Given local ancestry along the genome, the method estimates time-stratified coalescence rates between the focal genome and reference groups, and reciprocally infers local ancestry from these rates in an iterative framework. (b) Simulation model used for benchmarking: an admixed population (focal) formed 10,000 years ago from two ancestral groups (Pop-A and Pop-B) that diverged 100,000 years ago, with 20% ancestry from Pop-A. (c) Genome-wide inferred coalescence rates (ICRs) between the focal individual and Pop-B reveal two major components corresponding to the two ancestral sources. GhostBuster separates these, quantifying their ancestry contributions. (d) Example comparison between ground-truth and inferred local ancestry along one simulated chromosome. GhostBuster accurately recovers ancestry segments (R² = 0.59, AUPRC = 0.85). (e) Performance across simulations with varying split and admixture times, quantified by area under the precision–recall curve (AUPRC). Error bars: mean ± 95% confidence intervals from 100 jack-knife replicates.

After applying GhostBuster, we can feed local ancestry posteriors into several downstream analyses, including admixture dating via co-ancestry curves^33^, evaluating expected accuracy^34,35^, or probing differential mutation and recombination patterns.

### Simulations and recent admixtures

To validate GhostBuster’s ability to detect admixture from unsampled source populations, we simulate a focal (admixed) population formed as a two-way admixture between populations A (contributing 20%) and B (80%) occurring 10,000 years ago. The two source populations split from a common ancestor approximately 100,000 years ago. Crucially, we do not sample population A, so it represents a ghost population, and only population B is provided as a reference (Figure 1b). Few events of this type have been identified within these time ranges to date^36^, reflecting the limited power of existing methods and scarcity of aDNA from such periods.

For this scenario, GhostBuster reliably infers two ancestry components (Figure 1c) and accurately infers local ancestry (R² = 0.59 vs. true ancestry; Figure 1d). Component 1 captures genomic regions that coalesce more recently with population B, while component 2 captures regions that coalesce more distantly, indicative of ancestry from an unsampled source. In 72 additional simulation settings varying the admixture and divergence time between populations A and B, and more complex scenarios, GhostBuster maintains high accuracy (Figure 1e). When samples from population A are available (Supplementary Figure 1), performance improves further, as expected. Finally, for a scenario mimicking a recent four-way admixture event (Supplementary Figures 2-3) or Denisovan admixture into Papuan people (Supplementary Figures 4-5), GhostBuster performs favourably compared to a leading HMM-based method^37^. GhostBuster is also able to flag scenarios without any simulated admixture event (Supplementary Figure 6).

We also validated GhostBuster’s performance using well-documented cases of recent admixture using real data. We compared GhostBuster’s local ancestry posteriors to those inferred by Mosaic^35^ in three HGDP populations^38^: Hazara, Bedouin, and Maya. In all three cases, GhostBuster successfully recovers the expected admixture signal, producing local ancestry posteriors strongly concordant with Mosaic (Supplementary Figures 7-9).

### Africa-wide Eurasian back-migration events spanning at least 10,000 years

To investigate admixture history within Africa, we assembled a dataset comprising 17 modern sub-Saharan African groups^38–40^, three archaic Neanderthal^41–43^ and one Denisovan hominins^44^, and five high-coverage ancient genomes: a ∼7,900-year-old from Shum Laka, Cameroon^45^, a ∼4,500-year-old from Mota Cave, Ethiopia^32,46^; a ∼1,900-year-old from Ballito Bay, South Africa and two 400-year-old from Eland Cave, and Newcastle, South Africa^46^ (Supplementary Figure 10, Supplementary Table 1).

Before investigating deeper events, we first searched for Eurasian-like ancestry across sub-Saharan Africa. We collapsed multiple Eurasian populations into a single super-population and used GhostBuster in ‘supervised’ mode, fixing coalescence rates for each ancestry component to their inferred combined African and Eurasian genome-wide values, derived from the genealogies (Methods). GhostBuster-estimated proportions range from 1.8% in South African San up to 23.4% in Somali and 15.2% in Masai from East Africa (Figure 2a,b, Supplementary Table 2). In ancients, we identify 1.6% and 2.4% Eurasian-like ancestry in the 400-year old Newcastle and Eland cave individuals from South Africa, 1.1% in the 7,900-year-old Shum Laka individual from Cameroon, 0.7% in the 4,500-year-old Mota individual from Ethiopia and just 0.5% in the 1,900-year-old Ballito Bay individual from South Africa.

**Figure 2:**
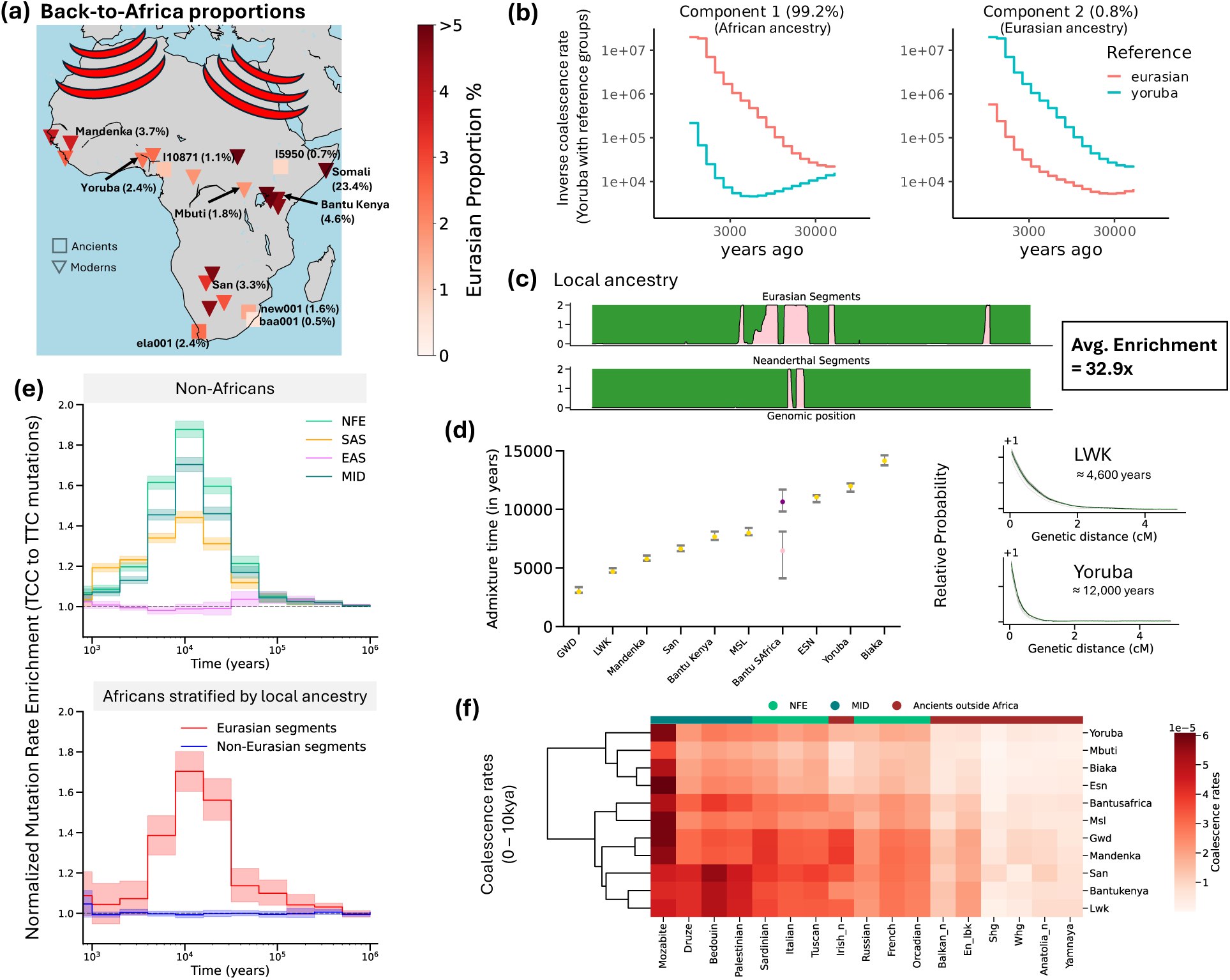
GhostBuster reveals Eurasian backflow into Africa and its genomic consequences. (a) Map of African populations analysed, showing inferred proportions of Eurasian-like ancestry. (b) Example decomposition of genome-wide inverse coalescence rates (ICRs) for a Yoruba individual, revealing two components corresponding to African-like and Eurasian-like ancestries. (c) Representative local ancestry reconstruction highlighting 32.9x average Neanderthal-enrichment in Eurasian-like ancestry segments. (d) Estimated admixture times across African populations inferred from co-ancestry curves (left), with examples for LWK and Yoruba shown on the right. Error bars: mean ± 95% confidence intervals from 100 jack-knife replicates. (e) Normalised mutation rate enrichment of TCC to TTC trinucleotide substitutions versus mutation age for non-African populations (top) and African genomes stratified by local ancestry (bottom). Shaded regions: 95% confidence intervals, using 1,000 bootstrap replicates. (f) Heatmap of coalescence rates between African and ancient/modern reference populations over the past 10,000 years. The dendrogram is inferred using the UPGMA algorithm with ICRs with non-African groups as the distance matrix. Non-Finnish Europeans (NFE), East Asians (EAS), South Asians (SAS), Middle Eastern (MID).

Previous studies have identified traces of Neanderthal ancestry in modern African genomes, likely introduced through back-migration from Eurasia^12,38,47,48^. We again used supervised GhostBuster (Methods), this time to identify Neanderthal ancestry tracts, observing 0.009% (Mbuti) to 0.049% (Bantu Kenya) Neanderthal ancestry among modern African groups. Though inferred separately, Neanderthal and Eurasian ancestry proportions show strong correlation (p = 2.4 × 10⁻²⁴; slope = 1.05%, intercept ≈ 0; Extended Data Figure 1), suggesting that all Neanderthal ancestry in Africans is explained by Eurasian ancestry. Indeed, Eurasian-like ancestry segments within African genomes are 32.9-fold enriched for Neanderthal ancestry compared to non-Eurasian-like segments (Figure 2c). In ancient Africans, Neanderthal ancestry occurs in the ∼400-year-old Newcastle (0.019%), Eland Cave (0.037%), and ∼1,900-year-old Ballito Bay (0.004%) individuals but is undetectable in Mota and Shum Laka. We therefore hypothesise that Eurasian gene flow had not yet impacted Mota and Shum Laka and that their weak Eurasian-like signal likely reflects a deeper admixture layer lacking Neanderthal input (see next section). To date Eurasian back-migration, we constructed co-ancestry curves only from Eurasian segments containing at least one Neanderthal tract (Methods), yielding dates from ∼3,000 to ∼14,000 years ago (Figure 2d). A single-pulse model fits well to all populations except Bantu South Africans, who show evidence of two back-migration episodes. The inferred dates are notably younger for East African and coastal West African populations such as Gambia, Luhya, and Mandenka (∼3,000–6,000 years), and older for inland West African and rainforest hunter-gatherer groups, including Esan, Yoruba, and Biaka (∼10,000–14,000 years), consistent with prolonged back migration impacting coastal regions.

In search of distinct evidence to support our genealogical inferences, we investigated whether GhostBuster-inferred Eurasian-like segments exhibited distinctive mutational signatures. We examined TCC to TTC trinucleotide changes as these are known to be enriched in West Eurasian genomes, observing a significant enrichment within Eurasian-like segments (Figure 2e, Supplementary Figure 11). These were present at varying magnitudes, suggesting differences in the timing and/or sources of Eurasian backflow across Africa (Extended Data Figure 2) for all HGDP + 1,000 GP dataset modern African groups. Together with Neanderthal ancestry, this offers strong confirmation that inferred tracts reflect genuine admixture events and that they possess a predominantly West Eurasian origin. As a final signature, we observe weaker GCbGC in Eurasian versus non-Eurasian segments, expected due to a bottlenecked history^49^ (Extended Data Figure 2).

To uncover the uncharacterised potential sources and nature of the admixture, we inferred coalescence rates of inferred Eurasian-like ancestry segments across many modern and ancient human populations (Methods). Ancient Eurasian segments in West African populations showed faster coalescence with North African groups, including Mozabites, and appear basal to available ancient Eurasian samples. Contrastingly, more recent segments in East Africans and San coalesce more rapidly with Middle Eastern groups such as the Bedouin, and ancient Neolithic farmer-like groups (Figure 2f). Mandenka and GWD, West African coastal populations possessing a more recent admixture date, show a third pattern: recent coalescence with western Neolithic farmer populations (Figure 2f). Together, these patterns point to at least three temporally and geographically distinct back-migration events that differentially impacted African regions (Discussion).

### Expansion of human groups within and out of Africa

We next decomposed African populations in our dataset using GhostBuster in ‘unsupervised’ mode (here, GhostBuster learns coalescence rates from the data) for evidence of deeper admixture (Methods). For every African group analysed, GhostBuster infers that two ancestral components provide a better fit to the coalescent history than a single component (Supplementary Figure 12). In all African populations, the minor component (Figure 3a, Supplementary Table 2; ∼6% to ∼33%, lowest in San and rainforest hunter-gatherer groups and highest in East Africa) has closer affinity to Eurasia, while the major component has closer affinity to Africa. Unlike for back-migration, Ancient individuals show similar inferred proportions to the corresponding modern populations: the ∼1,900-year-old Ballito Bay individual exhibited the lowest proportion (∼6%), whereas the ∼4,500-year-old Mota individual displays one of the highest (∼30%).

**Figure 3:**
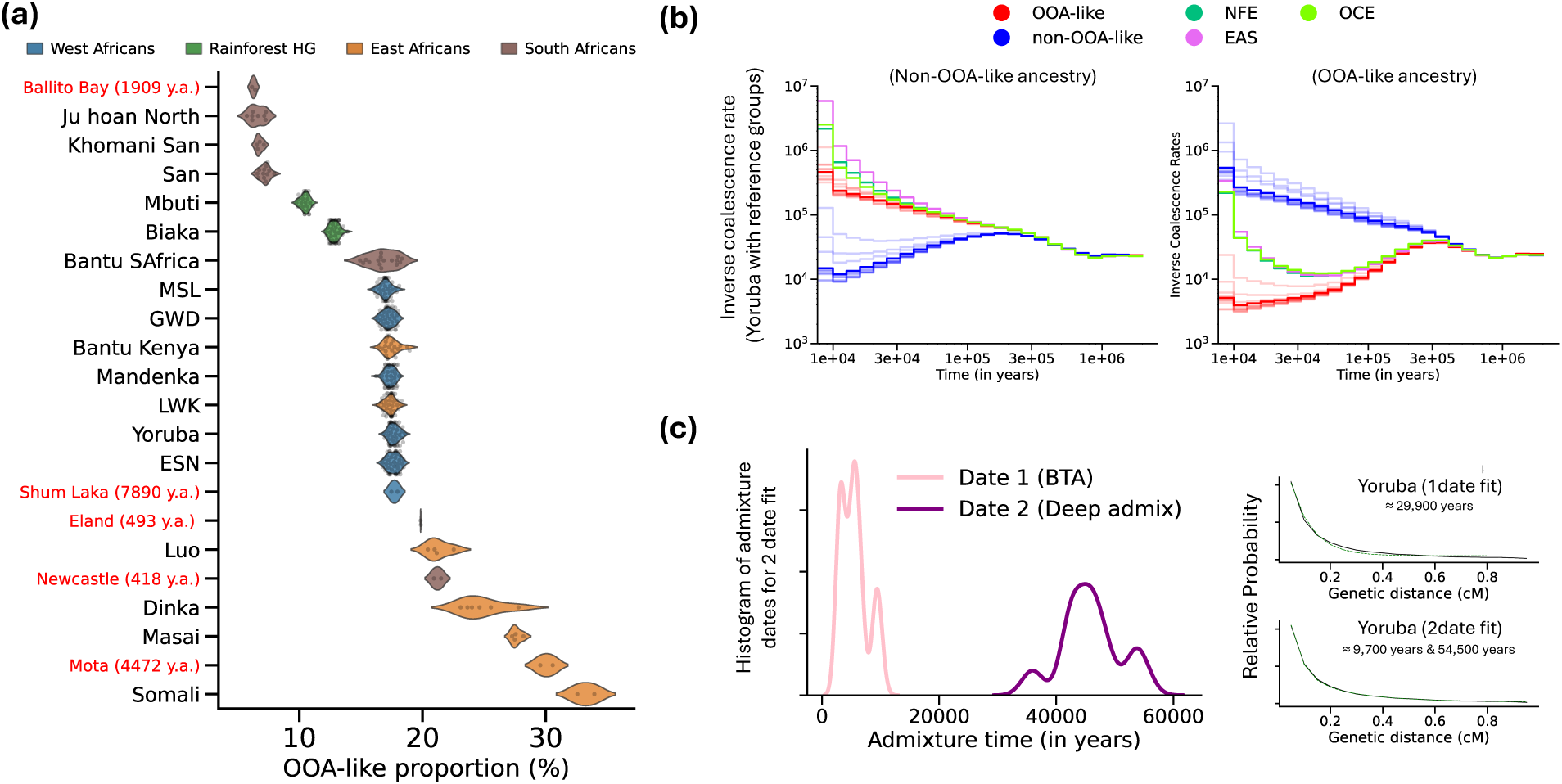
GhostBuster identifies a deep admixture event in Africa. (a) Estimated proportions of out-of-Africa (OOA)-like ancestry across ancient and present-day African populations. Each point represents an individual, colored by regional grouping: West Africans (blue), Rainforest hunter-gatherers (green), East Africans (orange), and South Africans (brown). Ancient individuals are labelled in red with sample age (years ago). Dots represent OOA-like proportions for specific individuals. Back-to-Africa gene flow was removed to calculate OOA-like proportions. (b) Inverse coalescence rate profiles between ancestry components in Yoruba. Shaded bands denote different population groups within Africa. (c) Genome-wide distribution of admixture dates for all modern Africans under a two-date model. Right: co-ancestry curve fits for single- and two-date models for Yoruba. Non-Finnish Europeans (NFE), East Asians (EAS), Oceanians (OCE).

Using the GhostBuster fit for all African individuals, we estimated coalescence rates between the two ancestral components across the inferred local ancestries of each African individual (Methods). The resulting coalescence profiles indicate that GhostBuster fitted very similar components in all individuals (Figure 3b). The minority component coalesces rapidly with out-of-Africa (OOA)-related populations (OOA-like component) and splits around 300 KYA from the majority component that is more distinctively African (non-OOA-like component). Coalescence rates between present-day non-African populations diverge from the OOA-like component around ∼50 KYA, coinciding approximately with estimates of out-of-Africa migrations^2^. Co-ancestry curve–based admixture dating supported a two-date model across all African groups (Extended Data Figure 3). The younger dates (∼3,000–11,000 years) overlap the dates of the previously analysed more recent back-to-Africa migrations, which were (re)detected here because of the overlapping coalescence rate profiles between the two runs. The older date estimates range from ∼35,000 to 57,000 years ago, but may be underestimated if recombination map uncertainty is taken into account, and correspond to ancient population structure within Africa (Figure 3c). We further confirmed that the inferred mixture event is generally robust to running the method without a Eurasian reference population in GhostBuster (Supplementary Figure 13). High expected R^2^ values^35^(Supplementary Figure 14) evidence confident local ancestry calls, and we observe little to no correlation with the local recombination rate, B-statistic^50^ measuring background selection effects (Supplementary Figure 15) and local ancestry estimates from ^3^ (Supplementary Figure 16). To avoid ambiguity, we use “Eurasian ancestry” in African populations to refer specifically to the Holocene-era back-migration flow, as described in the previous section, whereas “OOA-like ancestry” denotes the ancestral component associated with this deep admixture event.

To corroborate this mixture event, we again examined patterns of mutation types within each ancestry component, information not used when fitting GhostBuster components. We observed differences in recombination-associated mutational patterns between the two ancestral components. Two major groups of PRDM9 alleles occur in humans today^51^, and lead to mostly non-overlapping hotspot landscapes^51,52,53^. The A-type alleles are ubiquitous across populations and predominant outside Africa, while the C-type alleles are most frequent in African populations. In our dataset, the PRDM9-C allele tagged by SNP rs6889665^53^ has a frequency of 33.7% in Africans, 8.4% in East Asians, 7.4% in South Asians, and 6.3% in non-Finnish Europeans. To assess historical recombination activity, we analysed GCbGC around these hotspot regions within OOA-like and non-OOA-like ancestry segments (Methods). We observe higher levels of weak-to-strong mutations at PRDM9-A hotspots in the OOA-like component (p = 0.009; Figure 4a), whereas PRDM9-C hotspots showed more weak-to-strong mutations in the non-OOA-like component (p = 0.033; Figure 4a). These contrasting patterns agree with inferred ancestry around the PRDM9 gene, where we estimate that PRDM9-C has a markedly higher frequency in the non-OOA-like group (44.1%) compared to only 2.7% in the OOA-like group (Fisher’s exact p = 2.5 × 10⁻^15^, Extended Data Figure 4).

**Figure 4:**
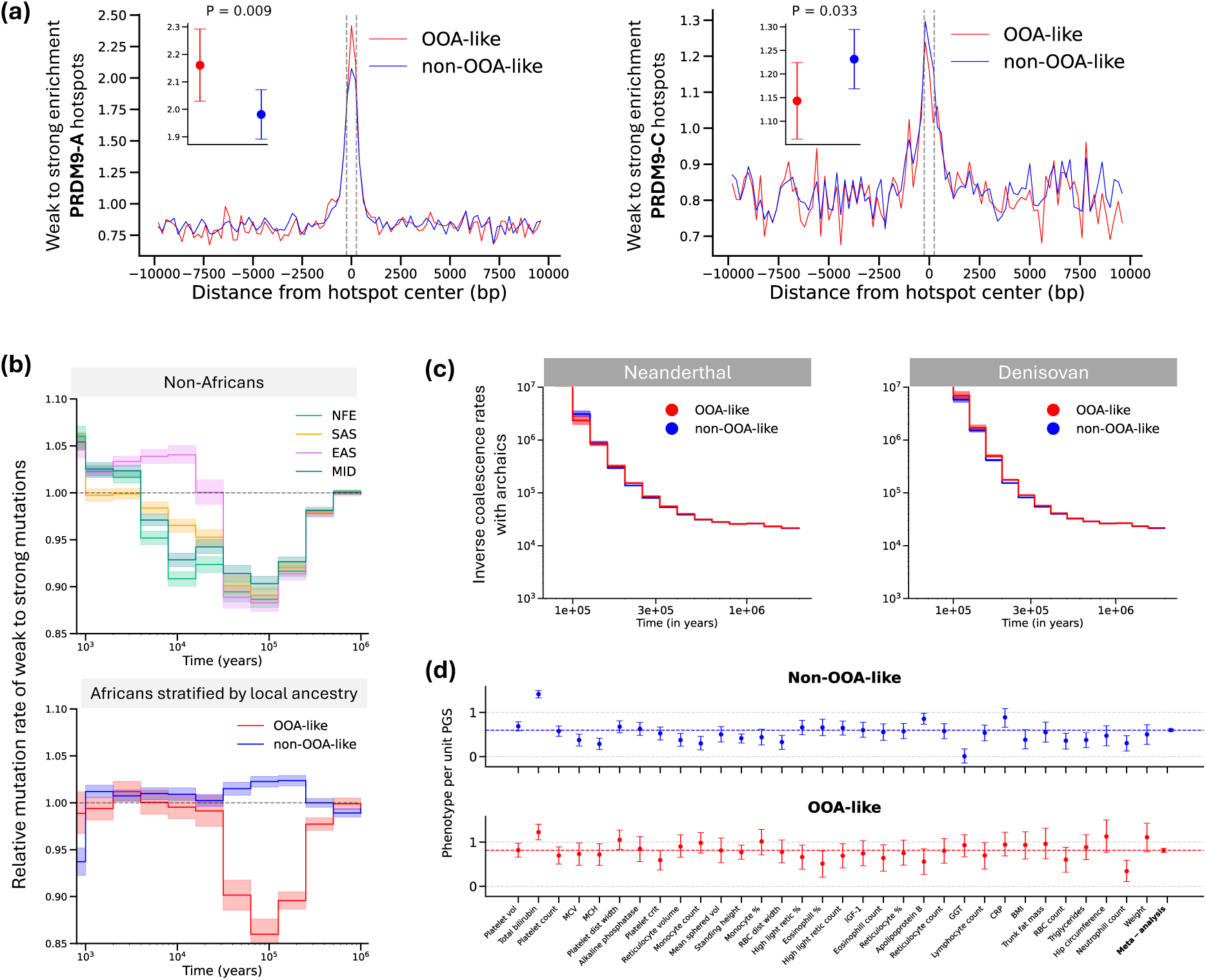
Characterising the deep admixture event in Africa. (a) Weak to strong enrichment at PRDM9 hotspots showing more pronounced GCbGC in OOA-like (red) ancestry around PRDM9-A (left) and non-OOA-like (blue) ancestry around PRDM9-C (right) hotspots. Insets: total enrichment levels ± 95% confidence intervals focusing on ± 250bp around the hotspot centre; Confidence intervals and statistical significance use 1,000 bootstrap replicates (P = 0.009, P = 0.033, respectively). (b) Normalised mutation rate enrichment of weak-to-strong substitutions through time. Top: non-African populations (NFE, SAS, EAS, MID). Bottom: African genomes stratified by local ancestry. Shaded ribbons: 95% confidence intervals from 1,000 bootstrap replicates. (c) Inverse coalescence rates between ancestry components and archaic genomes (Chagyrskaya Neanderthal and Denisovan). (d) Polygenic score portability difference between ancestry components. Each point represents the regression coefficient (phenotype per unit PGS) ± 95% confidence interval from 1,000 bootstrap replicates for non-OOA-like (top) and OOA-like (bottom) ancestry components. The dashed line represents the mean regression coefficient across the traits analysed. Non-Finnish Europeans (NFE), East Asians (EAS), South Asians (SAS), Middle Eastern (MID).

The OOA-like component also has significantly reduced overall weak to strong enrichment, consistent with a smaller effective population size reducing GCbGC strength, compared to the non-OOA-like component. The magnitude and timing of this reduction closely matches non-African populations, suggesting that the OOA-like group in Africa has experienced a similar population bottleneck (Figure 4b, Supplementary Figure 17). The level of enrichment is consistent across all African populations analysed, supporting a single shared ancestral population underlying this component (Extended Data Figure 5).

To assess whether either ancestral component showed differential affinity to archaic humans outside Africa, we computed D-statistics of the form D(OOA-like, non-OOA-like; Neanderthal/Denisovan, Chimpanzee) (Methods). No D-statistic is significantly non-zero, indicating no detectable differences in archaic affinity between the OOA-like and non-OOA-like components. Coalescence rate profiles with Neanderthals and Denisovans also suggest very similar affinities of each with the two ancestry components (Figure 4c), with Denisovans slightly closer to the non-OOA-like group.

Finally, to evaluate the functional consequences of this deep population structure, we assessed the portability of PGS (polygenic scores) within Africa. PGS derived from European genome-wide association studies are known to perform poorly in African populations^54,55^. We constructed PGS for 32 blood-related and anthropometric traits using Quickdraws^56^. We used ANCHOR^57^ to decompose these scores, for 4,557 African-ancestry UK Biobank individuals, into contributions from their OOA-like and non-OOA-like ancestries (Methods, Supplementary Figure 18). The OOA-like component was more predictive than the non-OOA-like component for 31 of 32 traits (Figure 4d; paired t-test, p = 5.3 × 10⁻⁶, mean predictive power ∼70% higher). Therefore, genomic segments of OOA-like ancestry retain substantial cross-population PGS transferability, with the reduced predictive performance observed in African individuals predominantly driven by non-OOA-like ancestry.

### Gene-flow into Neanderthals implicates million-year deep structure in hominins

To examine deeper hominin evolution, we focused on the coalescent history of archaic hominins using (African) humans, Neanderthals and Denisovans. Previous studies, using uniparental markers and autosomal ancestry^13,58–60^, suggested a model whereby Neanderthals received limited gene flow from the modern human lineage after it split from a population ancestral to both Neanderthals and Denisovans. Denisovans were hypothesised to also possess a small proportion of older, “super-archaic” ancestry^14,43^. In particular, the D-statistic D(Neanderthal, Denisovan; Human, Chimpanzee) is significantly positive across a range of human allele frequencies (Extended Data Figure 6)^43^. Instead of stratifying by Relate-inferred mutational age^61^, we observe a strong D-statistic peak around 300,000-500,000 years ago (Figure 5a; Extended Data Figure 6), consistent with gene-flow between an ancestral human group and Neanderthals more recently than this interval.

**Figure 5:**
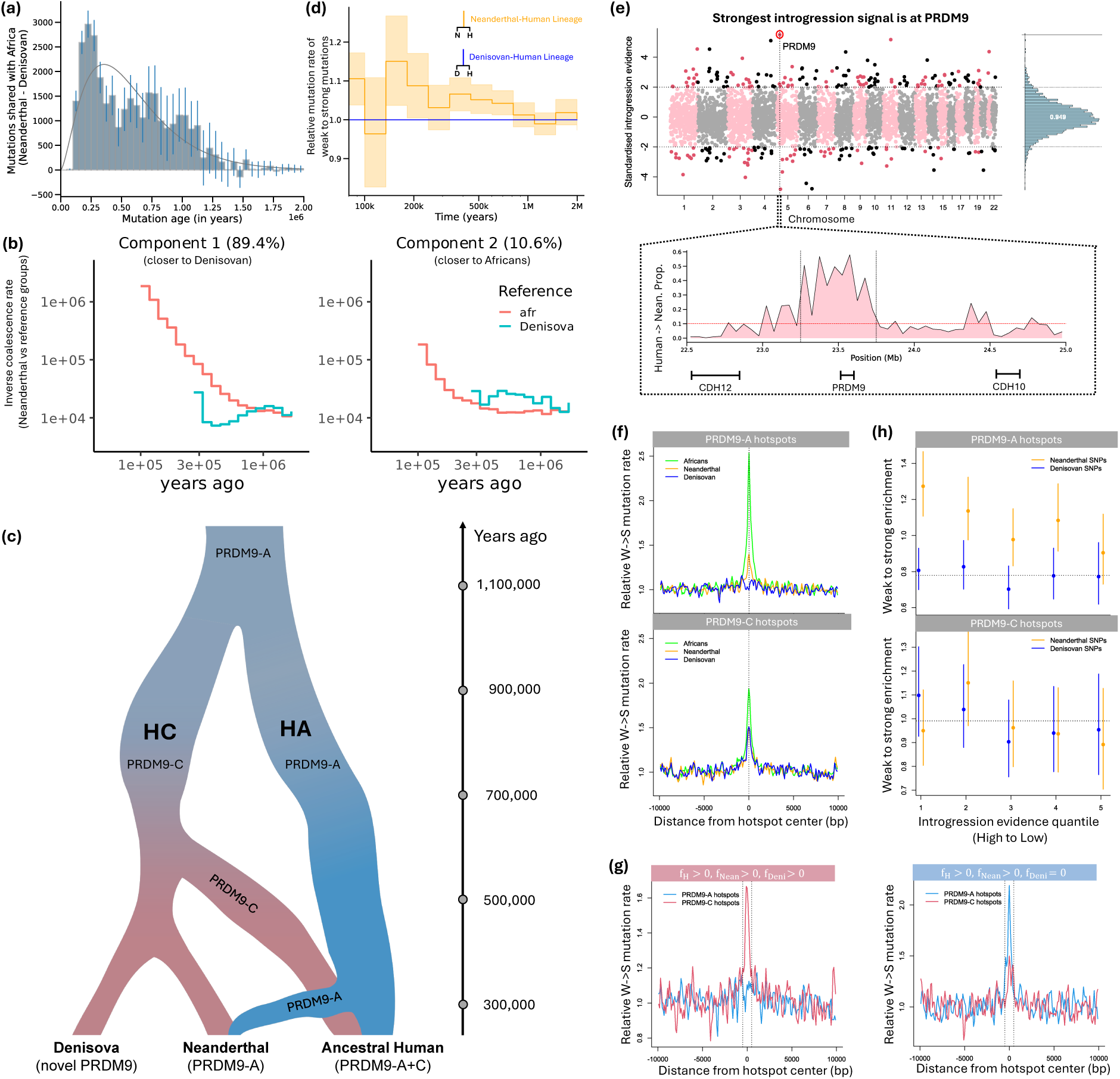
Human to Neanderthal gene-flow and deeper structure in Humans. (a) Number of excess variants shared between African and Chagyrskaya Neanderthal vs. Denisovan genomes, stratified by lower mutational age. The smooth curve indicates the fitted density. Error bars: 95% confidence intervals using 1,000 bootstrap replicates. (b) GhostBuster-based decomposition of genome-wide inverse coalescence rates (ICRs) for the Chagyrskaya Neanderthal, revealing two components corresponding to the HC (left) and HA (right) ancestries. (c) Proposed model of deep human history showing the divergence and persistence of different PRDM9 lineages across Denisovans, Neanderthals, and ancestral humans over the past >1 million years. HA and HC denote population groups carrying PRDM9-A-like and PRDM9-C-like PRDM9 alleles, respectively. (d) Relative weak-to-strong mutation rates through time for the Neanderthal–human lineage, relative to the Denisovan–human lineage. Shaded regions: 95% confidence intervals using 1,000 bootstrap replicates. (e) Genome-wide scan of standardised introgression evidence identifies the strongest signal at PRDM9. The inset shows localised enrichment of GhostBuster-inferred HA ancestry proportions nearby PRDM9 on chromosome 5. Dotted vertical lines: 500kb region boundaries, dotted horizontal lines: mean local ancestry posterior genome-wide. (f) Relative weak to strong mutation rates in the 20kb surroundings of PRDM9-A and PRDM9-C recombination hotspots in Africans, Neanderthals, and Denisovans. (g) Relative weak to strong mutation rates at PRDM9-A and PRDM9-C hotspots partitioned by observed derived-allele status (0 or >0 copies) in humans, Neanderthals, and Denisovans. (h) Weak to strong enrichment at PRDM9-A and PRDM9-C hotspot centres (± 250bp), stratified by introgression evidence quantiles. Error bars: 95% exact binomial confidence intervals.

Applying GhostBuster to decompose Neanderthal ancestry into two components identifies one component closer to Denisovans, and a minority component instead coalescing more quickly with modern humans, particularly >300KYA, while showing no clear bottleneck and so suggesting a maintained large size (Figure 5b). To fit this event, which is of extreme antiquity (at least >150KYA^13^), we fixed ancestry proportions, but did not otherwise constrain the model fit (Methods). We analysed patterns of shared mutations across age categories to obtain a range of 8.5-12.3% admixture (Extended Data Figure 7; Methods). Inference results across this range were similar (Supplementary Figure 19), and we present results using an admixture fraction of 10%.

Intriguingly, the coalescence rate curves of the two inferred components converge only in the deep past: >1 MYA (Figure 5b). In particular, the minor “human-like” ancestry remains strongly distinct from the lineage ancestral to Denisovans, based on the inferred coalescence rates with modern humans. This is unexpected based on prior divergence time estimates of 500 - 700KYA^2^ between humans and the population ancestral to Neanderthals and Denisovans, which would be expected to represent an approximate upper bound on coalescence rate differences, and so suggests unexpectedly deep divergence between the admixing groups. The major ancestry in Neanderthals, which we label population “HC”, coalesces recently with Denisovans and shows similar rates of coalescence with both Denisovan and human lineages >800KYA (Figure 5b), so might be considered as more recently ancestral to all three groups. The minor population “HA” coalesces recently (by ∼300KYA) with modern human lineages and admixes into Neanderthals, but shows separation from the Denisovan lineage until >1MYA. Because both groups therefore coalesce much more recently than this time with modern humans, if correct, this inference implies the existence of two distinct, ancient groups at least partially separated >1MYA and with both contributing ancestry to modern humans (Figure 5c).

Given the age (and resulting subtlety) of this event and signal, we next asked whether, similar to our previous events, we could detect distinct “signatures”, i.e. mutational patterns, specific to these suggested groups, to exclude artefactual explanations and corroborate its validity. We start by quantifying genome-wide weak to strong enrichment along two lineage-specific sets of mutations: those shared between Neanderthals and modern humans but absent in Denisovans (N+H), and those shared between Denisovans and modern humans but absent in Neanderthals (D+H). Under a simple split model where Neanderthals lack additional recent hominin genetic contributions relative to Denisovans, these should be symmetric, but in fact, we observe that N+H lineages show significantly greater weak to strong enrichment than D+H lineages (Figure 5d). This confirms that a hominin group tagged by N+H lineages (consistent with HA) has stronger GCbGC, indicating a larger effective population size, than D+H lineages (consistent with HC), in agreement with our GhostBuster prediction.

### Deep hominin structure evolved different PRDM9 alleles and signatures

Our GhostBuster results predict several properties of regions introgressed in Neanderthals: high GhostBuster-inferred local ancestry (corresponding to the minor component), recent coalescence times between humans and Neanderthals but deep coalescence between Neanderthals and Denisovans, and sharing of mutations between humans and Neanderthals but not Denisovans. We leveraged these signals to identify candidate outlying large introgressed regions within the Neanderthal genome, predicted to arise if a selective “sweep” drove a favourable introgressed variant, and the surrounding haplotype background, to fixation in Neanderthal (Supplementary Note). The strongest such signal, indicating an approximately 500kb introgressed region, contained a single gene, PRDM9 (Figure 5e; Supplementary Table 3a; empirical p=0.0002), with an introgression signal across the three high-coverage Neanderthals indicating a rapid spread of a potentially distinct PRDM9 allele into the Neanderthal population. Notably, the mean Neanderthal-Denisovan inferred coalescence time across this region is very deep: 1.403MYA, supporting a deep split model. This suggests the introgressing HA group and the HC ancestry might differ in their recombination landscapes. To test this, we again examined GCbGC mutational signatures around individual A-type and C-type recombination hotspots, and also the separate effect of “hotspot death”, acting to disrupt the PRDM9 binding motif itself (Supplementary Note).

Genome-wide, the strength of the GCbGC signal (Figure 5f) indicates that all of humans, Neanderthals, and Denisovans have historical recombination activity at PRDM9-C hotspots, strongest in humans, showing a higher level, and similar and weaker in Neanderthals and Denisovans. At PRDM9-A hotspots, however, only humans and Neanderthals, but not Denisovans, show activity. This pattern could be explained either by strongly oscillating PRDM9 A and C frequencies, or by admixture, if population HC is active for PRDM9-C, while (only) population HA has PRDM9-A activity.

To distinguish these, we partitioned mutations (Methods) based on which hominins carried their derived alleles (e.g. Human-Neanderthal-Denisovan shared, or Human-Neanderthal shared, or Human-only). This revealed (Figure 5g) that the PRDM9-A weak-to-strong enrichment in Neanderthals is almost entirely driven by mutations also shared with modern humans, with only a very weak Neanderthal-only signal, and no signal in other combinations. This suggests this signal indeed arose due to admixture into Neanderthals from a PRDM9-A carrying population (Extended Data Figure 8). In contrast, for PRDM9-C, we observe (Figure 5g) clear weak-to-strong activity for mutations jointly shared by humans, Neanderthal and Denisovan, and weaker signals for both Denisovan-human and Neanderthal-human shared mutations, of similar strength in both archaics, and so potentially explained by mutations arising in the HC population ancestral to humans, Denisovans, and Neanderthals and variably segregating through incomplete lineage sorting. Mutations absent in humans show no signal in either Denisovans, Neanderthals or both, suggesting populations ancestral to archaics but not humans were too short-lived and/or small to generate GCbGC, or carried different PRDM9 alleles (Extended Data Figure 8).

We additionally stratified hotspots based on whether Neanderthals showed evidence of recently shared ancestry with humans locally and examined their GCbGC signals. As this examines recombination hotspots, we did not use GhostBuster local ancestry calls (which we expect to be very unreliable within hotspots for this ancient event), but instead stratified hotspots using their distance to the nearest Neanderthal-human young (and so potentially introgressed) shared SNP (Methods). This showed (Figure 5h) a strong and significant (p=5×10^-6^) weak-to-strong drive at PRDM9-A hotspots nearby such mutations, evidencing likely introgression of the hotspot region itself. This signal disappeared (p=0.32) at regions suggesting no introgression, for PRDM9-C hotspots (Figure 5h), and for Denisovan GCbGC across both hotspot types.

To provide an additional test for the event, we further examined signatures of hotspot death (Supplementary Note). Hotspot death, although restricted to recombination motifs, is mechanistically strong relative to GCbGC and drives rapid hotspot turnover in other species^29,62^. Consistent with the GCbGC signal, we find significant hotspot-death signal in high-frequency human-specific mutations for both PRDM9 alleles (OR > 6, p < 10⁻¹⁷), human–Neanderthal shared mutations at PRDM9-A hotspots (OR = 4.5, p < 10⁻¹⁵), and human, Neanderthal, and Denisovan shared mutations at PRDM9-C hotspots (OR = 1.8, p = 0.012). No other categories show significant excess (Extended Data Figure 9a-c; Supplementary Table 3c-d). We additionally tested whether these hotspot death signals are associated with nearby admixture signals in Neanderthals. PRDM9-A hotspot death mutations are significantly enriched on introgressed segments even relative to nearby mutations of similar frequencies (p = 0.021; Extended Data Figure 9d–f; Supplementary Table 3c-d), implying PRDM9-A was active in the introgressing population. In contrast, PRDM9-C hotspot death mutations avoid introgressed segments and are instead enriched on non-introgressed backgrounds (p = 0.035), implying PRDM9-C was not historically active in the introgressing population. Control tests swapping PRDM9-A and PRDM9-C and retesting yielded non-significant p-values (Supplementary Note). Together, these results indicate that introgressed regions in Neanderthals reflect ancestry from a PRDM9-A active lineage. Non-introgressed regions and the Denisovan genome instead derive from a PRDM9-C active lineage, supporting long-standing separation between two deeply diverged populations with distinct recombination landscapes, and inconsistent with a model of sequential allele replacement within a single ancestral lineage.

Overall, our results are fully and naturally explained (Figure 5c, Extended Data Figure 10) by a model involving a distantly split (e.g. >1MYA) population HC (carrying PRDM9-C hotspot activity, which might have evolved in this group) and a population HA (carrying PRDM9-A), as suggested by the GhostBuster results. Such a deep divergence would also allow time for both hotspot death and weak-to-strong mutations to separately accumulate in these populations by the time of population separation and admixture events. We have not been able to identify strongly distinct alternative possibilities that might explain all of our results, given the quite strong constraints our findings impose, although models incorporating, for example, more gradual gene flow among populations might well be possible.

Finally, we asked whether admixture might have directly altered PRDM9 alleles in these archaic hominins. Using a likelihood-based approach to infer PRDM9 genotypes from sequencing reads spanning the zinc finger array (Methods; Supplementary Table 3b), we identified clear differences between Neanderthals and Denisovans^63^. In support of our model (Figure 5c), Neanderthals carry either the canonical PRDM9-A allele (Altai Neanderthal, homozygous) or closely related derived variants containing a signature Q zinc finger motif, including the M10 type and the related L24 type, which is also observed in humans carrying Neanderthal introgression at PRDM9 (Supplementary Note). In contrast, Denisovan PRDM9 alleles are distinct from present-day human genotypes, and at least one allele is more similar to PRDM9-C than to PRDM9-A (Supplementary Table 3b). These results provide further support for distinct PRDM9 histories in Neanderthals and Denisovans and suggest that admixture contributed directly to shaping their recombination landscapes.

## Discussion

The ability of GhostBuster to identify and characterise admixture events in simulations and real data, together with related approaches that leverage coalescence times and mutational ages, illustrates the power of genome-wide genealogies to resolve complex admixture histories. We identify three distinct events shaping deep human ancestry, each involving lineages no longer present in an unadmixed form anywhere, and such events may prove to be very common. Complementing GhostBuster, several analyses involving mutational signatures provide supportive orthogonal evidence relying solely on mutational categories rather than coalescent history to validate inferred admixture events. These leverage processes, including GCbGC, recombination hotspot evolution, and mutation rate pulses, to reveal properties of the contributing populations, including differences in effective population size and recombination landscapes. While not exhaustive, these examples illustrate how integrating genealogical inference with mutation-level analyses can strengthen historical inference. Additional evolving processes, such as transposon activity or other mutational classes, may provide similar opportunities for other populations or species.

Although our results on back-to-Africa migrations partially represent a positive control given previous evidence, we also identify their timings and source populations, indicating that such migrations in fact occurred at multiple times, 3,000-14,000 years ago and involved different migrating groups. The earliest dates, for West and Central African individuals, involve groups related to today’s North African (Mozabite) individuals and might involve migrations during past “warm Sahara” periods^64^, with a later movement contributing much more ancestry into East Africa from groups most closely related to groups including the Bedouin from the Levant region and early farmer-like groups potentially also of Levantine origin. Finally, a migration around ∼3KYA involves groups particularly closely related to western Neolithic farmer-like populations, with a geographically localised impact in West African coastal populations, perhaps suggesting a coastal or maritime route.

Further back in time, GhostBuster identifies the expansion of a bottlenecked group contributing the majority ancestry to the out-of-Africa migration of modern humans and also expanding within Africa, but which did not yet carry Neanderthal ancestry. Our results imply that the population bottleneck seen in today’s populations outside Africa at least partly predates their separation from the wider OOA population. This event may have facilitated the spread of the Y-chromosome CT lineage, which arose around 60-80KYA and is today found as the majority type within and outside of Africa^2,65^. Our analysis is also consistent with *H. sapiens* diversifying across the African continent, as reflected in morphological and material cultural variation in early *H. sapien*s across Africa^66^, suggesting deep population structure over timescales overlapping the ∼300K separation we observe here. In support of this, non-OOA ancestry shows earlier and stronger splits (using coalescent rates) between populations (Figure 3b).

Finally, the oldest events we identify involve hominin groups ancestral to humans, Neanderthals, and Denisovans and implicating an admixed history over hundreds of thousands of years, and remarkably leading to divergent PRDM9-driven recombination landscapes. Our analysis proposes that both humans and Neanderthals arise as similar mixtures of ancestral populations, but in different proportions. These groups show completely different recombination landscapes, which provide very strong evidence for their existence, as well as supporting a long-term split between them. This fact might also explain previous observations^51^ of diversity patterns at PRDM9 in humans: there are diverse alleles, each clearly more similar to either PRDM9-A or PRDM9-C, but few apparent recombinants between these allelic types despite experimental evidence from sperm typing that such alleles can arise. This can be explained by a long history of separate evolution of these alleles within distinct populations.

Our findings relating to humans are broadly in agreement with a previously inferred deep human admixture event^3^, despite very different approaches of (in our case) decomposing the Neanderthal genome and recombination landscapes, versus fitting humans directly. Two additional features suggest these might be somewhat similar events: the fact that the PRDM9-A containing population HA is large, and the >1MY divergence time (vs ∼1.5MY in ^3^). Although it was estimated^3^ that this group contributed only ∼20% to human ancestry, the fact that the PRDM9-A-related drive is today stronger than the PRDM9-C drive, generating more high-frequency and even fixed mutations, might, tentatively, suggest a larger actual contribution. Our results suggest admixture into Neanderthals within the past ∼300KYA. We are not able to date the mixing event in humans, but given PRDM9-A and C drive is observed in the later OOA and non-OOA populations, mixing likely occurred by ∼300KYA when these populations split. Finally, it is intriguing that the clearest signal in the genome for introgression of HA into Neanderthals is a region containing PRDM9, and this resulted in Neanderthals acquiring a recombination landscape similar to that in most modern-day humans. The large size (500kb) of the accompanying introgressed selective sweep-like signal suggests strong selection favouring this allele (∼0.5%^67^), providing the first specific example, to our knowledge, of a contemporary selective event at this locus, despite known rapid PRDM9 and hotspot evolution across many species.

After accounting for hominin admixture into Neanderthals, we observe symmetric relationships of Denisovans and Neanderthals to humans, including at fixed human mutations (Extended Data Figure 6). Our analysis, therefore, does not support substantial additional ancient “super-archaic” admixture into Denisovans, as previously proposed^14,43,58^, although we cannot exclude very small contributions.

Our present work uses only genetic evidence, and paleoanthropological evidence might, in the future, shed light on the identities and natures of these mixing groups. Populations HA and HC were extant in the middle Pleistocene, a time period sometimes referred to as the “muddle in the middle” in hominin evolution. Based on our results, HA ought to represent a long-lived (>1MY, to 300KYA or less) group with a large range to allow it to mix with HC lineages both within Africa (to form modern humans) and in West Eurasia (to form Neanderthals), but which perhaps lacks representation in Asia, given no signal in Denisovans. Evolutionary classification of hominin fossils from this time period, which possess a range of modern and archaic features^68,69^, remains uncertain and controversial. However, among current classifications, we note that fossils attributed to *H. heidelbergensis* are a candidate for HA, as they possess the geographic and temporal properties suggested above.

Because our approach is applicable across a wide range of timescales, from deep population history to more recent admixture events, it offers a general framework for resolving such histories across human populations and other species.

## Data availability

Positions of centromeres, telomeres, and the HLA region: https://genome.ucsc.edu/goldenPath/help/ftp.html; Phased haplotypes for unified HGDP+1,000GP: gs://gcp-public-data--gnomad/resources/hgdp_1kg/phased_haplotypes_v2/; Phased haplotypes for SGDP: https://sharehost.hms.harvard.edu/genetics/reich_lab/sgdp/ (PS2 version); 1,000 GP VCF files: https://ftp.1000genomes.ebi.ac.uk/vol1/ftp/release/20130502/; Input files required for running Relate (including genomic masks and recombination rate maps): https://zenodo.org/records/15801307; Archaic genomes VCFs and mask: https://ftp.eva.mpg.de/neandertal/; Ancient samples: Downloaded from ENA (Supplementary Table 1); Chimpanzee fasta files: https://hgdownload.cse.ucsc.edu/goldenPath/hg19/vsPanTro6/; Posterior effect estimates from Quickdraws are available at https://www.stats.ox.ac.uk/publication-data/sge/ppg/quickdraws/. The individual-level genotype and phenotype data are available to approved researchers through the UK Biobank (http://www.ukbiobank.ac.uk); We will deposit the trees, inferred local ancestries and coalescence rates upon acceptance of the manuscript.

## Code availability

GhostBuster (https://github.com/MyersGroup/GhostBuster); External software used in the current study was obtained from the following URLs: Relate (https://myersgroup.github.io/relate/), Relate_lib (https://github.com/leospeidel/relate_lib) ANCHOR (https://github.com/MyersGroup/ANCHOR), Colate (https://github.com/leospeidel/Colate), Chromopainter (https://github.com/MyersGroup/GhostBuster/tree/master/src/fastchromopainter), Mosaic (maths.ucd.ie/∼mst/MOSAIC/), HMMIX (https://github.com/LauritsSkov/Introgression-detection), msprime (https://tskit.dev/software/msprime.html), SHAPEIT4 (https://odelaneau.github.io/shapeit4/), Bcftools (https://samtools.github.io/bcftools/bcftools.html), CoalRate (https://github.com/leospeidel/CoalRate/).

## Acknowledgements

We thank John Novembre, Geoff Nicholls, Robert Davies, Garrett Hellenthal, Daniel Lawson, and Daniel Falush for helpful discussion and suggestions. This work was conducted using the UK Biobank resource (application no. 43206). We thank the participants of the UK Biobank project. This work was supported by the Clarendon Scholarship (to H.L.); Wellcome Trust (grant no. 108861/Z/15/Z to H.L. and grant no. 212284/Z/18/Z awarded to S.R.M); Medical Sciences Doctoral Training Centre Studentship (to H.L.). L.S. acknowledges support through JSPS KAKENHI grant 24K23946. Computation used the Oxford Biomedical Research Computing facility, a joint development between the Wellcome Centre for Human Genetics and the Big Data Institute, supported by Health Data Research UK and the National Institute for Health and Care Research (NIHR) Oxford Biomedical Research Centre. The views expressed are those of the author(s) and not necessarily those of the National Health Service, the NIHR or the Department of Health. For the purpose of open access, the authors have applied a Creative Commons Attribution (CC BY) licence to any Author Accepted Manuscript (AAM) version arising from this submission.

## Author contributions

H.L., L.S. and S.M. developed the method. L.S. and S.M. supervised the project. H.L., L.S. and S.M. analysed the data and wrote the manuscript. H.L., L.S., S.M., P.F.P., and A.G.H. interpreted the results and edited the manuscript.

## Methods

### GhostBuster implementation

GhostBuster takes the sample IDs of target genomes, a genealogy in tree sequence format, a recombination map, and a file storing population assignment of genomes as input. Additional parameters will be described below.

### GhostBuster modes

GhostBuster models observed coalescence events between target and reference genomes as drawn from a mixture of components. Each component is characterised by its own coalescence rates to each reference group, which GhostBuster learns from the data, while simultaneously inferring admixture proportions, local ancestry, and admixture dates. GhostBuster can be run in two different modes.

First, when the contributing source groups are known and included in the reference panel, the model can be run in ‘supervised’ mode. In this setting, coalescence rates are fixed to the genome-wide estimates between each source population and other reference populations. Only the global ancestry proportions are updated (in the M-step, Supplementary Note), and GhostBuster outputs local ancestry as usual (inferred in the E-step, Supplementary Note). This supervised approach enables targeted fitting of a sample’s history, reducing the risk of fitting unrelated (e.g., potentially older) events. Alternatively, if ancestry proportions are known, these can be fixed with coalescence rates updated in the M-step, and local ancestry updated in the E-step.

Second, GhostBuster can be run in ‘unsupervised’ mode. In this setting, both coalescence rates and proportions are updated in the M-step, and local ancestry is inferred in the E-step.

### GhostBuster parameters

Our implementation further provides several key parameters. First, we can exclude coalescence events younger or older than specific time boundaries. This option allows us to focus the analysis on admixture signals from specific time periods. Second, GhostBuster can be run with or without an HMM that leverages linkage information across local ancestry tracts across the genome. This option is beneficial for fitting recent admixture events. For older events, ancestry linkage is negligible and hence enforcing local continuity in ancestry may introduce slight biases in the inferred coalescence rates. We therefore recommend disabling the HMM to obtain more accurate coalescence rate estimates when running GhostBuster in unsupervised mode for admixture events more than 1000 generations old.

### Choosing the number of components

We specify the number of components GhostBuster will fit. To ensure we determine the appropriate number of components, we conduct several tests to optimise this choice.

First, we assess the model’s log-likelihood on independent, held-out chromosomes. This allows us to distinguish admixture events that replicate consistently across chromosomes from biases that are chromosome-specific and do not generalise well. By default, we select the number of components as follows: we estimate all parameters, such as coalescence rates and proportions, using our EM algorithm on a subset of chromosomes (by default, we use chromosomes 2-5). We then use the fitted parameters to calculate the log-likelihood on a held-out chromosome (by default, we use chromosome 1), always turning off the HMM (treating grid points independently). The optimal number of components is identified by determining where the total log-likelihood across held-out chromosomes is maximised.

Second, we analyse the posterior probabilities of local ancestry by plotting a histogram of the target’s local ancestry posteriors and calculating the expected coefficient of determination (expected *R*^2^), as defined in^34,35^. Expected *R*^2^ estimates the expected correlation between true and inferred local ancestry, assuming the true local ancestry (one-hot encoded) as a random variable following a Bernoulli distribution with the same mean as the inferred local ancestry. The calculation of expected *R*^2^ therefore, does not require true local ancestry. The expected *R*^2^ helps assess the confidence of the cluster assignments, where values greater than 0.7 generally indicate confident local ancestry inference. We use a diploid version of expected *R*^2^, which sums the local ancestry for both haploids; more details and derivation for the diploid version can be found in^35^.

Finally, we use the inferred local ancestry to perform coancestry curve-based dating of the admixture event. Often, we expect admixture LD to persist further along the genome than LD generated by spurious signals. Admixture dates exceeding 2,500 generations (where expected tract lengths fall below 0.05 cM) are challenging to distinguish from spurious events and might therefore be regarded with caution, as they may reflect biases in genealogical inference rather than true admixture. For such events, other evidence, such as the use of genomic “signatures” may be helpful for verification.

These approaches perform well in simulations (Figure 1, Supplementary Figures 1-6). Empirically, in real data, the number of components identified using this approach can be higher due to multiple admixture events in the history of a sample, or additional admixtures impacting putative sources. Thus, in practice, we select a conservative number of components based on how well the components can be interpreted in the context of real data. In most cases, this results in choosing two components. Ultimately, the choice of the number of components can be data-driven, guided by the held-out log-likelihood, expected *R*^2^, and co-ancestry dating, or informed by prior knowledge of the admixture event.

### Tree filtering

Relate trees have variable span along the genome. We only consider trees every 10kb or 0.05cM apart. We filter out trees in regions corresponding to centromeres, telomeres, and the HLA region, along with a 500kb buffer around these areas. The positions of centromeres, telomeres, and the HLA region in the appropriate genome build were obtained from the UCSC genome browser. For analyses of GCbGC around PRDM9 hotspots (e.g. Figure 4a), we additionally included trees spaced every 200 bp in the vicinity of these hotspots. For analyses of PGS portability (Figure 4d), we included trees overlapping the UK Biobank SNP markers used for PGS construction.

### Initialisation

To find a suitable initialisation, we perform a random search over 20 randomly initialised values for coalescence rates, proportions, and admixture times. The coalescence rates are sampled from a log-uniform distribution within the range 5 × 10^−2^to 1 × 10^−7^. The proportions are drawn from a uniform distribution between 0.01 and 0.99, while the admixture times are sampled from a log-uniform distribution spanning 20 to 2000 generations. For each random initialisation, the EM algorithm is run for 10 iterations. The initialisation that achieves the highest log-likelihood after these iterations is selected as the optimal starting point for further model fitting.

### GhostBuster output

GhostBuster outputs a per-component coalescence rate matrix, which records piecewise constant coalescence rates over time between the reference populations and each inferred ancestry component. It also outputs the estimated genome-wide ancestry proportions for each component and their local ancestry posterior probabilities across the genome.

The local ancestry posteriors are used for several downstream analyses. For instance, we infer co-ancestry curves to date admixture events following^33^, by correlating local ancestry posteriors across genomic distances. The expected coefficient of determination^34,35^, provides a genome-wide summary of local ancestry certainty (Supplementary Note). Additionally, by stratifying mutations or recombination events based on the inferred local ancestry surrounding them, GhostBuster enables the detection of ancestry-specific mutational signatures or recombination hotspot activity.

### Simulations

We tested GhostBuster on several simulations (Supplementary Note). All simulations were generated using msprime^70,71^.

### Two-way admixture simulation

We simulated three populations: A, B, and a focal population. The focal population was modelled to have 20% of its ancestry from population A and 80% from population B. All populations, including the merged ancestral population formed after populations A and B merge, were assigned a constant haploid effective population size of 10,000. We sampled up to 45 diploid individuals from populations A and B, and 5 diploid individuals from the focal population. To evaluate the robustness of GhostBuster across a range of scenarios, we varied the admixture time and the divergence time between populations A and B (Figure 1e). We assumed a constant mutation rate of 1.25 × 10^−8^ per base per generation and used the HapMap3 human recombination map (Data Availability).

### Four-way admixture simulation

We simulated a four-way admixture where each population contributed 25% to form the focal population; we provide further details in the Supplementary Note.

### Denisovan admixture in Papuans simulation

We used the simulation model from ^37^ to emulate 5% Denisovan admixture into Papuans, 43,935 years ago.

### Modern sample selection and processing

We used a recent unified version of HGDP^38,72^ and the 1,000 Genomes Project^39^, where the genomes from the two projects were jointly called and phased^73^. We used version 2 of this release (Data Availability).

This dataset was called with the hg38 genome build. The phased data had already undergone variant QC to filter variants with (1) HWE ≥ 1 × 10^−30^, (2) F_MISSING ≤ 0.1, or (3) ExcHet ≥ 0.5 and ExcHet ≤ 1.5. We additionally excluded multi-allelic SNPs and Indels from the dataset. We also performed sample QC, retaining only a subset of African samples from the 1,000 GP (20 individuals from each of 6 African subgroups: ACB, ASW, ESN, LWK, GWD, MSL) and removing close relatives up to the 2nd degree, inferred in the Allen Ancient DNA Resource^74^. These QC steps were implemented to minimise biases introduced by close relatives during genealogy inference and to prevent data imbalance between the two datasets. Our final dataset retained 45.6 million variants and 1,026 samples across 58 population groups.

We additionally analysed Simons Genome Diversity Project (SGDP)^40^ data, which was called in the hg37 genome build and was therefore not merged with the HGDP + 1,000 GP dataset. We treated it separately as an independent validation dataset. The phased data for SGDP were obtained from the Reich lab website (Data Availability). Apart from the filtering done in^40,74^, we additionally excluded multi-allelic SNPs and Indels from our analysis. We retained 28.3 million variants and 278 samples across 130 population groups in this dataset.

### Ancient sample selection and processing

We additionally considered 30 ancient and 4 archaic samples. These samples were selected to have >10x sequencing coverage, with an average coverage of 17.3 × (range: 10.4 − 65.2 ×). The ancient and archaic DNA samples used in the analysis are tabulated in Supplementary Table 1.

The ancient DNA BAM files were processed using bcftools mpileup to generate variant calls, selecting only reads with a mapping quality of 20 or higher, a base quality of 20 or higher, and a minimum depth of coverage of 5. The variant calls from the ancient DNA samples were then merged with the 1,000 GP (Data Availability) and phased using SHAPEIT4. We used a two-step phasing procedure, first phasing variants observed in the 1,000 GP Phase 3 reference panel and then using this phasing as a scaffold to phase variants not observed in the reference panel. We additionally randomly phased singletons. Subsequently, the ancient DNA dataset was lifted over to the hg38 reference genome using Picard LiftoverVcf and merged with the unified version of HGDP + 1,000 GP using the bcftools merge -0 option. After merging, we filtered out multi-allelic variants, duplicate variants, and indels. As a check, we compared allele frequencies between the HGDP + 1,000 GP dataset and the ancient DNA dataset, finding a high correlation between the allele frequencies with minimal missingness (Supplementary Figure 20).

### Details of Relate inference

We use Relate^15^ to construct genealogies for both modern and ancient samples. To identify the most likely ancestral allele for each SNP, we use the chimpanzee reference genome lifted over to the relevant human genome build (Data Availability).

For genealogies involving only modern samples, we apply the 1,000 Genomes Project pilot mask (lifting it for hg38 datasets), retaining only regions marked as “passing” to exclude variants with high uncertainty (Data Availability). Additionally, we mask sites where the chimp ancestral genome shows missing data, as well as sites corresponding to multi-allelic SNPs or indels. For genealogies that include archaic samples, we further refine this mask by considering only sites that pass in the Neanderthal and Denisovan masks (Data Availability). Additionally, for genealogies with ancient samples, we filter out sites where ancient samples exhibit more than 10% missingness before phasing and imputation. The masks corresponding to ancient and archaic samples are lifted over to the hg38 reference genome. We additionally filter all CpG site patterns in the ancestral genome to effectively remove CpGs when building trees with ancient or archaic samples.

We utilise the HapMap3-inferred recombination rate maps for builds hg37 and hg38 (Data Availability). The mutation rate is set to 1.25 × 10^−8^per base per generation. We modified Relate to enforce the construction of topologies every 10kb using the --fb option, in addition to any tree changes already inferred within Relate. When including ancient samples, we provide their estimated age. We assume 28 years per generation while running Relate and all subsequent analyses.

We estimate the joint effective population size of the dataset by iteratively fitting time-varying population sizes and branch lengths for chromosome 1, over five iterations. This converged dataset-wide population size estimate is then used as a prior for estimating branch lengths on the remaining chromosomes. Finally, the resulting trees are converted to tskit^71,75^ format, using the Convert function provided in the relate_lib package (Code Availability), where we have implemented an option to retain only the trees to be used for model fitting in GhostBuster to reduce file size.

Overall, we compile four datasets for which we infer genealogies:

1. **HGDP + 1,000GP**: Includes all modern samples from HGDP and a subset of African samples from the 1,000 Genomes Project.
2. **HGDP + 1,000GP + aDNA**: Includes HGDP + 1,000GP samples along with 30 high-coverage ancient DNA samples.
3. **HGDP + 1,000GP + aDNA + archaic**: Extends the HGDP + 1,000GP + aDNA set by adding 4 archaic samples (Vindija, Chagyrskaya, Altai Neanderthals and Denisovan).
4. **SGDP**: Contains all modern samples from SGDP.
5. **Modern Africans + archaic**: Includes African samples from the HGDP + 1,000GP, supplemented with one Neanderthal (selected from Altai, Chagyrskaya, or Vindija) and one Denisovan genome.

We confirmed consistency across genealogies inferred for each dataset by evaluating the inferred inverse coalescence rates between populations (Supplementary Figure 21). For most of our data analyses, we primarily focus on using samples from HGDP and 1,000GP, while incorporating SGDP samples to augment the analyses where feasible.

## Running GhostBuster on real data

### Finding Eurasian ancestry in Africans

We applied GhostBuster to detect Eurasian ancestry in African individuals using the HGDP + 1,000GP genealogies and HGDP + 1000GP + aDNA genealogies. Among reference populations, non-Finnish Europeans, East Asians, and South Asians were combined into a single Eurasian “super-population”, while African populations were treated separately. Groups known to possess strong recent admixture, including Middle Eastern and American individuals, were excluded from the reference panel to avoid confounding. GhostBuster was run in supervised mode with fixed coalescence rates for each ancestry using the Eurasian superpopulation and all African populations (except that the target sample is derived from) as references. We focused GhostBuster on the time period between 1,000 and 50,000 years ago, which was partitioned into 20 log-spaced epochs.

For local ancestry-based analyses involving mutational enrichment and Neanderthal enrichment, we only considered Eurasian segments at least 0.2 cM in length (corresponding to segments within the past 14,000YBP, approximately) and excluded shorter tracts (which might be associated with the deeper rather than more recent admixture event).

### Finding OOA/non-OOA ancestry in Africans

We applied GhostBuster to understand deeper admixture in African individuals using the HGDP + 1,000GP genealogies and HGDP + 1000GP + aDNA genealogies. Similar to the back-to-Africa analysis, non-Finnish Europeans, East Asians, and South Asians were combined into a single Eurasian superpopulation, while African populations were treated separately. Middle Eastern and American individuals were excluded from the reference panel. GhostBuster was run in unsupervised mode to fit two components. Analyses were restricted to an older time period, focusing on 50,000 and 300,000 years ago, partitioned into 20 log-spaced epochs. Additionally, given the ancient nature of the event, GhostBuster was run without the HMM, treating each tree as independent of others.

For local ancestry–based analyses of mutational enrichment and GCbGC, we excluded regions within ±100 kb of loci where the posterior probability of Eurasian ancestry (associated with back-to-Africa) inferred by GhostBuster exceeded 0.1. This was done to focus on the deeper ancestral signal rather than more recent admixture events. However, for coancestry curve–based dating, local ancestry tracts associated with Eurasian gene flow were retained, as these segments are essential for accurate inference of admixture timing.

### Finding Neanderthal ancestry in Africans and non-Africans

We applied GhostBuster to detect Neanderthal ancestry in African modern and ancient individuals using the HGDP + 1,000GP + aDNA + archaic dataset. To detect Neanderthal ancestry, we ran GhostBuster in supervised mode. We focused on coalescence events between 50,000 and 2 million years ago, excluding more recent events to minimise the influence of post-admixture demography. To isolate Neanderthal-to-modern human gene flow, we used Neanderthals as the only reference group, which excluded all branches not involving Neanderthals. We fitted two components, one defined by genome-wide coalescence rates between Neanderthals and another defined by coalescence rates between Africans and Neanderthals, both derived from the genealogies themselves.

To validate our approach, we first estimated Neanderthal ancestry in six modern Eurasian populations (Han, Japanese, French, Russian, Pathan, and Brahui). We recovered genome-wide Neanderthal ancestry proportions of ∼0.9–1.1%, consistent with previous studies^43,76^. East Asian populations exhibited ∼20% higher Neanderthal ancestry compared to Europeans and South Asians, mirroring patterns reported in prior work^44,77^. In addition, we detected minimal to no Neanderthal ancestry in genomic regions previously identified as "archaic deserts"^78,79^ (Supplementary Figure 22).

### Finding “Human-like” ancestry in Neanderthals

We applied GhostBuster to the Modern Africans + archaic dataset to understand the human-like (HA) gene-flow into Neanderthals. We used Africans and Denisovans as reference populations, focusing on coalescence events between 100,000 and 2 million years ago, partitioned into 20 log-spaced bins. GhostBuster was run in unsupervised mode to fit two ancestry components, with the admixture proportions constrained to 10% and 90% to stabilise inference. We also excluded trees where Neanderthal and Denisovan lineages coalesced more recently than 300 KYA to avoid confounding from recent gene flow between Neanderthal and Denisovan groups. Each Neanderthal individual was analysed separately. Similar to finding OOA/non-OOA ancestry in Africa, GhostBuster was run without the HMM.

### Mutational enrichment analyses

To investigate if certain mutation types are enriched or depleted in inferred admixture backgrounds, we define a normalised mutation rate. For each mutation type (e.g., TCC to TTC), the normalised mutation rate is defined similarly to ^15^. We calculate the ratio of the number of mutations of that specific type in a population to the number of mutations of all other types in the same population. Formally, this is expressed as

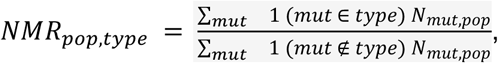

where the sum is over all mutations in the genealogy, *N_mut,pop_* counts the number of individuals belonging to the population ‘pop’ carrying mutation ‘mut’. We define an extension where we additionally stratify by allele age (defined as the midpoint of the branch carrying this mutation). We do this by counting *N^t^_mut,pop_*, the number of carriers of a mutation arising in time interval *t*, dividing time into 10 log-spaced epochs ranging from 1,000 years to 1 million years.

To facilitate comparisons across populations, we further divide these normalised mutation rates (NMR) by the normalised mutation rates (NMR) in African populations by computing 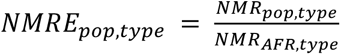. We refer to this quantity as the Normalised Mutation Rate Enrichment (NMRE). A significant deviation from unity indicates that the specific mutation type is either enriched (greater than 1) or depleted (less than 1) in the population relative to the African baseline. We examine this enrichment across various continental populations, including all non-Finnish Europeans (NFE), East Asians (EAS), South Asians (SAS) and Middle Eastern and North African (MID) groups in HGDP.

To estimate the NMRE for different ancestry components inferred by GhostBuster, we first discretised the local ancestry posteriors by thresholding at 0.5. The local ancestry assignment for each target haplotype and mutation was defined using the nearest local ancestry call, and mutations located more than 5 kb away from any local ancestry call were excluded from the analysis. We further downsampled individuals at any given site such that we had an equal number of descendants from each ancestry type. We expect this to correct for potential mutational biases in local ancestry inference. Ancestry components were then analysed separately by counting, for each mutation, the number of individuals of a given ancestry type who carry that mutation. To compute NMREs, we normalised these NMRs by the genome-wide NMRs estimated for all African samples in the dataset. Confidence intervals around the NMRE estimates were computed using 1,000 chromosome-wise bootstrap replicates. NMREs for TCC to TTC and weak to strong mutations are presented in Figure 2e, Extended Data Figure 2, 5, and for all 96 trinucleotide types shown in Supplementary Figure 11,17.

### Identifying source populations for the Back-to-Africa events

To identify putative source groups, we computed coalescence rates between inferred local ancestry tracts and samples not used in the GhostBuster fitting. We used the CoalRate function with the --mode local_ancestry flag (Code Availability) and focused on 0 to 10,000 years ago. Ancient samples were grouped into seven categories: Scandinavian Hunter-Gatherers (SHG), Yamnaya, Western Hunter-Gatherers (WHG), Irish Neolithic farmers, Balkan Neolithic farmers, Anatolian Neolithic farmers, and Early Neolithic Linearbandkeramik (LBK). We then constructed a coalescence rate matrix quantifying the coalescence rates of Eurasian-like segments with other modern and ancient groups. Finally, we generated a dendrogram of African samples using the UPGMA algorithm based on this coalescence rate matrix.

### Dating the Back-to-Africa event

We consider the presence of Neanderthal ancestry as a hallmark of the back-migration event. We first identify continuous Eurasian-like local ancestry segments by merging local ancestry gridpoints that have Eurasian-like posterior probabilities greater than 0.5. We then subset these segments to retain only those containing at least one gridpoint with a confident Neanderthal ancestry call (posterior > 0.5). Gridpoints within these segments are coded as 1, and all other gridpoints as 0. We then construct co-ancestry curves within and across the Eurasian and non-Eurasian updated local ancestry calls to date the event.

### Estimating coalescence rates between deep ancestry components

We estimate the coalescence rates between ancestry components (e.g. OOA-like with OOA-like, or OOA-like with non-OOA-like in Figure 3b) using the CoalRate function with the --mode local_ancestry flag (Code Availability). We thresholded the GhostBuster-inferred local ancestry at 0.8 to classify segments as either OOA-like or non-OOA-like while running CoalRate. As with outputs from GhostBuster and Relate, CoalRate produces a stepwise continuous coalescence-rate tensor across predefined time epochs. To quantify uncertainty in these estimates, we computed standard errors using 1,000 bootstrap replicates across genomic blocks.

### GCbGC at PRDM9 hotspots

In order to understand differences in recombination landscapes between different populations or ancestries, we analysed the intensity of GCbGC at PRDM9-A and PRDM9-C hotspots along the genome. Hotspot positions were obtained from previous studies^80,81^, filtered by hotspot heat and proximity to other hotspot types, and analysed independently (Supplementary Note). Furthermore, to avoid confounding effects of hypermutability, CpG-forming sites were excluded from all analyses.

We focused on mutations from A or T to G or C (weak to strong) to quantify and visualise the effect of GCbGC around hotspot centres across populations and different mutation sets. To do this, we divided the region spanning 10 kilobases upstream and downstream of each hotspot centre into 200 base pair bins. In each bin, we recorded the number of weak to strong mutations. To account for differences in local base composition, mutation counts were further normalised by the local AT vs GC fraction in the ancestral genome, calculated from the 200 base pair sequence window surrounding each mutation. We further normalise this mutation rate estimate by the background expectation obtained by averaging mutation rates 5-10kb away from the hotspot centre. We refer to the resulting metric as relative weak to strong mutation rate, which we plot as a function of distance from the hotspot centre in Figure 5f-g, Extended Data Figure 4b-c, 8 and Supplementary Figure 23 summarising patterns across all hotspots genome-wide.

To isolate and more formally test the effect of GCbGC across different ancestry components (as done in Figure 4a, 5h), we define a modified measure called weak to strong enrichment. Weak to strong enrichment is defined as the ratio of the number of weak to strong mutation carriers to the number of strong to weak mutation carriers, normalising each term by its relevant base compositions. We estimate the weak to strong enrichment across different ancestry backgrounds by discretising the posterior and restricting the analysis to individuals exhibiting the same local ancestry within a ±10 kb window surrounding the hotspot. We stratify this measure as a function of distance from the hotspot centre. To control for potential mutational biases introduced during local ancestry inference, we downsampled ancestry segments to ensure equal representation across ancestry types. Looking at the ratio of weak to strong and strong to weak mutations enables correcting for overall mutation rate differences between ancestries or some other conditioning. Finally, to formally test for GCbGC differences between OOA-like and non-OOA-like ancestries, we compared the weak to strong enrichment 500bp around the hotspot centre (± 250bp), obtaining confidence intervals and p-values using 1000 bootstraps along the genome.

To evaluate the effect of hominin-to-archaic introgression on the recombination landscape of archaic genomes (as shown in Figure 5h), we analysed GCbGC at PRDM9 hotspots by stratifying them based on how human-like the surrounding archaic sequence appears. Specifically, we categorised hotspots based on minimum distance from a likely “introgressed” SNP, defined as a SNP at least 1kb away from any hotspot centre, having descendants in Human and Neanderthal, but not Denisovan, with estimated allele age (in the Modern Africans + archaic genealogy) < 500kya (Supplementary Note). Similar to the analysis comparing GCbGC between OOA-like and non-OOA-like ancestry, we computed weak to strong enrichment ± 250bp around the hotspot centre, computing confidence using exact binomial testing (Supplementary Note).

We performed additional PRDM9-based analyses to further characterise hotspot activity across hominin lineages. Analysis of GCbGC across human, chimpanzee and gorilla genomes confirmed strong PRDM9-A activity on the ancestral human lineage and supports a more recent, derived origin for PRDM9-C (Supplementary Figure 23, Supplementary Note). We further complemented these analyses with tests of “hotspot death”, based on motif-disrupting mutations, which provide a stronger and more direct signature of historical PRDM9 binding. These results are highly consistent with the GCbGC-based inferences, reinforcing distinct temporal and population-specific activity of PRDM9-A and PRDM9-C hotspots (Extended Data Figure 9, Supplementary Figure 24, Supplementary Note).

### Testing for large introgressed regions in Neanderthals

To identify selection signals, we scanned the genome for large introgressed regions using 500 kb windows and three complementary measures of human–Neanderthal introgression: (i) GhostBuster-inferred introgression probability, (ii) human–Neanderthal coalescence time, and (iii) the fraction of SNPs shared between humans and Neanderthals but absent in Denisovans. These measures were standardised and combined into a single genome-wide statistic to detect regions with excess introgression. The top-ranked region mapped to the PRDM9 locus, showing strong evidence of introgression, with elevated human–Neanderthal sharing and increased Neanderthal–Denisovan divergence (Supplementary Note; Supplementary Table 3a).

### PRDM9 typing in archaics

We inferred PRDM9 alleles for Neanderthals, Denisovans, and additional individuals (including Ust’-Ishim and a Neanderthal–Denisovan hybrid) using raw sequence reads from the PRDM9 region. We processed each archaic separately, using a likelihood-based approach, previously applied to humans^53^. Further methodological details and results are provided in the Supplementary Note and Supplementary Table 3b.

### Details of ANCHOR analysis

We applied the ANCHOR analysis pipeline^57^ that enables quantifying polygenic score (PGS) portability depending on local ancestry calls (Supplementary Note). We applied this framework to assess portability differences between the OOA-like and non-OOA-like ancestries, which we show in Figure 4d.

### Estimating human-like ancestry proportion in Neanderthal

To estimate the proportion of human-like ancestry in Neanderthals, we develop a bespoke approach comparing Neanderthal–human and Denisovan–human coalescence patterns. We assume that Denisovans did not receive gene flow from another hominin group and therefore use Denisovan–human coalescences as a baseline representing unadmixed archaic ancestry. Hominin introgressed segments in Neanderthals are therefore expected to coalesce with human lineages more recently than non-admixed regions.

To define our estimator, we compute the probability that the human–Neanderthal coalescence time *T*_*HN*_ is below a threshold *T*. Let *λ* denote the fraction of genomic positions in Neanderthals that carry introgressed human-like ancestry. Under the assumption that, outside introgressed regions, the distribution of human–Neanderthal coalescence times matches that of human–Denisovan coalescence times, we obtain:

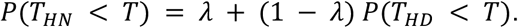

We can solve this equation for *λ* and obtain the estimator.

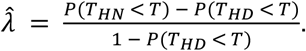

We estimate these probabilities empirically using Relate-inferred genealogies across the genome, weighting trees by their genomic span (equivalent to sampling random genomic positions). To minimise biases arising from differences in sample ages, we focus primarily on the Chagyrskaya Neanderthal, whose age is comparable to that of the Denisovan individual used. We further exclude outlier trees in which the inferred Neanderthal–Denisovan coalescence time exceeds 3 million years (2.6% of trees), as these may reflect inference artefacts or potential super-archaic admixture.

We estimate 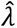 across a range of time thresholds *T*. For small *T*, the estimator underestimates introgression because not all admixed segments coalesce before *T*, whereas for large *T*, most lineages coalesce irrespective of admixture status, increasing uncertainty. To balance these effects, we compute estimates across 50 KYA intervals between 500 KYA and 2.5 MYA and assess uncertainty using block bootstrapping (blocks of 1000 trees).

Estimates increase with *T* and plateau around *T* ≈ 1.25 MYA, where 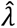 ≈ 10.1%. This value is stable for larger *T*, with estimates ranging from 8.5% to 12.3%, all within overlapping 95% bootstrap confidence intervals (Extended Data Figure 7). Consistent estimates are observed across individuals (Altai: 9.89%; Vindija: 11.0%), indicating limited sensitivity to sample age differences for sufficiently large *T*. Based on these analyses, we adopt an approximate value of 10% for downstream analyses.

We verified this proportion estimate using an independent approach based on local coalescence patterns at the PRDM9 locus, assuming the locus to be fully introgressed. By comparing mean coalescence times in this region to genome-wide expectations, we obtained a consistent estimate of ∼10% introgressed ancestry in Neanderthals (Supplementary Note).

### D-statistic calculations for gene flow into Neanderthal

To investigate the nature and timing of gene flow into Neanderthals, we compute a D-statistic^82,83^, which is defined as

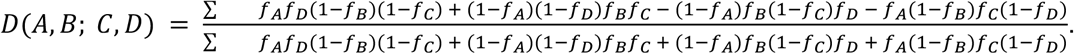

Here, *f*_*X*_ denotes the population-level observed allele-frequency of population *X*. The sum is over all mutations considered. The D-statistics calculates the normalised difference between the expected number of ABBA and BABA site patterns. A positive D-statistic indicates greater allele sharing between populations A and D or B and C and a negative value indicates greater allele sharing between population A and C, or B and D. A D-statistic value indistinguishable from 0, indicates symmetric relation of populations A or B to populations C or D. To assess statistical confidence, standard errors are estimated via a block jackknife procedure, dividing the genome into 500 contiguous equal-sized blocks.

To resolve the temporal dynamics of introgression, we stratify mutations either by their allele frequency in African populations^43^ or by their estimated allele age^61^, defined as the midpoint of the branch on which the mutation arose in the Modern Africans + archaic genealogy. We exclude regions overlapping PRDM9 recombination hotspots ± 10kb to avoid artefacts caused by recombinational differences between archaics and modern humans. Additionally, in calculating the D-statistic between Africans, Chimpanzee, Neanderthal, and Denisovan (Extended Data Figure 6), we remove the effect of back-to-Africa migrations by calculating allele frequencies conditioning out Eurasian local ancestry inferred by GhostBuster (posterior ≥ 0.1). We also remove the effect of introgression into archaics by removing trees with recent archaic-human coalescence.

Additionally, we also use D-statistics to assess the relationship of archaics to OOA-like and non-OOA-like populations. For this analysis, we restrict the analysis to mutations located within Neanderthal deserts, defined as regions at least 500 kb in length where the average Neanderthal ancestry in Eurasians is <0.1%, to prevent bias due to the presence of Neanderthal introgression in the reference populations used for local ancestry inference. Finally, in Extended Data Figure 6b, we also compute D-statistics on mutations simulated on genealogies using twigstats^61^.

---

**Extended Data Figure 1:**
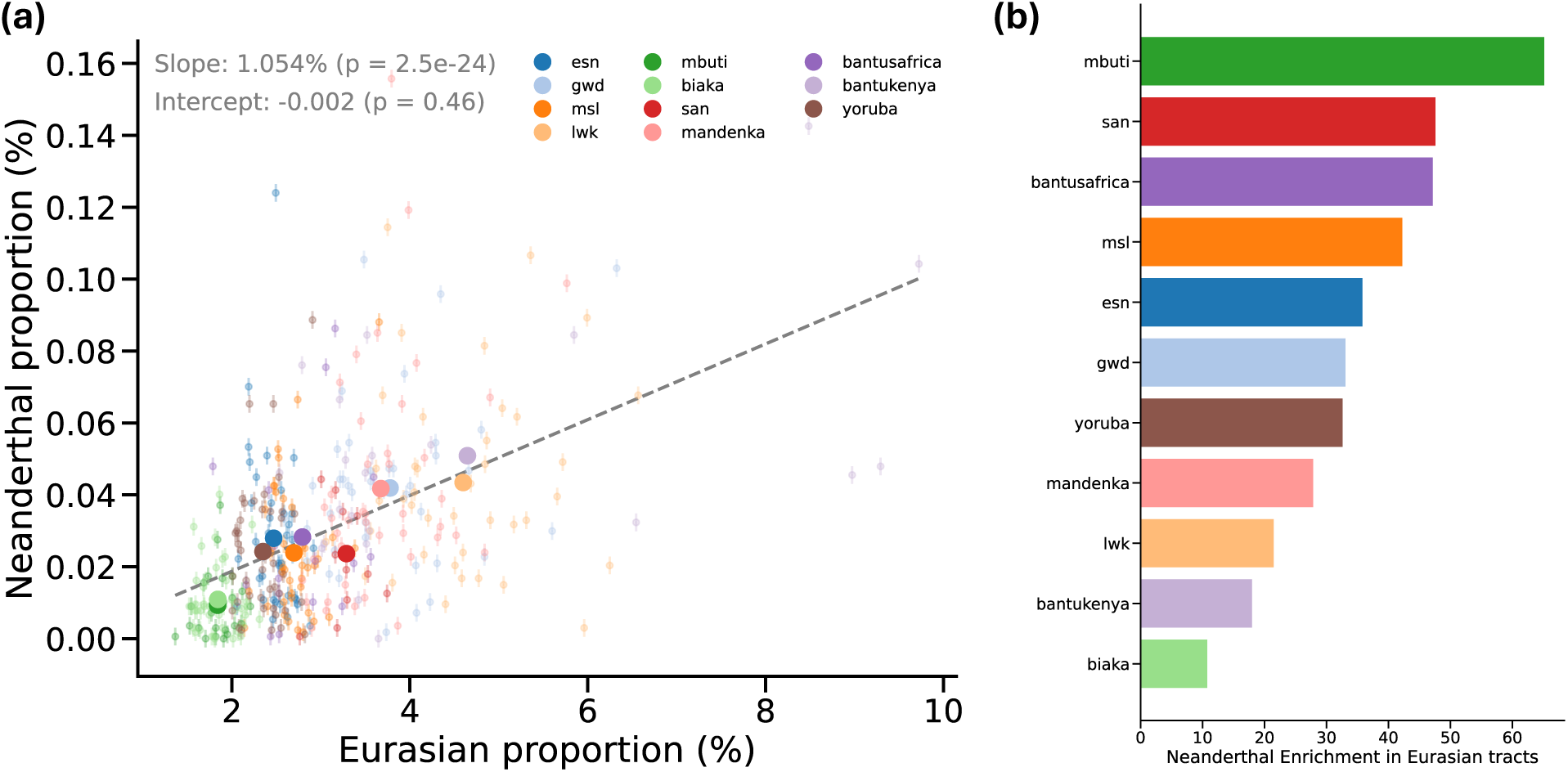
Neanderthal ancestry in African populations. (a) Relationship between Eurasian ancestry proportion (x-axis) and inferred Neanderthal ancestry proportion (y-axis) across modern African populations. Each point represents an individual, colored by population. Larger points denote population means. The dashed line indicates the linear fit across individuals (slope = 1.054%, *P* = 2.5 × 10⁻²⁴; intercept not significantly different from zero, *P* = 0.46), highlighting a strong correlation between Eurasian ancestry and Neanderthal ancestry, and no Neanderthal ancestry is predicted in individuals lacking Eurasian ancestry. (b) Fold-enrichment of Neanderthal ancestry within inferred Eurasian-like tracts across African populations. Bars represent the relative increase in Neanderthal ancestry within Eurasian ancestry segments compared to non-Eurasian ancestry segments, stratified by population. Populations are ordered by decreasing enrichment.

**Extended Data Figure 2:**
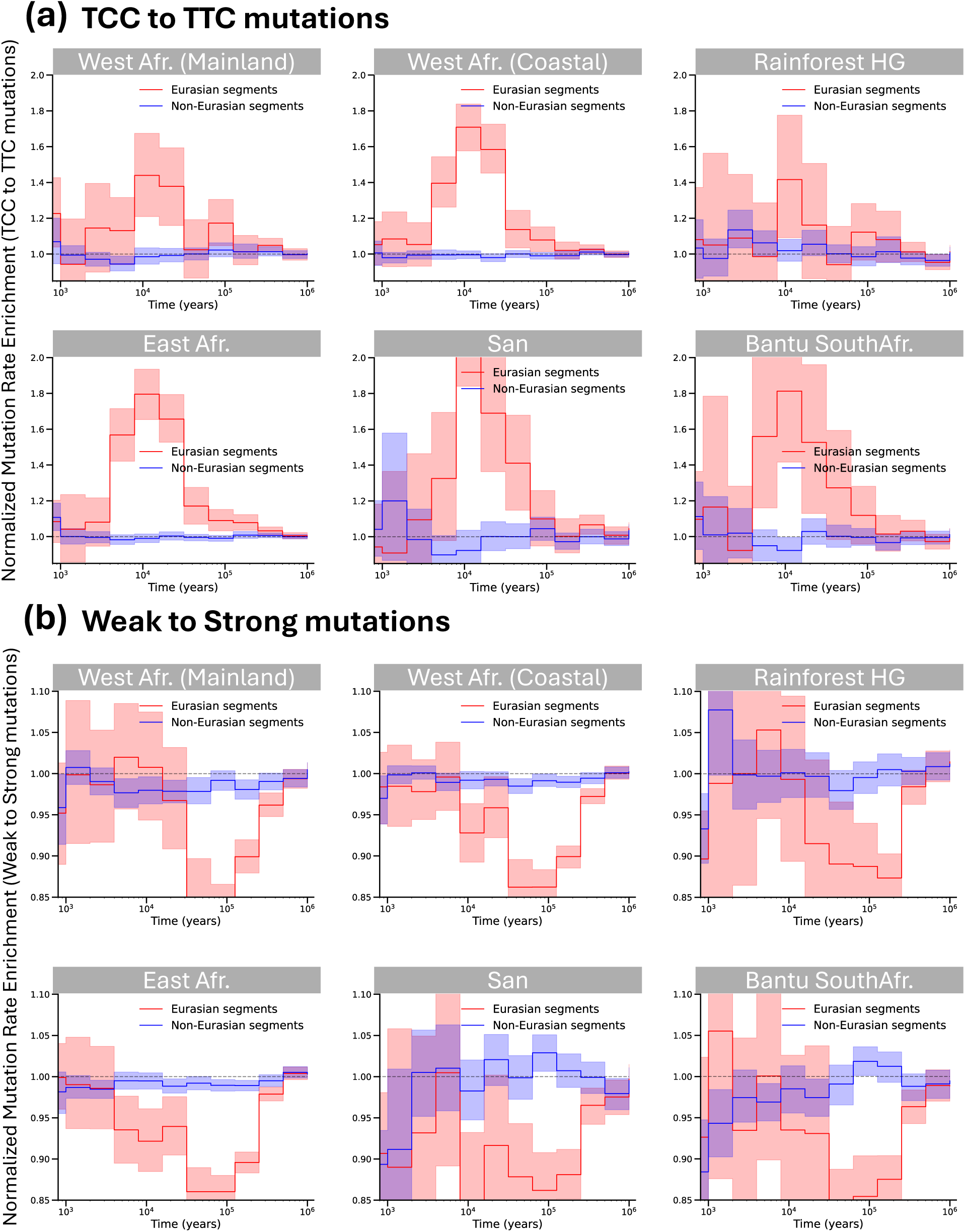
Mutational signatures within Eurasian-like ancestry segments. (a) Normalised mutation rate enrichment of TCC→TTC substitutions through time across African population groups. Panels show West Africans (mainland), West Africans (coastal), Rainforest hunter-gatherers, East Africans, San, and Bantu South Africans. Red and blue curves denote mutations occurring within inferred Eurasian-like and non-Eurasian ancestry segments, respectively. Shaded regions represent 95% confidence intervals estimated using bootstrap resampling. Eurasian ancestry segments show a pronounced enrichment of TCC→TTC mutations at peaking around 10⁴ years ago, consistent across multiple populations. (b) Normalised mutation rate enrichment of weak-to-strong (W→S) substitutions through time across the same population groups. As in (a), red and blue curves correspond to Eurasian-like and non-Eurasian ancestry segments, respectively, with shaded regions indicating 95% confidence intervals. West Africans (mainland) = Yoruba, ESN; West Africans (coastal) = GWD, MSL, Mandenka; Rainforest HG = Mbuti, Biaka; East Africans = LWK, BantuKenya.

**Extended Data Figure 3:**
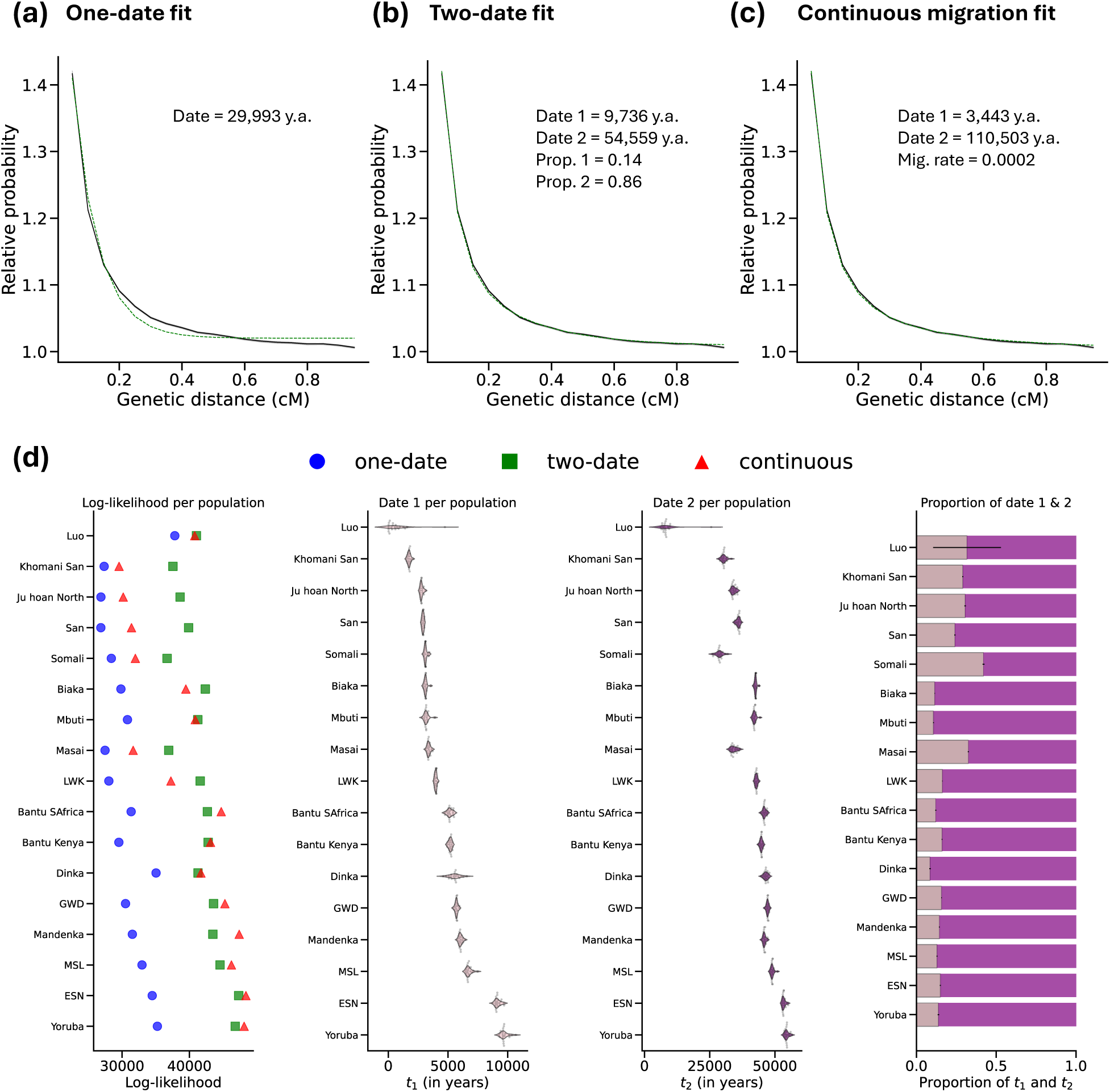
Dating the deep OOA/non-OOA admixture event in African populations. (a–c) Co-ancestry curve fits for Yoruba under three admixture models: (a) single-pulse (one-date), (b) two-pulse (two-date), and (c) continuous migration. Solid lines denote observed co-ancestry decay, and dashed lines indicate model fits. Parameter estimates for each model are shown within panels, including inferred admixture dates (in years ago), mixture proportions, and migration rate for the continuous model. (d) Model comparison and inferred admixture parameters across populations. Left: log-likelihood per population for each model (one-date, two-date, and continuous), aggregated across 20 jackknife replicates. Middle panels: inferred admixture dates from the two-date model, showing the younger (t₁) and older (t₂) components (in years ago). Right: corresponding mixture proportions for the two components, with pink indicating the more recent pulse and purple the older pulse. Points denote individual jackknife estimates overlaid on violin plots. Local ancestry tracts corresponding to Eurasian gene flow were retained for this analysis to allow full date inference.

**Extended Data Figure 4:**
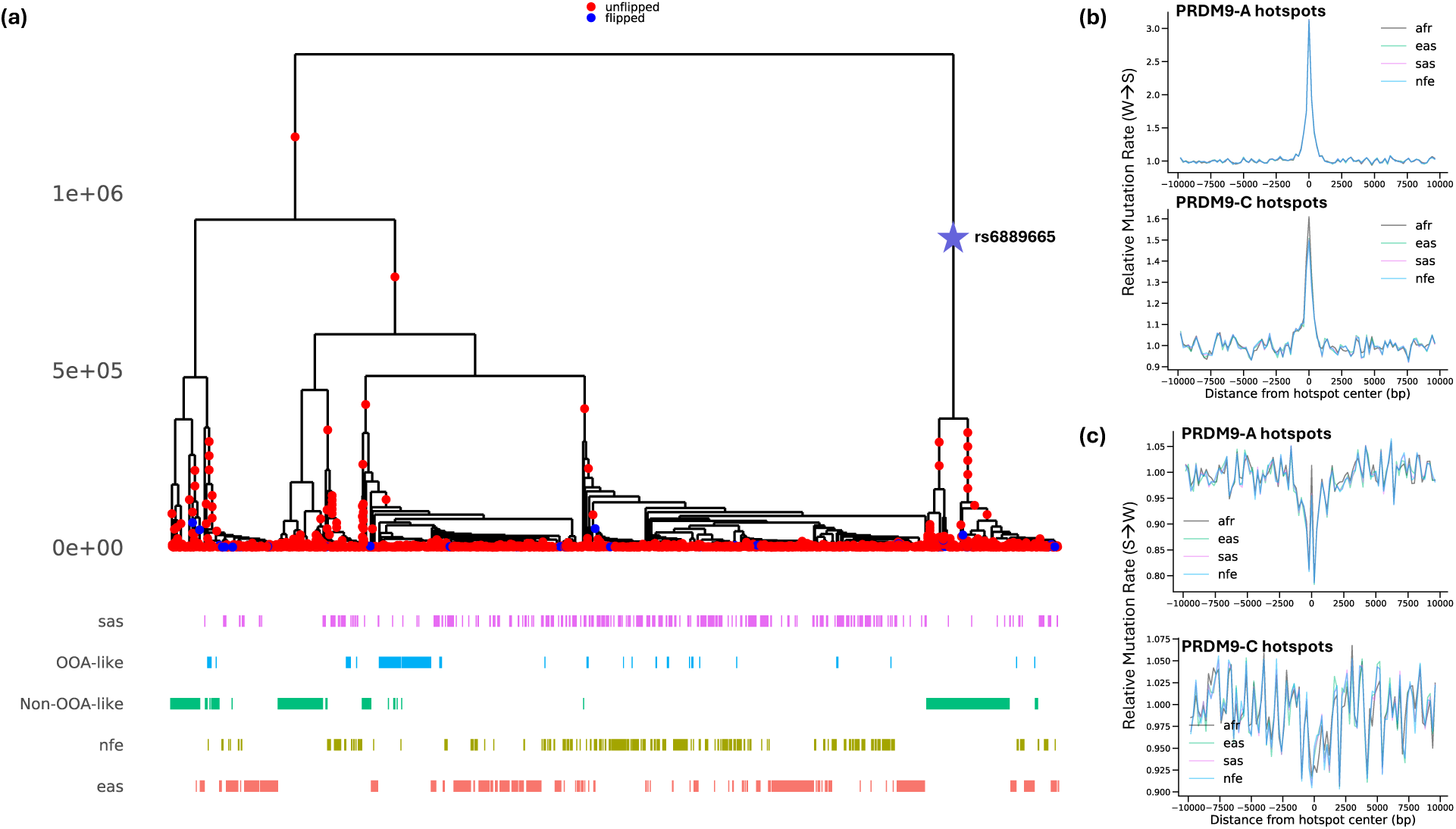
Relate-inferred genealogy at the PRDM9 locus and GCbGC in global populations. (a) Local genealogical tree inferred by Relate at the PRDM9 locus. The y-axis measures time in years before present; variant rs6889665 tagging the PRDM9-C allele is indicated as a star. The lower panel shows the distribution of haplotypes across populations (SAS, NFE, EAS) and inferred ancestry components (OOA-like and non-OOA-like). (b) Relative mutation rate of weak-to-strong (W→S) substitutions around PRDM9-A (top) and PRDM9-C (bottom) hotspots across populations. (c) Relative mutation rate of strong-to-weak (S→W) substitutions around PRDM9-A (top) and PRDM9-C (bottom) hotspots across the same populations. AFR = Africans, EAS = East Asians, SAS = South Asians, NFE = Non-Finnish Europeans.

**Extended Data Figure 5:**
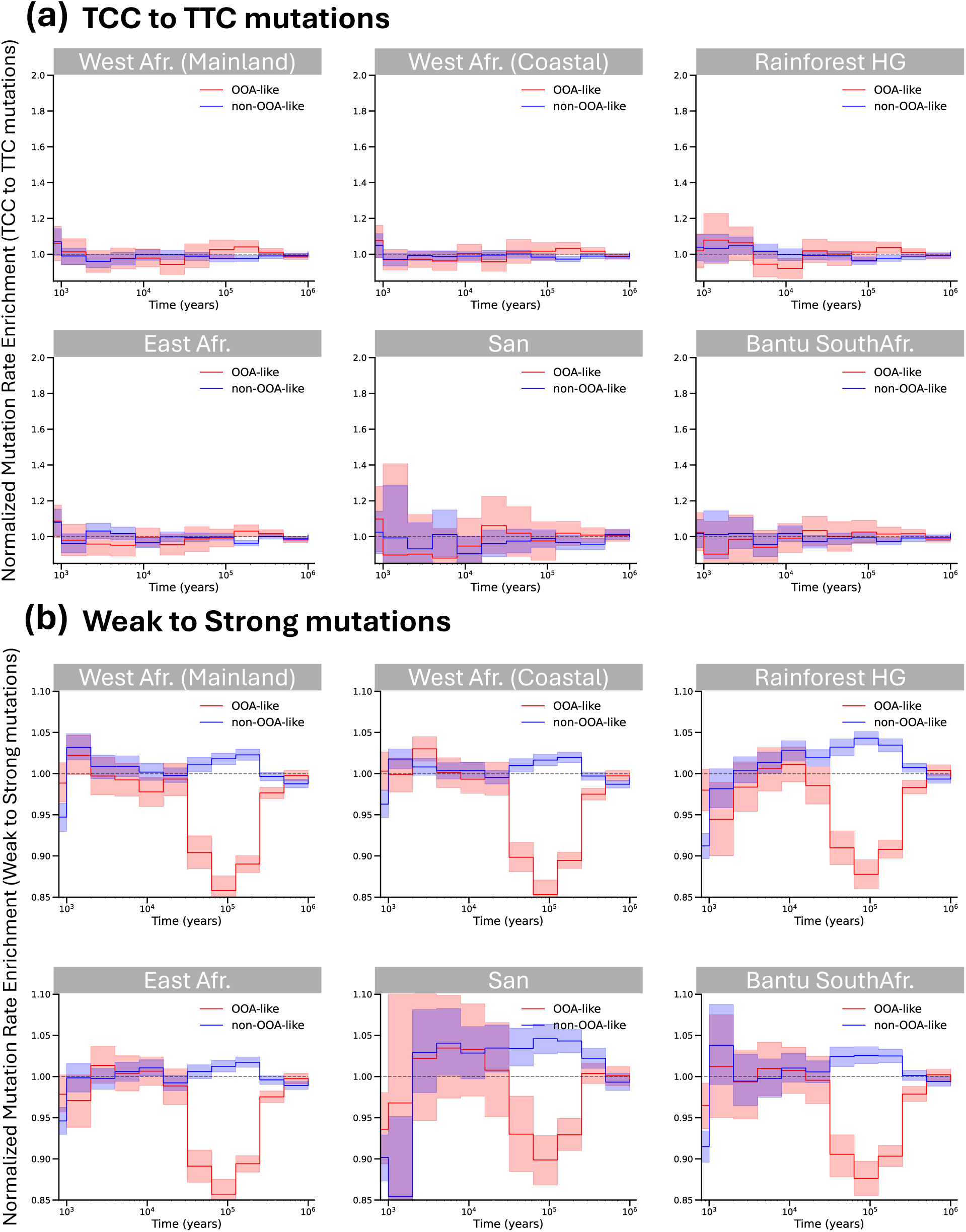
Mutational signatures of the deep admixture event. (a) Normalised mutation rate enrichment of TCC→TTC substitutions through time across African population groups. Panels show West Africans (mainland), West Africans (coastal), Rainforest hunter-gatherers, East Africans, San, and Bantu South Africans. Red and blue curves denote mutations occurring within OOA-like and non-OOA-like ancestry segments, respectively. Shaded regions represent 95% confidence intervals estimated using bootstrap resampling. (b) Normalised mutation rate enrichment of weak-to-strong (W→S) substitutions through time across the same population groups. As in (a), red and blue curves correspond to OOA-like and non-OOA-like ancestry segments, respectively, with shaded regions indicating 95% confidence intervals. OOA-like segments show a relative depletion of W→S mutations at intermediate timescales (∼3×10^4^–3×10^5^ years), while non-OOA-like segments remain close to baseline across populations. Population definitions (titles) are as for Extended Data Figure 2. Eurasian gene flow was removed to calculate the relative mutation rates.

**Extended Data Figure 6:**
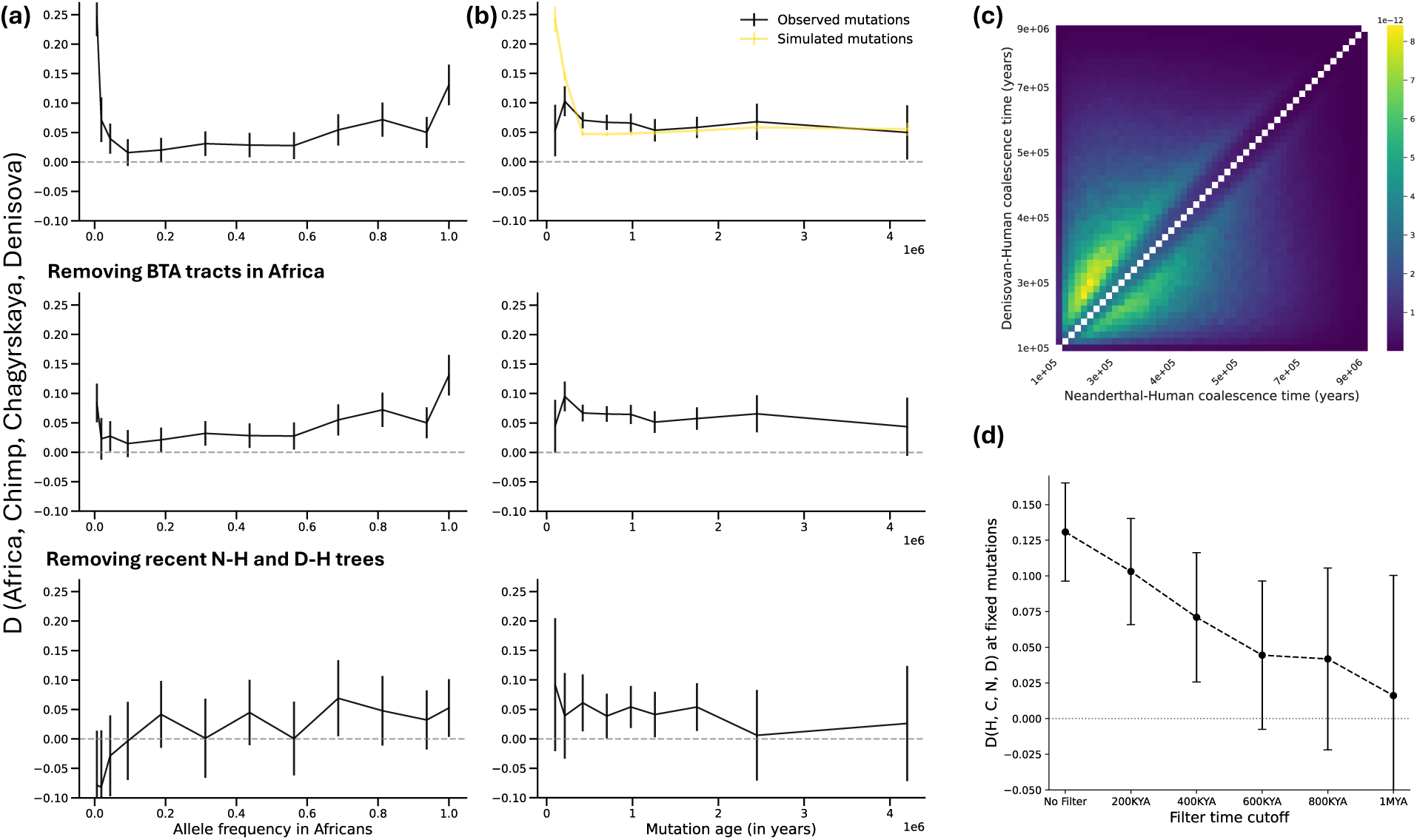
D-statistic stratified by mutation age and allele frequency. (a,b) D-statistics comparing allele sharing between Africans, Chimpanzee, the Chagyrskaya Neanderthal, and Denisovan, stratified by (a) allele frequency in Africans and (b) mutation age. Top: all sites; middle: after removing Back-to-Africa (BTA) ancestry tracts; bottom: after additionally excluding trees with recent Neanderthal–human or Denisovan–human coalescence (first N-H or D-H coalescence < 500KYA). Error bars denote 95% confidence intervals. (c) Joint distribution of Neanderthal–human and Denisovan–human coalescence times inferred from Relate. Density is concentrated above the diagonal, consistent with more recent Neanderthal–human coalescence relative to Denisovan–human across a subset of the genome. (d) D-statistics in (a) restricted to fixed mutations in Africans. For each fixed mutation, we use the genealogy to determine the maximum time of first coalescence of an African lineage with the Chagyrskaya Neanderthal or the Denisovan. We filter fixed mutations, excluding those with a maximum first coalescence time younger than the date shown on the x-axis. This shows attenuation of the signal as more human-like introgression signal is removed. Error bars denote 95% confidence intervals.

**Extended Data Figure 7:**
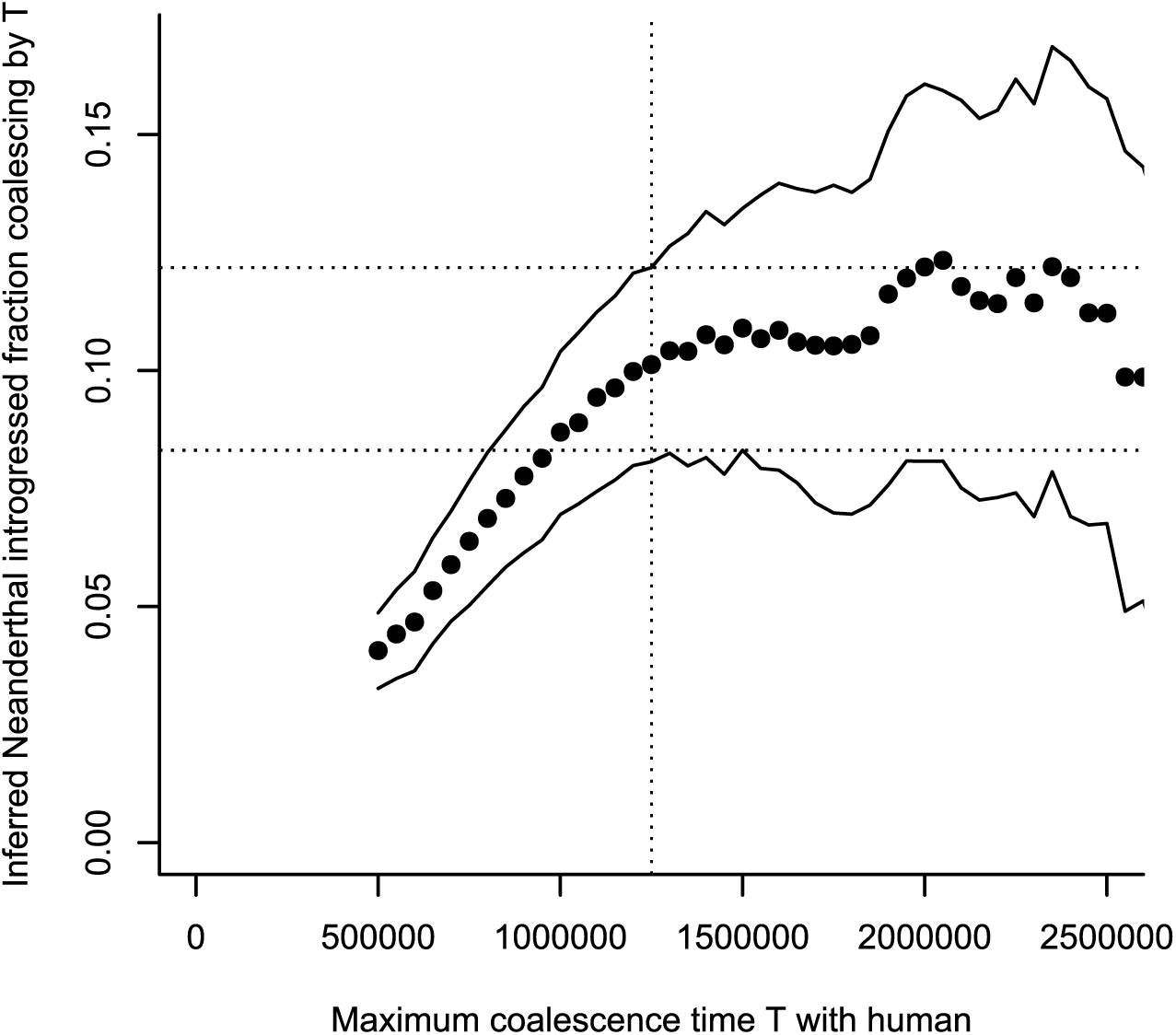
Genome-wide estimate of human-like introgression into Neanderthals. Estimated proportion of introgressed human-like ancestry in Neanderthals as a function of the maximum coalescence time threshold *T*. Points denote estimates computed from the difference between human–Neanderthal and human–Denisovan coalescence probabilities (Methods). Solid lines indicate 95% confidence intervals obtained by block bootstrap resampling. Estimates increase as expected with *T*, and plateau at an admixture estimate of around ∼10% at *T* ≈ 1.25 million years (dashed line).

**Extended Data Figure 8:**
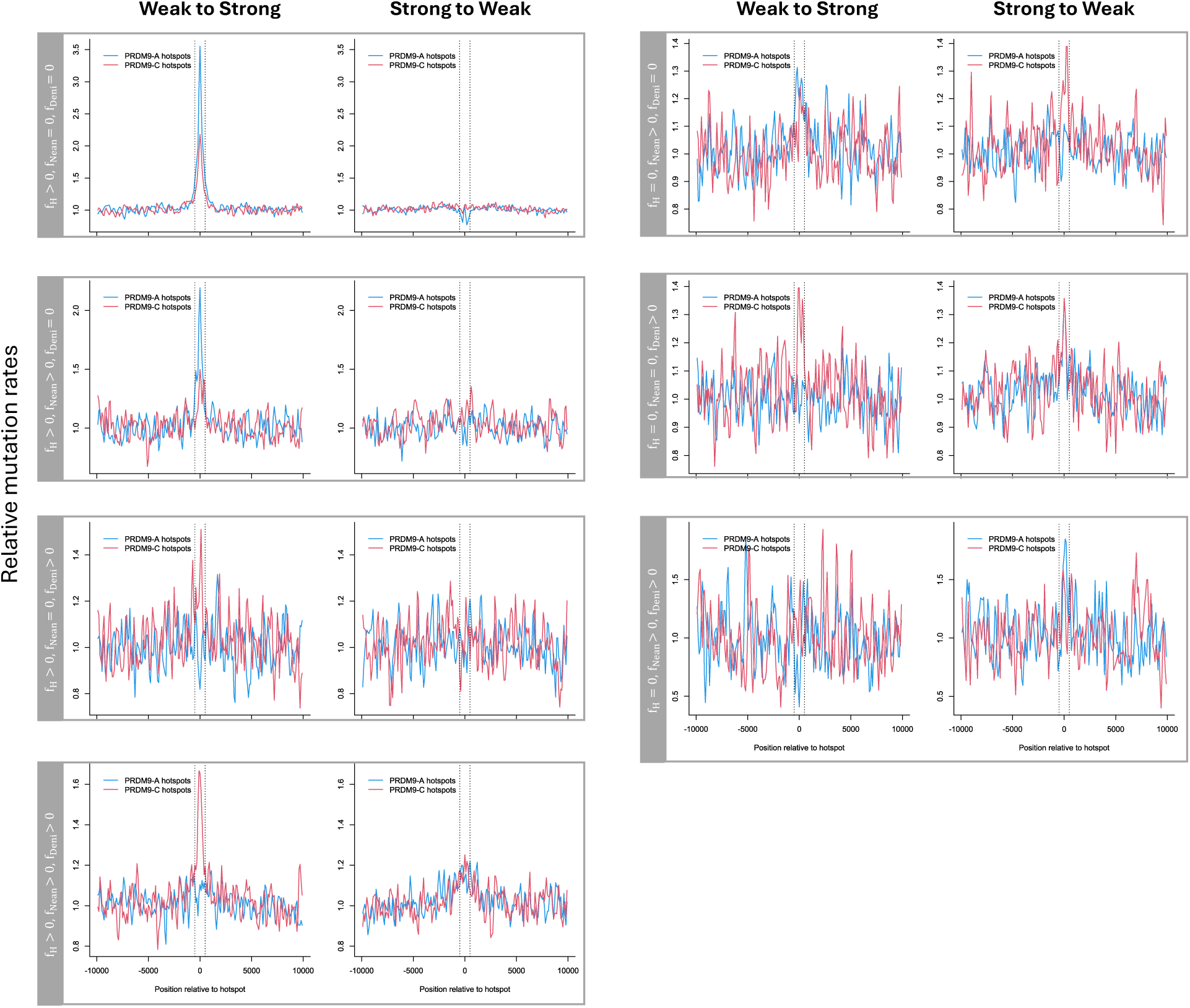
GCbGC in archaic hominins and humans across different mutation sets. Relative mutation rates of weak-to-strong (W→S) and strong-to-weak (S→W) substitutions around PRDM9-A (blue) and PRDM9-C (red) recombination hotspots. Rows correspond to mutation sets defined by sharing patterns across humans (H), Neanderthals (N), and Denisovans (D), as indicated. Note that different plots have different y-axis scales. Only mutations segregating in the combined human, Neanderthal, and Denisovan dataset are used. Relative mutation rates are shown as a function of distance from hotspot centres (dashed lines) and normalised to flanking regions. Enrichment of W→S mutations at hotspot centres is observed primarily in human-only and human–Neanderthal shared mutation sets at PRDM9-A hotspots, and in human-only and human–Neanderthal–Denisovan shared mutation sets at PRDM9-C hotspots.

**Extended Data Figure 9:**
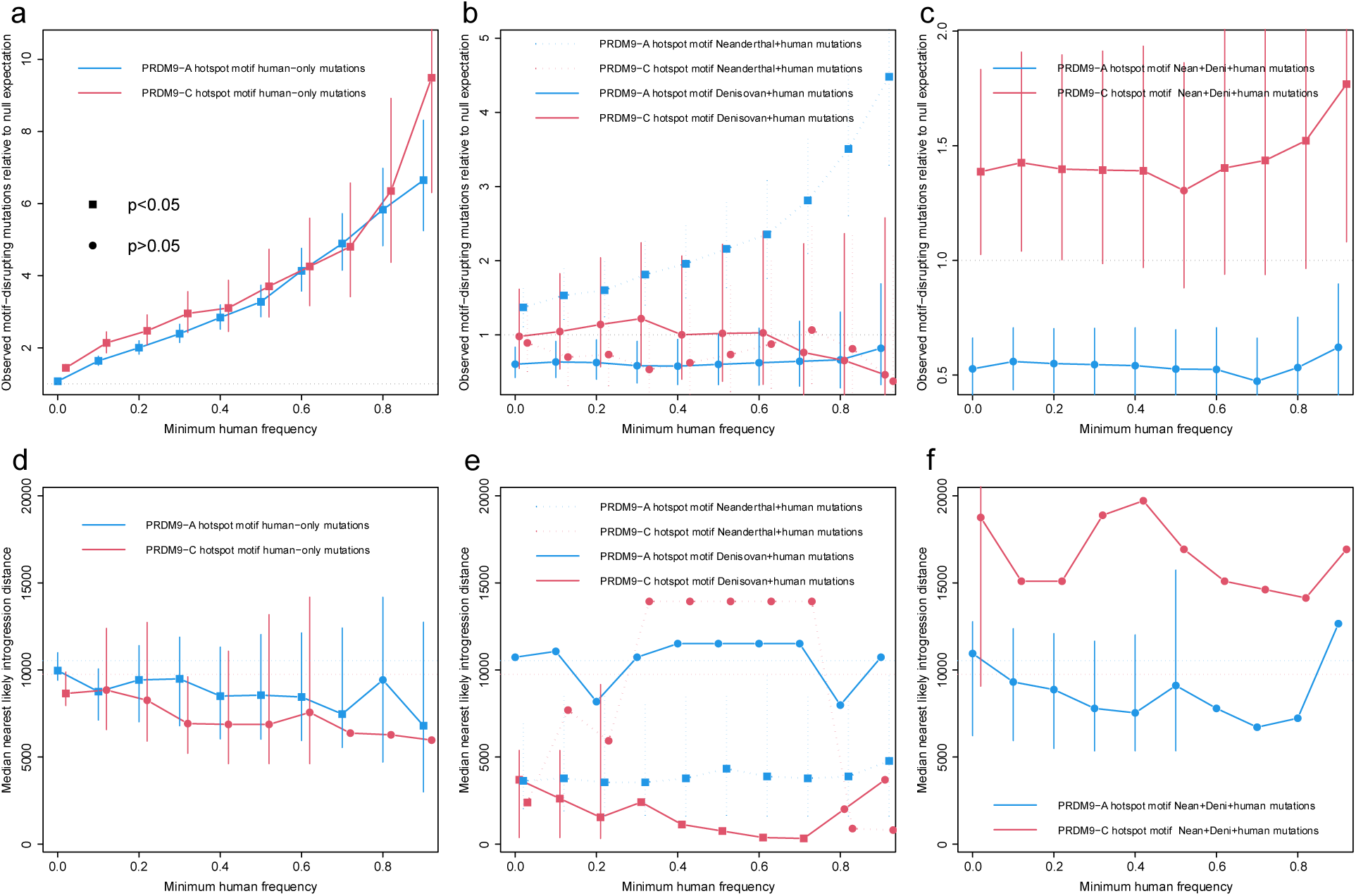
Hotspot death signals in humans and archaic hominins. (a–c) Enrichment of PRDM9 motif-disrupting mutations relative to a local mutation-rate null, stratified by minimum derived allele frequency in humans (Supplementary Note). Mutations are classified by carrier status (a) human-only, (b) Neanderthal–human, Denisovan–human, or (c) shared across all three and by disruption of PRDM9-A or PRDM9-C binding motifs. Points denote observed-to-expected ratios, with error bars indicating 95% confidence intervals; filled symbols indicate nominal significance (*P* < 0.05). A strong enrichment is observed in Neanderthal–human shared mutations at PRDM9-A hotspots and in mutations shared across humans, Neanderthals, and Denisovans at PRDM9-C hotspots. (d–f) Median distance of motif-disrupting mutations to the nearest candidate human-to-Neanderthal introgressed site, stratified and classified as in (a-c) and for the same mutations. Lower values indicate closer proximity to introgressed regions. Dotted horizontal lines: control expectations from nearby non-disrupting mutations. Distinct patterns across mutation categories reveal that A-allele hotspot-disrupting mutations are enriched near introgressed regions, while C-allele mutations are preferentially absent from Neanderthal near introgressed regions, consistent with differential hotspot activity across ancestral populations. P-values and underlying data are provided in Supplementary Table 3c-d.

**Extended Data Figure 10:**
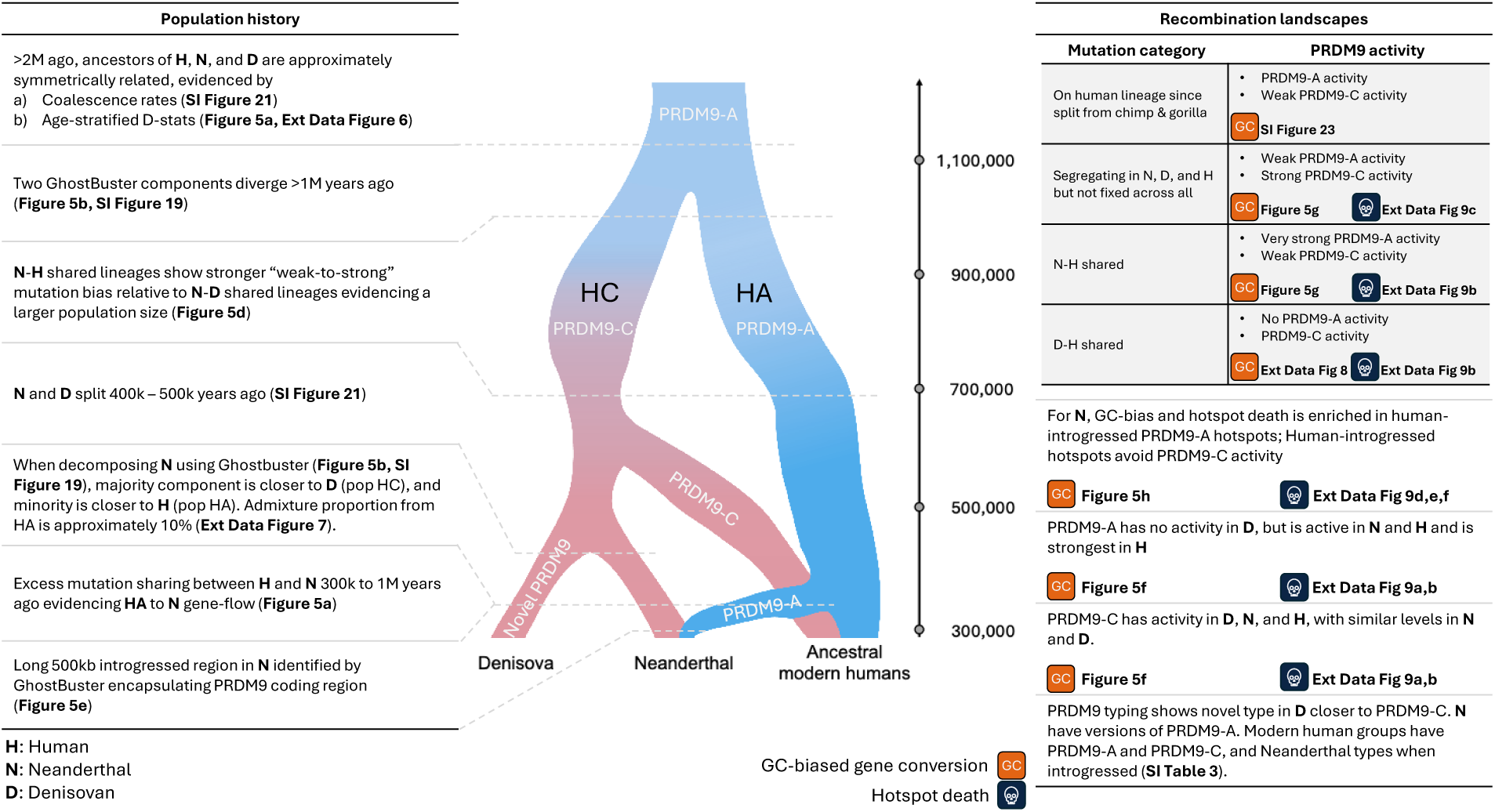
**Overview of the evidence supporting the joint population and recombination history of modern humans, Neanderthals, and Denisovans.**

