## Supplementary Note for "Genome-wide genealogies reveal deep admixtures forming modern humans"

**Supplementary Figures** **46**

**Supplementary Tables** **68**

**References** **70**

### Supplementary Note

### 1 Background

We will first introduce the general concept of genealogies and mixture models before presenting the theoretical framework of GhostBuster.

#### 1.1 The coalescent model

GhostBuster takes genome-wide genealogies as input, which describe how individual genomes share ancestors back in time. The coalescent model defines a distribution on these genealogies, and we briefly introduce the key concepts from coalescent theory that we will use in GhostBuster.

The simplest model for simulating genetic inheritance is the Wright-Fisher model [1, 2], which describes how populations evolve forward in time. It is defined as follows: (i) Each generation has $N$ haploid individuals. (ii) Generations are replaced instantaneously with a new set of $N$ haploid individuals. (iii) Each individual chooses one parent uniformly at random from individuals in the previous generation.

We can now sample individual haplotypes in the most recent generation and trace their ancestors back in time. If we simulate enough generations, they will eventually coalesce into a single ancestor. Supplementary Note Figure **1** shows an example simulation.

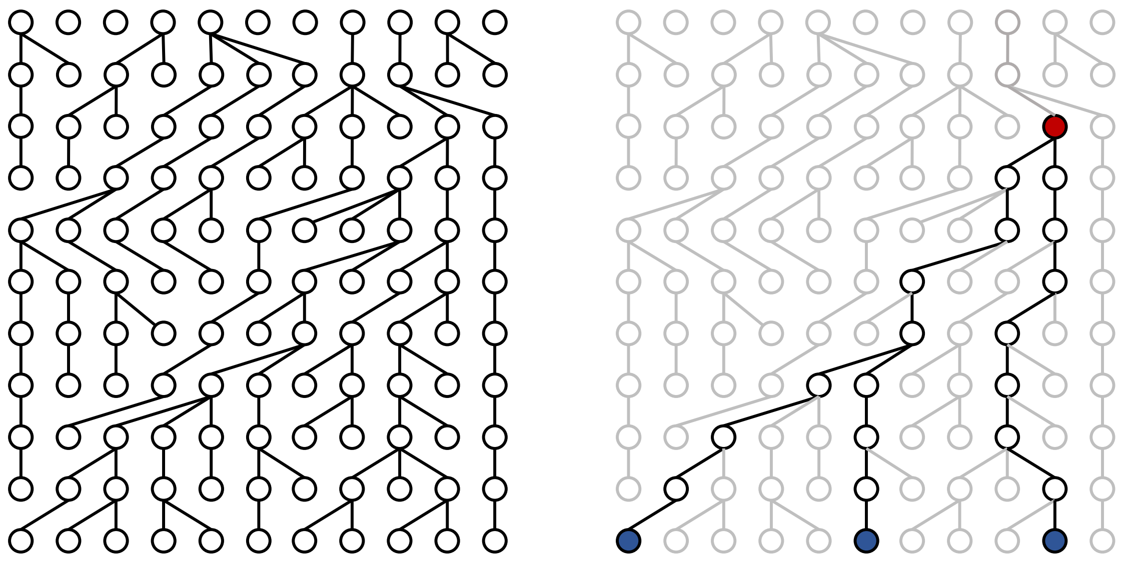

**Supplementary Note Figure 1**: **Example of three lineages coalescing backwards in time in the Wright Fisher model.** Each individual chooses its parent from the set of individuals with replacement until all the lineages coalesce into a single common ancestor. The three samples are coloured blue, and the common ancestor for the three samples is red.

We call the binary tree structure relating haplotypes a genealogy. It is possible to derive the probability distribution of the shape and the most recent common ancestors (TMRCAs) under the Wright-Fisher model.

For simplicity, we assume a constant effective population size $N_e$, although in GhostBuster, $N_e$ is variable. For a pair of haplotypes, the TMRCA is geometrically distributed and takes the form:

$$P(\tau_2 = t) = \left(1 - \frac{1}{N_e}\right)^{t-1} \frac{1}{N_e}, \tag{1}$$

where $\tau_2$ is the time to the most recent common ancestor, and $N_e$ is the effective population size. The cumulative distribution function (cdf) of $\tau_2$ is given by

$$P(\tau_2 \le t) = 1 - \left(1 - \frac{1}{N_e}\right)^{t}. \tag{2}$$

For large population sizes $N_e$, TMRCAs are expected to be sufficiently old so that generations can be effectively modelled in continuous time. Formally, taking $N_e \to \infty$ and setting $T_2 = \frac{\tau_2}{N_e}$ we can approximate the cdf as follows:

$$\begin{aligned} P(T_2 \le t) &= P(\tau_2 \le N_e t) = P(\tau_2 \le \lfloor N_e t \rfloor) \\ &= 1 - \left(1 - \frac{1}{N_e}\right)^{\lfloor N_e t \rfloor} \\ &\to 1 - e^{-t} \text{ as } N_e \to \infty. \end{aligned} \tag{3}$$

Therefore, the TMRCA has an exponential distribution. This is the coalescent model for two samples.

We can extend the coalescent model to $n$ haplotypes instead of a pair. Let us define $\xi(\tau)$ as the number of lineages remaining at $\tau$ generations ago. Going backwards in time with the initialisation $\xi(0) = n$, $\xi(\tau)$ behaves as a Markov process. That is, the distribution of $\xi(\tau + 1)$ only depends on the previous one, $\xi(\tau)$. We can define the transition probability of going from $i$ lineages to $j$ lineages in one generation as:

$$p_{i,j} = P(\xi(\tau + 1) = j \mid \xi(\tau) = i). \tag{4}$$

In any generation, the probability of two lineages choosing the same parent is given by $\frac{\binom{i}{2}}{N_e}$. Therefore, ignoring second-order terms where more than two lineages choose the same parent, the transition matrix is given by:

$$\begin{aligned} p_{i,i} &\approx 1 - \frac{\binom{i}{2}}{N_e}, \\ p_{i,i-1} &\approx \frac{\binom{i}{2}}{N_e}, \\ p_{i,j} &\approx 0 \text{ for any } j < i - 1. \end{aligned} \tag{5}$$

In continuous time, the rate of coalescence therefore depends on the number of lineages present at a given time. Let $T_j = \frac{\tau_j}{N_e}$ be the times to go from $j$ to $j-1$ ancestors (scaled by the population size $N_e$) then according to the coalescent $T_j$ is independent of $T_i$ $(i \neq j)$ and is distributed as follows:

$$T_j \sim exp\left(\binom{j}{2}\right), \; f_j(t) = \binom{j}{2} e^{-\binom{j}{2}t}. \tag{6}$$

In particular, the coalescent for $n$ samples only allows up to one pair of lineages to coalesce at a time, making a binary "coalescent" tree. This now defines a distribution of possible coalescent trees and is known as the coalescent model introduced by three scientists in 1982-1983: Kingman, Hudson, and Tajima [3–7].

We can generalise the above model for time-varying population sizes. Let us define the effective population size at $\tau$ generations ago as $N_e(\tau)$. So long as there exists an $M$ such that $N_e(\tau)/M \in (0, \infty)$, we can rescale discrete generations to continuous time by setting $t = \tau/M$ and taking the limit $M \to \infty$. The distribution of coalescence times is now given by an inhomogeneous exponential distribution, defined by

$$P(T_j > t \mid T_n + T_{n-1} + ... + T_{j+1} = s) = exp\left(-\binom{j}{2} \int_s^{s+t} \frac{1}{\nu(u)} du\right). \tag{7}$$

The above equation simplifies to the standard coalescent with constant population size if we set $\nu(t) \equiv 1$.

##### 1.1.1 Coalescent with recombination

The coalescent model defines a distribution on the shape of genealogies in non-recombining clonal organisms. In diploid sexually reproducing organisms, two homologous chromosomes are condensed into a single mosaic chromosome during meiosis using a process called recombination. A consequence is that the ancestors from whom we inherit our DNA change as we move along the genome.

We can modify the coalescent model to account for recombination as follows: Each pair of lineages cannot only coalesce but also undergo recombination events, which split the ancestry of a single lineage into two ancestral lineages. This coalescent model with recombination [8] creates a more complex ancestral structure than a simple coalescent tree. The resulting genealogy is an Ancestral Recombination Graph (ARG) [9]. An illustrative example of an ARG is depicted in Supplementary Note Figure **2**.

ARGs generate a sequence of coalescent trees that change along the genome (see Supplementary Note Figure **2**b). At any given site, the marginal genealogy is a coalescent tree, and the simple coalescent model without recombination describes its marginal distribution.

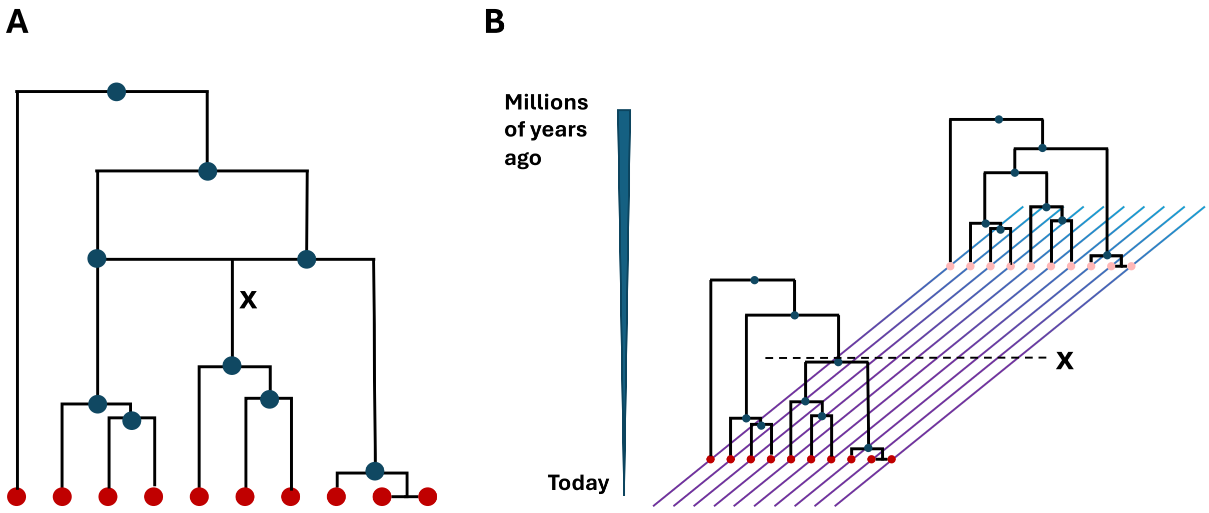

**Supplementary Note Figure 2**: **Example of an ancestral recombination graph (ARG) and the corresponding sequence of trees.** A. In the coalescent with recombination model, the lineages going backwards in time can either coalesce or recombine, which means different portions of the genome have slightly different histories. B. Corresponding sequence of trees from the ARG, **X** denotes the recombination position.

##### 1.1.2 Coalescent with mutations

Given an ARG, we can simulate mutations by dropping mutations onto its branches. Each branch in an ARG persists for a different span $L$ along the genome. We model mutations occurring as a homogeneous Poisson process on each branch, such that the total number of mutation events will follow a Poisson distribution with mean $\mu t L$, where $\mu$ is the mutation rate per base pair per generation, $t$ is the time to the most recent common ancestor in generations.

#### 1.2 Inference of coalescence rates for a pair of lineages

Given a coalescent tree, we detail how we infer a coalescence rate between a pair of lineages. This estimator was derived in [10]. A modified version of this estimator will also appear in GhostBuster.

We assume that the coalescent tree was sampled under the standard coalescent with a time-varying coalescence rate. We infer piecewise constant coalescence rates defined by fixed time intervals or epochs, labelled as $I_1$, $I_2$, ... $I_m$.

Let us say the coalescence rate between a fixed pair of samples is given by $\gamma(e)$, where $e \in 1, \ldots, m$ indexes time epochs. These $\gamma(e)$ are the parameters to be inferred.

In the $\ell$-th tree, the coalescence time is denoted by $t_\ell$. We denote by $e_\ell$ the epoch in which this coalescence event occurs. Therefore $I_{e_l} \leq t_l \leq I_{e_l+1}$.

Extending the coalescent from Equation 7 for piecewise constant coalescence rates, we can write the likelihood of the coalescence time (for a pair of lineages) as:

$$P(t_l) = \gamma(e_l) e^{-\gamma(e_l)(t_l - I_{e_l})} \prod_{e=1}^{e_l} e^{-\gamma(e-1)(I_e - I_{e-1})} \tag{8}$$

For maximum likelihood estimation, we assume the $\mathbf{L}$ trees (or data points) are independent, and so we sum the log-likelihood across trees. We maximise the log-likelihood given by

$$\begin{aligned} \log P(t_l) &= \log \gamma(e_l) - \gamma(e_l)(t_l - I_{e_l}) - \sum_{e=1}^{e_l} \gamma(e-1)(I_e - I_{e-1}), \\ \log P(t) &= \sum_{l=1}^{\mathbf{L}} \left( \log \gamma(e_l) - \gamma(e_l)(t_l - I_{e_l}) - \sum_{e=1}^{e_l} \gamma(e-1)(I_e - I_{e-1}) \right). \end{aligned} \tag{9}$$

We can compute the root of the first derivative to find a closed-form formula for the maximum likelihood estimate of the coalescence rates, which is given by

$$\hat{\gamma}(e) = \frac{n_e}{\sum_{l:e=e_l}(t_l - I_e) + \sum_{l:e<e_l}(I_e - I_{e-1})}. \tag{10}$$

Here, $n_e$ is the number of trees in which the lineages coalesce in the $e$-th epoch. We refer to $n_e$ as the "coalescence count". The denominator $\sum_{l:e=e_l}(t_l - I_e) + \sum_{l:e<e_l}(I_e - I_{e-1})$ is the sum of branch length until the lineage coalesces and is referred to as the "opportunity".

#### 1.3 Introduction to mixture models

Mixture models are probabilistic models that model how subpopulations combine to make up a larger population. They assume that the data is generated from a mixture of distributions, each corresponding to a subpopulation or cluster.

GhostBuster is a mixture model, and as is common with mixture models, we will fit the GhostBuster model using an Expectation-Maximisation (EM) algorithm. Here, we will introduce how an EM algorithm can be used to fit the simplest version of a mixture model.

##### 1.3.1 Gaussian mixture models

One of the simplest and most widely used mixture models is the Gaussian Mixture Model (GMM). The GMM assumes that the data is generated from a mixture of multiple Gaussian distributions, where each observation is drawn from one of the Gaussian components.

Let us consider a dataset $x = (x_1, x_2, \ldots, x_n)$, where $x$ is a vector of $n$ independent observations. For simplicity, assume that these observations are sampled from a mixture of two Gaussian distributions. To model this mixture, we introduce a set of hidden variables $z = (z_1, z_2, \ldots, z_n)$, where each $z_i$ is a categorical variable indicating which Gaussian component generated the corresponding observation $x_i$. Specifically, $z_i = 0$ if $x_i$ is drawn from the first Gaussian component, and $z_i = 1$ if $x_i$ is drawn from the second. The conditional distribution of the random variable $X_i$, given its corresponding hidden variable $Z_i$, is defined as follows:

$$X_i|(Z_i = 0) \sim \mathcal{N}(\mu_1, \Sigma_1),$$
$$X_i|(Z_i = 1) \sim \mathcal{N}(\mu_2, \Sigma_2),$$

where, $\mathcal{N}(\mu_1, \Sigma_1)$ and $\mathcal{N}(\mu_2, \Sigma_2)$ denote Gaussian distributions with means $\mu_1$ and $\mu_2$, and covariance matrices $\Sigma_1$ and $\Sigma_2$, respectively. Additionally, let us assume the prior probabilities of the components are given by $P(Z_i = 0) = \pi_1$ and $P(Z_i = 1) = \pi_2$, where $\pi_2 = 1 - \pi_1$.

The GMM aims to estimate the parameters corresponding to the mixture of two Gaussians, namely $\theta = (\pi_1, \mu_1, \Sigma_1, \mu_2, \Sigma_2)$, by maximising the likelihood of the observed data. However, it is important to note that the vector $z$ is a hidden variable, meaning we do not know which observation corresponds to which cluster. Consequently, we need to maximise the "incomplete-data likelihood," which integrates out the hidden variable $z$:

$$L(\theta; x) = \prod_{i=1}^{n} \Big[\pi_1 \mathcal{N}(x_i|\mu_1, \Sigma_1) + \pi_2 \mathcal{N}(x_i|\mu_2, \Sigma_2)\Big]. \tag{11}$$

Here, $\mathcal{N}(x_i|\mu_j, \Sigma_j)$ represents the Gaussian probability density function for the $j$-th component, evaluated at $x_i$.

Since the likelihood involves a summation inside the product, direct maximisation is challenging. To address this, the Expectation-Maximisation (EM) algorithm is employed [11]. Instead of directly maximising the incomplete-data likelihood, the EM algorithm operates on the "complete-data likelihood," which assumes knowledge of the hidden variables $z$:

$$L(\theta; x, z) = \prod_{i=1}^{n} \prod_{j=1}^{2} \Big[\pi_j \mathcal{N}(x_i|\mu_j, \Sigma_j)\Big]^{\mathbb{1}(z_i = j)} \tag{12}$$

where, $\mathbb{1}(z_i = j)$ is an indicator variable that equals 1 if the observation $x_i$ belongs to component $j$, and 0 otherwise. This formulation represents the likelihood as a product and the log-likelihood as a sum over components and observations, which eventually simplifies the optimisation process.

$$\begin{aligned} \log L(\theta; x, z) &= \sum_{i=1}^{n} \sum_{j=1}^{2} \mathbb{1}(z_i = j) \Big[\log \pi_j + \log \mathcal{N}(x_i|\mu_j, \Sigma_j)\Big] \\ &= \sum_{i=1}^{n} \sum_{j=1}^{2} \mathbb{1}(z_i = j) \Big[\log \pi_j - \frac{1}{2} \log|\Sigma_j| - \frac{1}{2}(x_i - \mu_j)^\top \Sigma_j^{-1}(x_i - \mu_j) - \frac{d}{2}\log(2\pi)\Big]. \end{aligned} \tag{13}$$

The EM algorithm alternates between the E-step, where the expected value of the log-likelihood over the hidden variables $z_i$ is computed given the current parameter estimates (in this example, requiring computing the expected hidden variables themselves because the log-likelihood is linear in these variables), and the M-step, which maximises the expected complete-data log-likelihood with respect to the parameters $\theta$. Through these iterative updates, the EM algorithm refines the parameter estimates, ultimately converging to a local maximum of the incomplete-data likelihood [11] (see Supplementary Note Figure **3**). The EM algorithm for GMM operates as follows:

1. **E-step**: Calculates the expected log-likelihood of the complete data, given the current parameter

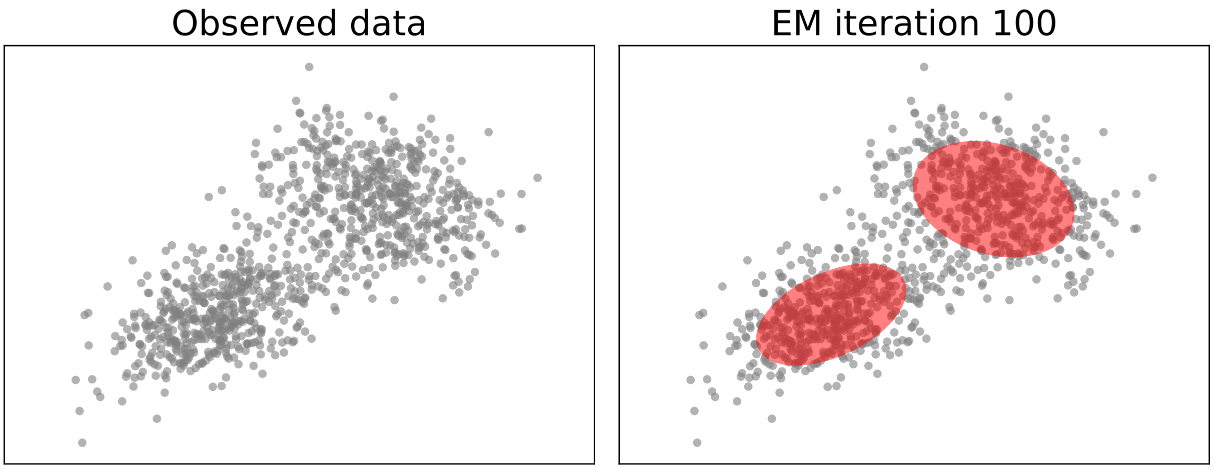

**Supplementary Note Figure 3**: **Gaussian mixture model.** The observed data and inferred clusters using the Gaussian mixture model after 100 EM iterations, the Gaussian mixtures inferred are represented by the red ellipse.

estimates.

$$
\begin{aligned}
&\mathbb{E}_{Z|X=x;\theta^{(t)}}\left[\log L(\theta; X, Z)\right] \\
&= \mathbb{E}_{Z|X=x;\theta^{(t)}}\left[\log \prod_{i=1}^{n} L(\theta; x_i, Z_i)\right] \\
&= \mathbb{E}_{Z|X=x;\theta^{(t)}}\left[\sum_{i=1}^{n} \log L(\theta; x_i, Z_i)\right] \\
&= \sum_{i=1}^{n} \mathbb{E}_{Z_i|X_i=x_i;\theta^{(t)}}\left[\log L(\theta; x_i, Z_i)\right] \\
&= \sum_{i=1}^{n}\sum_{j=1}^{2} P(Z_i = j|X_i = x_i; \theta^{(t)}) \log L(\theta_j; x_i, j) \\
&= \sum_{i=1}^{n}\sum_{j=1}^{2} \gamma_j^{(t)}(x_i)\left[\log \tau_j - \frac{1}{2}\log|\Sigma_j| - \frac{1}{2}(x_i - \mu_j)^\top \Sigma_j^{-1}(x_i - \mu_j) - \frac{d}{2}\log(2\pi)\right].
\end{aligned}
\tag{14}
$$

Here, the log-likelihood of the complete data comes from Equation 13; $\gamma_j(x_i)$ is the posterior probability that the data point $x_i$ was generated by the $j$-th Gaussian distribution. Using Bayes' rule:

$$
\gamma_j^{(t)}(x_i) = \frac{\pi_j^{(t)}\mathcal{N}(x_i|\mu_j^{(t)}, \Sigma_j^{(t)})}{\sum_{j=1}^{2}\pi_j^{(t)}\mathcal{N}(x_i|\mu_j^{(t)}, \Sigma_j^{(t)})}. \tag{15}
$$

2. **M-step**: Maximises the expected log-likelihood, computed in the E-step, with respect to the model

parameters $\theta$. For a GMM, this results in straightforward analytical update equations, as shown below:

$$\pi_j^{(t+1)} = \frac{1}{n}\sum_{i=1}^{n}\gamma_j^{(t)}(x_i)$$
$$\mu_j^{(t+1)} = \frac{\sum_{i=1}^{n}\gamma_j^{(t)}(x_i)x_i}{\sum_{j=1}^{n}\gamma_j^{(t)}(x_i)}$$
$$\Sigma_j^{(t+1)} = \frac{\sum_{i=1}^{n}\gamma_j^{(t)}(x_i)(x_i - \mu_j^{t+1})(x_i - \mu_j^{t+1})^\top}{\sum_{j=1}^{n}\gamma_j^{(t)}(x_i)}. \tag{16}$$

### 2 Theoretical framework for GhostBuster

GhostBuster models the ancestry of individuals as a mixture of source populations. It takes **genome-wide genealogies** as input data and fits a **mixture model** to the observed coalescences with other individuals back in time.

#### 2.1 Mixture model for the coalescent

We aim to extend the basic example of mixture models to genealogies by specifically modelling coalescence events on a given target lineage as coming from a mixture distribution of ancestry components. As input, our method uses, for each site, information about which individuals the target sample coalesces with and the corresponding coalescence times as we go backwards in time. Our goal is to model this as a mixture distribution, where each site's coalescence history is drawn from one ancestry component. In essence, we therefore cluster the genome into different ancestries, each producing different coalescence histories. The output of our model is an inferred per-component coalescence rate matrix, which specifies how the target individual coalesces with each reference population across the components of the mixture, admixture proportions of each ancestry, and local ancestry along the genome.

Our method leverages the fact that admixture can be characterised by coalescence rates with a set of reference populations. To illustrate this, consider the example of Denisovan admixture in Papuans. It is well known that Papuans carry up to 5% Denisovan ancestry in their genomes contributed around 40,000 years ago [12]. When a genomic segment is inherited from Denisovans, this archaic ancestry has different coalescence rates to reference populations compared to the majority of Papuan ancestry: regions with Denisovan ancestry exhibit more frequent coalescence events with Denisovans recently (last 40,000 − 700,000 years), but do not tend to coalesce with modern human lineages until the time of the Denisovan-human split, estimated around 700,000 years ago [12]. GhostBuster will leverage this clear difference in the coalescence history between archaic and non-archaic genomic regions in Papuans. Importantly, our method does not require explicit source groups or unadmixed outgroups to infer an admixture event. In the Papuan example, Denisovan ancestry not only exhibits quicker coalescence with Denisovans but also shows slower coalescence with other Papuans and modern human groups. We can leverage these differences in coalescence rates with modern human groups alone to characterise the admixture, enabling us to infer admixture without relying on explicit source groups or additional assumptions about outgroups.

##### 2.1.1 The generative model

To formalise the model, we first define the probabilistic process that gives rise to the observed data, which in our case are coalescent trees along the genome. Our approach is based on a structured generative model where, given model parameters - namely, the global ancestry proportions ($\pi$), the admixture time ($\lambda$), and the coalescence rates with reference groups ($\gamma$) - we can generate the coalescent history of a target individual at each position under these assumptions.

1. Global ancestry proportions ($\pi$): The target lineage is assumed to originate from a mixture of ancestries, where the global ancestry proportions $\pi$ describe the expected fraction of the genome inherited from each ancestral source. Therefore, $\pi$ is a vector of length equal to the number of ancestries.

2. Admixture time ($\lambda$): The admixture event is assumed to have occurred $\lambda$ generations in the past. Therefore, $\lambda$ is a scalar.

3. Coalescence rates ($\gamma$): For each ancestry, the coalescence rates with reference populations are determined by the matrix $\gamma$, which encodes the instantaneous rate at which the target lineage coalesces with lineages belonging to each reference group. Therefore, $\gamma$ is a 3D-tensor of dimension equal to the number of ancestries $\times$ number of reference groups $\times$ number of time epochs. The ancestry the target

individual belongs to can change along the genome, and we will therefore refer to the local ancestry of a target individual.

The observed data under the model consist of the coalescence genealogies of the target individual with reference populations. The generative process for producing the observed genealogies is as follows:

1. **Sample local ancestry**: The local ancestry states along the genome are sampled according to a Markov model with global ancestry proportions $\pi$ and transition probabilities determined by the admixture time $\lambda$. More details about the Markov model can be found in section 2.5.
2. **Thread the target sample into the genealogy**: Given local ancestry, the target sample is integrated into an existing genealogy of reference samples, conditional on the corresponding coalescence rates of that ancestry. The coalescence rates provide the rate at which we coalesce with lineages belonging to sampled reference populations; however, in practice, ancestral lineages have a mixture of descendants from different populations. We therefore define a probability of coalescing with a particular lineage depending on the descendant composition of the coalescing lineage. Further details on the likelihood associated with coalescence with a lineage can be found in the next subsection.
3. **Construct the final genealogy**: The resulting genealogy consists of the original reference samples plus the target sample at each genomic position.

###### 2.1.2 Coalescence rates with a 'mixed' lineage

As demonstrated in Section 1.1, the properties of the coalescent provide a closed-form solution for the Maximum Likelihood Estimate (MLE) of coalescence rates when only one reference population is considered. However, when dealing with lineages whose descendants originate from a mixture of populations, computing the likelihood becomes more complex.

Such mixed lineages arise primarily from two factors: (1) historical gene-flow or coalescences predating population splits and (2) uncertainty in genealogy inference from genetic variation data. To illustrate this, we revisit the case of Denisovan admixture in Papuans. Suppose we analyse a Papuan individual ($\approx 5\%$ Denisovan ancestry) alongside other Papuans (who also carry some Denisovan ancestry) and archaic Denisovans as reference populations. In this scenario, the target Papuan individual may carry local ancestry derived from Denisovans but still coalesce with lineages containing both Papuan and Denisovan samples. This could be due to the merging of Denisovan and human populations over 700,000 years ago, Denisovan admixture in Papuan reference samples more recently, or simply uncertainty in tree topology and dating estimates in genealogy inference methods, leading to noisier split times.

To compute the likelihood of a lineage coalescing with a mixed lineage, we leverage the descendant composition of lineages - i.e., the reference population assignments at the leaves of the tree - to inform the coalescence rate with the target. This likelihood is calculated in two steps: (1) We first determine the population of the lineage and (2) sample the coalescence time conditioned on this information. Specifically, we assume that the population label of a lineage is drawn uniformly from the set of its descendants. For instance, if a lineage has five Papuan and one Denisovan descendant, it is assigned to the Papuan population with probability $\frac{5}{6}$ and to the Denisovan population with probability $\frac{1}{6}$. Given the population label, the coalescence time with the target follows an exponential distribution, with a rate determined by the coalescence rate between the target and the sampled population. This approach integrates naturally with the EM algorithm introduced later. By treating the lineage label as a latent variable, we effectively "complete" the data, enabling analytical updates at each iteration of the algorithm.

Beyond its computational efficiency, we believe this approach is also one of the simplest ways to model 'mixed' lineages. Instead of relying on knowledge of the population history in the reference population, our method samples descendants uniformly. This increases robustness to errors in genealogy inference by preventing abrupt lineage label switches. Additionally, in a clean split model, when populations merge, their coalescence rates become equal, rendering the choice of averaging inconsequential. Finally, under the assumption of approximately equal sample sizes and genetic drift across reference populations — and without prior knowledge of the genetic history of reference samples - this weighting provides an unbiased likelihood of coalescence with a given lineage. Future work can relax some of these assumptions with improved weighting schemes.

This approach of randomly sampling a descendant and computing the pairwise likelihood leads to a mixture of exponentials weighted by the descendants. It is important to note that this differs from a simpler composite likelihood approach, which assumes pairwise independence between lineages and, by not conditioning on the inferred tree topology, overcounts non-independent events, leading to overconfident estimates and reduced statistical power.

Additionally, the maximum likelihood estimation (MLE) of coalescence rates derived in Section 1.1 assumes independence of trees along the genome, which is an unrealistic assumption. To address this, we employ a Hidden Markov Model (HMM) to account for ancestry switches along the genome, modelling each location as Markovian. This approach relaxes the independence assumption and improves the robustness and accuracy of our likelihood estimation.

##### 2.1.3 Likelihood of a single coalescence event

Under our model, we derive the likelihood of a single coalescence event involving a lineage whose descendants are drawn from a mixture of populations. Let the coalescing clade or lineage have $n_g$ descendants from population $g$, with the total number of descendants across $R$ reference populations given by $\sum_{g=1}^{R} n_g$. The probability density for observing a coalescence event at time $t$, given the matrix of coalescence rates per group per epoch $\gamma$, is expressed as:

$$P(t \mid \gamma) = \frac{\sum_{g=1}^{R} \gamma_g(m) n_g}{\sum_{g=1}^{R} n_g} e^{-\sum_{g=1}^{R} \gamma_g \cdot O_g(t)}, \tag{17}$$

where, $n_g$ is the number of descendants in group $g$, $\gamma_g(m)$ is the coalescence rate of group $g$ during epoch $m$, and $\gamma_g \cdot O_g(t)$ is the dot product (over time epochs) of the coalescence rates $\gamma_g$ for group $g$ and opportunity $O_g(t)$, defined as the sum of branch lengths from the previous coalescence time to $t$ for all group $g$ individuals.

This weighting is equivalent to randomly sampling a descendant from the coalescing lineage and computing the pairwise coalescence likelihoods. Let $q$ be a categorical random variable that indicates which individual coalesces with the target. The probability of selecting the $k$-th descendant is given by:

$$P(q = k \mid \gamma) = \frac{1}{\sum_{g=1}^{R} n_g}, \tag{18}$$

where $n_g$ is the number of descendants in group $g$. The joint probability of observing a coalescence event at time $t$ and the descendant $q = k$, assuming conditional independence between $t$ and $q$, is expressed as:

$$\begin{aligned} P(t, q = k \mid \gamma) &= P(t \mid q = k, \gamma) P(q = k \mid \gamma) \\ &= \gamma_{g(k)}(m) e^{-\sum_{g=1}^{R} \gamma_g \cdot O_g(t)} \frac{1}{\sum_{g=1}^{R} n_g}, \end{aligned} \tag{19}$$

where $P(t \mid q = k, \gamma)$ is derived from the pairwise coalescence likelihood. Summing over all possible descendants in the coalescing clade, we obtain the following expression for the likelihood of observing a coalescence event at time $t$ with this lineage:

$$\begin{aligned} P(t \mid \gamma) &= \sum_{k \in \text{Clade}} P(t, q = k \mid \gamma) \\ &= \frac{\sum_{g=1}^{R} \gamma_g(m) n_g}{\sum_{g=1}^{R} n_g} e^{-\sum_{g=1}^{R} \gamma_g \cdot O_g(t)}. \end{aligned} \tag{20}$$

This matches the expression in Equation 17.

#### 2.2 EM derivation: terminology

We aim to fit a mixture model to cluster the genome based on variations in coalescence rates. To reduce computational costs, we divide the genome into $T$ small grids and focus on inferring local ancestry at these grid points, rather than at every individual location on the genome (by default, grid points are equally spaced 10kb or 0.05cM apart). Additionally, we approximate coalescence rates as piecewise constant functions over time, defined by $E$ time epochs that are equally spaced on a logarithmic scale [13]. For the derivation of the EM algorithm, we use the following terminology:

##### 2.2.1 Fixed constants used in derivation

1. $C$: Total number of coalescence events between the target lineage and reference samples across the entire genome.
2. $T$: Total number of grid points.
3. $A$: Number of clusters assumed for the target lineage.
4. $E$: Total number of time epochs in which the coalescence rates are assumed to be piecewise constant (by default, $E = 20$).
5. $N$: Total number of reference individuals in the genealogy.
6. $R$: Total number of reference groups in the genealogy.
7. $N_{li}$: Number of reference individuals subtending the lineage which coalesces with the target lineage in the $i$-th coalescence event at the $l$-th grid position.
8. $N_c$: Number of reference individuals subtending the lineage corresponding to the $c$-th coalescence event.

##### 2.2.2 Subscripts/Superscripts used in derivation

1. $l$: Index of the grid position.
2. $i$: Index of the coalescence event involving the target in the local tree.
3. $g$: Predefined reference group (e.g., Papuans, Denisovans, etc.).
4. $j$: Index of the cluster or component in the mixture model.
5. $m$: Index of the time epoch, measured backwards in time.
6. $c$: Index of the coalescence event involving the target in the entire genealogy.
7. $v$: $v$-th EM iteration.

##### 2.2.3 Input data

1. $t_{li}, t_c \in \mathbb{R}^+$: Coalescence time, where $t_{li}$ denotes the time (measured relative to the previous coalescence event) for the $i$-th coalescence event at the $l$-th grid position, and $t_c$ represents the time for the $c$-th coalescence event.
2. $n_{lig}, n_{cg} \in \mathbb{Z}^+$: Number of individuals from group $g$ involved in a coalescence event, where $n_{lig}$ represents the number of individuals from group $g$ participating in the $i$-th coalescence event at the $l$-th grid position, and $n_{cg}$ denotes the number of individuals from group $g$ involved in the $c$-th coalescence event.
3. $Q_l \in \mathbb{Z}^+$: Total number of coalescence events between the target lineage and reference samples within the tree corresponding to the $l$-th grid position.
4. $e_{li}, e_c \in \{1, 2, 3, \ldots, E\}$: Time epoch during which a coalescence event occurs, where $e_{li}$ denotes the epoch of the $i$-th coalescence event at the $l$-th grid position, and $e_c$ represents the epoch of the $c$-th coalescence event.
5. $r_l \in \mathbb{R}^+$: Genetic distance between the $(l-1)$-th and $l$-th grid positions (in Morgans).

##### 2.2.4 Mixture model parameters

1. $\gamma \in \mathbb{R}_+^{A\times R\times E}$: Coalescence rate matrix, where $\gamma_{jg}(m)$ represents the coalescence rate between reference group $g$ and the target in the $j$-th mixture component during epoch $m$.
2. $\pi \in [0,1]^A$: Cluster proportions, where $\pi_j = P(z_l = j)$ denotes the proportion of the $j$-th mixture component. Additionally, $0 \leq \pi_j \leq 1, \quad \sum_{j=1}^{A} \pi_j = 1$.
3. $\lambda \in \mathbb{R}^+$: Admixture time, measured in generations.

##### 2.2.5 Hidden variables in the EM

1. $z \in \{1, 2, \ldots, A\}^T$: Local ancestry vector, where $z_l$ represents the local ancestry of the target at the $l$-th grid position, $z_c$ denotes the same for the $c$-th coalescence event.
2. $q \in \{1, 2, \ldots, N\}^C$: Reference individual indices, where $q_{li}$ denotes the index of the reference individual that coalesces in the $i$-th coalescence event at the $l$-th grid position, $q_c$ denotes the same for $c$-th coalescence event.
3. $q' \in \{1, 2, \ldots, R\}^C$: Reference individual groups, where $q'_{li}$ denotes the population group of the reference individual coalescing in the $i$-th coalescence event at the $l$-th grid position, $q'_c$ denotes the same for $c$-th coalescence event.
4. $\omega_l \in \mathbb{R}^+$: Number of recombination events since admixture, occurring between the $(l-1)$-th and $l$-th grid positions.

#### 2.3 EM derivation: the likelihood

##### 2.3.1 The "incomplete-data likelihood"

We aim to formalise the likelihood for the mixture model described in Section 2.1. Following the approach of the EM algorithm for GMMs outlined in Section 1.3, we first define the "incomplete-data likelihood," which is based solely on the directly observable quantities. These include: (1) The vector of coalescence times for each event between the target lineage and reference samples, denoted as $t = (t_1, t_2, \ldots, t_C)$. (2) The corresponding vector of time epochs, $e = (e_1, e_2, \ldots, e_C)$. (3) The clade distribution vector, $n_g = (n_{1g}, n_{2g}, \ldots, n_{Cg})$, which records the number of individuals from group $g$ corresponding to each coalescence event. The incomplete-data likelihood, which incorporates these observable quantities, is expressed as:

$$P(t|\gamma, \pi, \lambda) = \sum_z P(t|z, \gamma, \pi, \lambda) P(z|\gamma, \pi, \lambda). \tag{21}$$

Here, $z$ is a hidden categorical random variable that determines, for each coalescence event involving the target individual, which ancestry component it occurred in. The parameters to be inferred include $\gamma$, $\pi$, and $\lambda$: $\gamma$ represents the coalescence rate matrix, $\pi$ denotes the mixture proportions, and $\lambda$ is the admixture time. See section 2.2 for their definitions. To compute the incomplete-data likelihood, we sum over all possible configurations of the hidden variable $z$ using $\sum_z$. Notably, given the value of $z$, the conditional probability $P(t|z, \gamma, \pi, \lambda)$ can be factorised across coalescence events.

$$P(t|\gamma, \pi, \lambda) = \sum_z \left( \prod_{c=1}^{C} P(t_c|z_c, \gamma, \pi, \lambda) \right) P(z|\gamma, \pi, \lambda). \tag{22}$$

We can now utilise our model as described in Section 2.1 and Equation 17 to get $P(t_c|z_c, \gamma, \pi, \lambda)$:

$$P(t|\gamma, \pi, \lambda) = \sum_{z:z_c=j} \left( \prod_{c=1}^{C} \frac{\sum_{g=1}^{R} \gamma_{jg}(e_c) n_{cg}}{\sum_{g=1}^{R} n_{cg}} e^{-\sum_{g=1}^{R} \gamma_{jg} \cdot O_g(t)} \right) P(z|\gamma, \pi, \lambda). \tag{23}$$

Similar to equation 17, $\gamma_{jg} \cdot O_g(t)$ is the dot product (over time epochs) between the coalescence rates of the $j$-th component, $g$-th reference group and Opportunity of the $g$-th group.

##### 2.3.2 The "complete-data likelihood"

As with the GMM, the incomplete-data likelihood in Equation 23 is not directly optimisable due to its complex form. To address this, we employ the EM algorithm, which utilises the complete-data likelihood to maximise the incomplete-data likelihood. This assumes access to complete information, including both observable quantities and hidden variables. In our case, hidden variables to facilitate computations in the EM algorithm include: (1) $z = (z_1, z_2, \ldots, z_C)$, which records the mixture component assignment for the $c$-th coalescence event. (2) $q = (q_1, q_2, \ldots, q_C)$, which identifies the reference individual involved in the $c$-th coalescence event. If local ancestry is computed using an HMM, we introduce another hidden variable (3) $\omega = (\omega_1, \omega_2, \ldots, \omega_T)$, which identifies the number of recombination events (and hence possible ancestry change) since admixture between two genomic positions ($T$ is the total number of genomic positions considered).

The complete-data likelihood represents the joint probability of observing the coalescence times $t$, along with the hidden variables $z$ and $q$, given the model parameters $\gamma$, $\pi$, and $\lambda$. We express this likelihood as the product of probabilities for each coalescence event, where the product is across all coalescence events (relating the target) $c$, clusters $j$, reference individuals $k$, and time epochs $m$:

$$
\begin{aligned}
P(t, z, q|\gamma, \pi, \lambda) &= P(z|\pi, \gamma, \lambda) P(t, q|z, \pi, \gamma, \lambda) \\
&= P(z|\pi, \lambda) \prod_{c=1}^{C} P(t_c, q_c|z_c, \gamma) \\
&= \underbrace{P(z|\pi, \lambda)}_{\text{Term 1}} \prod_{c=1}^{C} \prod_{j=1}^{A} \prod_{k=1}^{N_c} \prod_{m=1}^{E} \underbrace{P(t_c, q_c = k|z_c = j, \gamma)^{\mathbb{1}_{\{z_c=j\}} \mathbb{1}_{\{q_c=k\}} \mathbb{1}_{\{e_c=m\}}}}_{\text{Term 2}}.
\end{aligned} \tag{24}
$$

Here, $C$ represents the total number of coalescence events involving the target individual, $A$ is the number of assumed clusters or components, $N_c$ denotes the number of reference individuals corresponding to the $c$-th coalescence event, and $E$ is the total number of time epochs. The indicator functions $\mathbb{1}_{\{z_c=j\}}$, $\mathbb{1}_{\{q_c=k\}}$, and $\mathbb{1}_{\{e_c=m\}}$ ensure that probability contributions are assigned to the appropriate cluster, reference individual, and time epoch, respectively. The complete-data likelihood is divided into two terms. The first term depends on the cluster proportions $\pi$ and the admixture time $\lambda$, while the second term depends on the coalescence rates $\gamma$. Notably, the second term simplifies to a product over coalescence events due to the assumption of conditional independence of coalescence times, $t_c$, given the hidden variable $z_c$.

Coalescence events in an ARG can span more than one tree, particularly for nodes lower in the tree, which are less likely to be affected by recombination and thus persist longer along the genome. The likelihood in Equation 24 can be reformulated as a product over genomic positions. To achieve this, we propose a windowing approach that partitions the genome into grids and appropriately scales the likelihood based on the persistence of a coalescence event along the genome. This approach enables us to maintain the structure of Equation 24 while transitioning from a formulation based on individual coalescence events to one that operates along genomic segments:

$$
\begin{aligned}
&P(t, z, q|\gamma, \pi, \lambda) = \\
&= P(z|\pi, \lambda) \prod_{l=1}^{T} \prod_{j=1}^{A} \prod_{c=1}^{C} \prod_{k=1}^{N_c} \prod_{m=1}^{E} \left( P(t_c, q_c = k|z_c = j, \gamma)^{\mathbb{1}_{\{z_c=j\}} \mathbb{1}_{\{q_c=k\}} \mathbb{1}_{\{e_c=m\}}} \right)^{\frac{\mathbb{1}_{\{w_c=l\}}}{K_c}}
\end{aligned} \tag{25}
$$

where, $\mathbb{1}_{\{w_c=l\}}$ is an indicator random variable that identifies whether the $l$-th grid point overlaps with the node corresponding to the $c$-th coalescence event, and $K_c$ represents the total number of grid points spanned by the node associated with the $c$-th coalescence event (referred to as node persistence). The likelihood in Equation 25 is scaled by $\frac{1}{K_c}$ to ensure that, when multiplied over all grids, it remains equivalent to the original likelihood in Equation 24. The likelihood in Equation 25 can be simplified further by considering only the coalescence events in the tree that overlap with the $l$-th grid position.

$$P(t, z, q|\gamma, \pi, \lambda) =$$
$$= P(z|\pi, \lambda) \prod_{l=1}^{T} \prod_{j=1}^{A} \prod_{i=1}^{Q_l} \prod_{k=1}^{N_{li}} \prod_{m=1}^{E} \left( P(t_{li}, q_{li} = k|z_{li} = j, \gamma)^{\mathbb{1}\{z_{li}=j\} \mathbb{1}\{q_{li}=k\} \mathbb{1}\{e_{li}=m\}} \right)^{\frac{1}{K_{li}}} \tag{26}$$

where, instead of taking the product over all coalescence events in the ARG, we consider only the coalescence events within the local tree at the $l$-th grid position. Consequently, coalescence events are indexed by the grid position $l$ and the order of the coalescence event within the tree, denoted as $i$. Here, $Q_l$ represents the total number of coalescence events between the target lineage and reference samples within the tree corresponding to the $l$-th grid position. Further details about the calculation of node persistence, $K_{li}$, can be found in section 3.3.

###### 2.3.3 An approximation to the "true" likelihood

The hidden variable $z_{li}$ in Equation 26 is indexed over coalescence events in the ARG. This indexing adds significant complexity to the likelihood computation and reduces its interpretability, as the hidden variables do not correspond directly to the local ancestry of the target individual at specific genomic positions. Instead, they remain tied to individual coalescence events, making it challenging to efficiently model ancestry transitions along the genome. To mitigate this, we introduce an approximation to the likelihood in Equation 26. Specifically, we replace $z_{li}$, which represents the cluster assignment for the $i$-th coalescence event at the $l$-th grid position, with a single cluster assignment $z_l$ for the entire grid position $l$. This approximation assumes that all coalescence events occurring at a given genomic position share the same cluster assignment. By reducing the number of hidden variables indexed over coalescence events, this approximation significantly simplifies the likelihood computation and makes it more interpretable.

$$P(t, z, q|\gamma, \pi, \lambda) \approx$$
$$\approx P(z|\pi, \lambda) \prod_{l=1}^{T} \prod_{j=1}^{A} \prod_{i=1}^{Q_l} \prod_{k=1}^{N_{li}} \prod_{m=1}^{E} \left( P(t_{li}, q_{li} = k|z_{l} = j, \gamma)^{\mathbb{1}\{z_{l}=j\} \mathbb{1}\{q_{li}=k\} \mathbb{1}\{e_{li}=m\}} \right)^{\frac{1}{K_{li}}} . \tag{27}$$

While this approximation is computationally efficient (as we show later in section 2.5 due to HMMs), it assumes that all coalescence events at a given position share the same ancestry cluster. This simplification may not fully capture the complexity of human demographic history, which involves multiple admixture and migration events. Retaining coalescence event-level indexing for $z$ could provide a richer model that better distinguishes older from recent ancestry shifts. However, we believe that this comes with significant computational challenges. Future work could explore this direction to achieve a more fine-grained representation of human ancestry, capturing the sequential impact of migration waves across time.

###### 2.3.4 Accounting for uncertain coalescence events

Genealogies inferred from genetic variation data come with the added complexity of uncertainty, as they are not always entirely accurate. The support for a particular node in the tree is stronger when there is a mutation mapped just above that node. To account for this uncertainty, we scale the likelihood by the confidence in each coalescence event. Specifically, as we move backwards in time along the target lineage, we count the number of coalescence events that occur before encountering a coalescence event supported by a mutation. All coalescence events between successive mutations are then weighted based on the number of such events. This adjustment is ad hoc, but it improves the robustness of our method to errors in inferred genealogies, as demonstrated in empirical analyses. Overall, this results in scaling the likelihood in Equation 27 by an additional term, $H_{li}$, which calculates the effective contribution of a coalescence event to the likelihood depending on its certainty. The updated likelihood is now given as:

$$
\begin{aligned}
&P(t,z,q|\gamma,\pi,\lambda) \approx \\
&\approx \underbrace{P(z|\pi,\lambda)}_{\text{Term 1}} \prod_{l=1}^{T}\prod_{j=1}^{A}\prod_{i=1}^{Q_l}\prod_{k=1}^{N_{li}}\prod_{m=1}^{E} \underbrace{\left(P(t_{li},q_{li}=k|z_l=j,\gamma)^{\mathbb{1}_{\{z_l=j\}}\mathbb{1}_{\{q_{li}=k\}}\mathbb{1}_{\{e_{li}=m\}}}\right)^{\frac{1}{B_{li}}}}_{\text{Term 2}}. \qquad (28)
\end{aligned}
$$

Here, $B_{li} = H_{li}K_{li}$ represents the effective scaling factor that accounts for both node persistence and the certainty of coalescence events in the inferred genealogies. Further details on the calculation of this weighting can be found in the subsequent sections 3.3 and 3.4.

As discussed earlier, Term 1 and Term 2 are handled separately. We use the expectation-maximisation (EM) algorithm to optimise Term 1 for estimating admixture time $\lambda$ and cluster proportions $\pi$ and Term 2 for estimating coalescence rates $\gamma$. In the following sections, we derive the EM updates for Term 1 (see Section 2.5) and Term 2 (see Section 2.4) independently.

#### 2.4 EM derivation: inference of coalescence rates

We write the likelihood Term 2 in Equation 28, which only depends on coalescence rates as follows:

$$
P(t,q|z,\gamma) = \prod_{l=1}^{T}\prod_{j=1}^{A}\prod_{i=1}^{Q_l}\prod_{k=1}^{N_{li}}\prod_{m=1}^{E} P(t_{li},q_{li}=k|z_l=j,\gamma)^{\frac{\mathbb{1}_{\{z_l=j\}}\mathbb{1}_{\{q_{li}=k\}}\mathbb{1}_{\{e_{li}=m\}}}{B_{li}}} \qquad (29)
$$

where, the indicator random variables $\mathbb{1}_{\{z_l=j\}}$, $\mathbb{1}_{\{e_{li}=m\}}$ and, $\mathbb{1}_{\{q_{li}=k\}}$ are present to simplify the equation when writing it down for all the trees and coalescence events in that tree. The log-likelihood is given by:

$$
\log P(t,q|z,\gamma) = \sum_{l=1}^{T}\sum_{j=1}^{A}\sum_{i=1}^{Q_l}\sum_{k=1}^{N_{li}}\sum_{m=1}^{E} \frac{\mathbb{1}_{\{q_{li}=k\}}\mathbb{1}_{\{z_l=j\}}\mathbb{1}_{\{e_{li}=m\}}\log P(t_{li},q_{li}=k|z_l=j,\gamma)}{B_{li}}. \qquad (30)
$$

We can simplify the likelihood by uniformly sampling a descendant from the coalescing lineage and considering its pairwise coalescence likelihood. See Section 2.1.3 for a detailed explanation and the underlying motivation for this model:

$$
\begin{aligned}
P(t_{li},q_{li}=k \mid z_l=j,\gamma) &= P(t_{li} \mid \gamma, z_l=j, q_{li}=k)P(q_{li}=k \mid z_l=j,\gamma) \\
P(t_{li},q_{li}=k \mid z_l=j,\gamma) &= \gamma_{jg(k)}(m)e^{-\sum_{g=1}^{R}\gamma_{jg}\cdot O_{lg}[i]}\frac{1}{\sum_{g=1}^{R} n_{lig}} \qquad (31)
\end{aligned}
$$

where $g(k)$ denotes the group assignment (among $R$ groups) of the $k$-th individual in the reference panel, $n_{lig}$ represents the number of group $g$ individuals coalescing in the $i$-th coalescence event at position $l$, and $O_{lg}[i]$ is the sum of branch lengths from the $(i-1)$-th to the $i$-th coalescence event corresponding to all group $g$ individuals. Overall, $P(t_{li},q_{li}=k \mid z_l=j,\gamma)$ can be simplified by focusing only on the terms dependent on the cluster membership $j$, as follows:

$$
\begin{aligned}
P(t_{li},q_{li}=k|\gamma,z_l=j) &= \frac{\gamma_{jg(k)}(m)e^{-\sum_{g=1}^{R}\gamma_{jg}\cdot O_{lg}[i]}}{\sum_{g=1}^{R} n_{lig}} \\
&\propto \gamma_{jg(k)}(m) \times e^{-\sum_{g=1}^{R}\gamma_{jg}\cdot O_{lg}[i]}. \qquad (32)
\end{aligned}
$$

In the following sections, we derive the E-step and M-step for the likelihood presented in Equation 29.

##### 2.4.1 E-step

In the E-step, we calculate the expected log-likelihood, where the expectation is with respect to the hidden variables conditioned on the data and previous parameter estimates. We therefore proceed by stating the expected log-likelihood in our case:

$$\mathbb{E}_{z,q|\gamma^{\upsilon-1},\lambda^{\upsilon-1},\pi^{\upsilon-1},t}\Big(\sum_{l=1}^{T}\sum_{j=1}^{A}\sum_{i=1}^{Q_l}\sum_{k=1}^{N_{li}}\sum_{m=1}^{E}\frac{\mathbb{1}_{\{q_{li}=k\}}\mathbb{1}_{\{z_l=j\}}\mathbb{1}_{\{e_{li}=m\}}\log P(t_{li},q_{li}=k|z_l=j,\gamma)}{B_{li}}\Big). \tag{33}$$

We start simplifying the expected log-likelihood by taking the expectation inside the summations, and transforming the expected value of indicator random variables as probabilities.

$$=\sum_{l=1}^{T}\sum_{j=1}^{A}\sum_{i=1}^{Q_l}\sum_{k=1}^{N_{li}}\sum_{m=1}^{E}\mathbb{1}_{\{e_{li}=m\}}P(q_{li}=k,z_l=j|\gamma^{\upsilon-1},\lambda^{\upsilon-1},\pi^{\upsilon-1},t)\frac{\log(P(t_{li},q_{li}=k|z_l=j,\gamma))}{B_{li}}. \tag{34}$$

Simplifying further and substituting the value of $\log(P(t_{li},q_{li}=k|z_l=j,\gamma))$ from Equation 32. Note that we omit the constants of proportionality as they do not depend on the coalescence rates and thus will not affect the maximisation in the M-step.

$$\begin{aligned}
&=\sum_{l=1}^{T}\sum_{j=1}^{A}\sum_{i=1}^{Q_l}\sum_{k=1}^{N_{li}}\sum_{m=1}^{E}\Bigg(P(q_{li}=k|\gamma^{\upsilon-1},t,z_l=j)P(z_l=j|t,\gamma^{\upsilon-1},\lambda^{\upsilon-1},\pi^{\upsilon-1})\\
&\quad\times\mathbb{1}_{\{e_{li}=m\}}\frac{\log(P(t_{li},q_{li}=k|z_l=j,\gamma))}{B_{li}}\Bigg)\\
&=\sum_{l=1}^{T}\sum_{j=1}^{A}\sum_{i=1}^{Q_l}\sum_{k=1}^{N_{li}}\sum_{m=1}^{E}\Bigg(P(q_{li}=k|\gamma^{\upsilon-1},t,z_l=j)P(z_l=j|t,\gamma^{\upsilon-1},\lambda^{\upsilon-1},\pi^{\upsilon-1})\\
&\quad\times\mathbb{1}_{\{e_{li}=m\}}\frac{\log(\gamma_{jg(k)}(m))-\sum_{g=1}^{R}\gamma_{jg}\cdot O_{lg}[i]}{B_{li}}\Bigg).
\end{aligned} \tag{35}$$

Note the probability $P(q_{li}=k|\gamma^{\upsilon-1},t,z_l=j)$ does not depend on admixture time and proportion. We can simplify the expected log-likelihood further and switch the individual assignment hidden variable $q_{li}$ to a more rapidly computable group assignment hidden variable $q'_{li}$:

$$\begin{aligned}
&\sum_{l=1}^{T}\sum_{j=1}^{A}\sum_{i=1}^{Q_l}\sum_{k=1}^{N_{li}}\sum_{m=1}^{E}\Bigg(P(q_{li}=k|\gamma^{\upsilon-1},z_l=j)P(z_l=j|t,\gamma^{\upsilon-1},\lambda^{\upsilon-1},\pi^{\upsilon-1})\\
&\quad\times\mathbb{1}_{\{e_{li}=m\}}\frac{\log(\gamma_{jg(k)}(m))}{B_{li}}\Bigg)\\
&-\sum_{l=1}^{T}\sum_{j=1}^{A}\sum_{i=1}^{Q_l}\sum_{k=1}^{N_{li}}\sum_{m=1}^{E}\Bigg(P(q_{li}=k|\gamma^{\upsilon-1},z_l=j)P(z_l=j|t,\gamma^{\upsilon-1},\lambda^{\upsilon-1},\pi^{\upsilon-1})\\
&\quad\times\mathbb{1}_{\{e_{li}=m\}}\frac{\sum_{g=1}^{R}\gamma_{jg}\cdot O_{lg}[i]}{B_{li}}\Bigg).
\end{aligned} \tag{36}$$

We group the individuals based on group assignment in the first term (red term) and use $\sum_{k=1}^{N_{li}}P(q_{li}=k|\gamma^{\upsilon-1},z_l=j)=1$ in the second term (blue term).

$$\begin{aligned}
&\sum_{l=1}^{T}\sum_{j=1}^{A}\sum_{i=1}^{Q_l}\sum_{g=1}^{R}\sum_{m=1}^{E}\Bigg(P(q'_{li}=g|\gamma^{\upsilon-1},z_l=j)P(z_l=j|t,\gamma^{\upsilon-1},\lambda^{\upsilon-1},\pi^{\upsilon-1})\\
&\quad\times\mathbb{1}_{\{e_{li}=m\}}\frac{\log(\gamma_{jg}(m))}{B_{li}}\Bigg)\\
&-\sum_{l=1}^{T}\sum_{j=1}^{A}\sum_{i=1}^{Q_l}\sum_{m=1}^{E}\Bigg(P(z_l=j|t,\gamma^{\upsilon-1},\lambda^{\upsilon-1},\pi^{\upsilon-1})\mathbb{1}_{\{e_{li}=m\}}\sum_{g=1}^{R}\frac{\gamma_{jg}\cdot O_{lg}[i]}{B_{li}}\Bigg).
\end{aligned} \tag{37}$$

It is important to note that we use the group-wise assignment $q'_{li}$ moving forward instead of the individual-wise assignment $q_{li}$. Let us denote $U^{v-1}_{l,i,g,j,m} = P(q'_{li} = g|\gamma^{v-1}, z_l = j)$ which is the first probability in the red term above, corresponding to the membership "group g individual coalesces in the $i$-th coalescence event". And, let us denote $T^{v-1}_{l,j} = P(z_l = j|t, \gamma^{v-1}, \lambda^{v-1}, \pi^{v-1})$ which is the first probability in the blue term above, corresponding to the local ancestry of the target.

We use the forward-backward algorithm for HMMs to evaluate $T_{l,j}$ in Section 2.5.1; if alternatively the HMM is turned off, we assume that local ancestry is independent across loci. We calculate $U^{v-1}_{l,i,g,j,m}$ based on our assumed model that the probability of coalescing with a reference individual is directly proportional to its coalescence rate and for a group of individuals on the weighted mean of their individual coalescence rates (see Section 2.1):

$$U^{v-1}_{l,i,g,j,m} = P(q'_{li} = g|\gamma^{v-1}, z_l = j) = \frac{\gamma^{v-1}_{jg}(m) n_{lig}}{\sum_{g=1}^{R} \gamma^{v-1}_{jg}(m) n_{lig}}. \tag{38}$$

##### 2.4.2 M-step

Once we have the expected log-likelihood, we can maximise it in the M-step. We substitute $U^{v-1}_{l,i,g,j,m}$ and $T^{v-1}_{l,j}$ as hidden variables from Equation 38 and the HMM forward-backward expectation into Equation 37. We then maximise the resulting expression with respect to $\gamma_{jg}(m)$:

$$\begin{aligned} &\sum_{l=1}^{T}\sum_{i=1}^{Q_l}\sum_{j=1}^{A}\sum_{g=1}^{R}\sum_{m=1}^{E}\left(U^{v-1}_{l,i,g,j,m} T^{v-1}_{l,j} \mathbb{1}_{\{e_{li}=m\}} \frac{\log(\gamma_{jg}(m))}{B_{li}}\right) \\ &-\sum_{l=1}^{T}\sum_{i=1}^{Q_l}\sum_{j=1}^{A}\sum_{m=1}^{E}\left(T^{v-1}_{l,j} \mathbb{1}_{\{e_{li}=m\}} \sum_{g=1}^{R} \frac{\gamma_{jg} \cdot O_{lg}[i]}{B_{li}}\right). \end{aligned} \tag{39}$$

Differentiating with respect to $\gamma_{jg}(m)$ and setting it to 0 we get:

$$\sum_{l=1}^{T}\sum_{i=1}^{Q_l}\left(\frac{\mathbb{1}_{\{e_{li}=m\}} U^{v-1}_{l,i,g,j,m} T^{v-1}_{l,j}}{\gamma_{jg}(m) B_{li}}\right) - \sum_{l=1}^{T}\sum_{i=1}^{Q_l}\left(\frac{\mathbb{1}_{\{e_{li}=m\}} T^{v-1}_{l,j} O_{lg}[i]}{B_{li}}\right) = 0 \tag{40}$$

$$\begin{aligned} \gamma^{v}_{jg}(m) &= \frac{\sum_{l=1}^{T}\sum_{i=1}^{Q_l} \mathbb{1}_{\{e_{li}=m\}} U^{v-1}_{l,i,g,j,m} T^{v-1}_{l,j} / B_{li}}{\sum_{l=1}^{T}\sum_{i=1}^{Q_l} \mathbb{1}_{\{e_{li}=m\}} T^{v-1}_{l,j} O_{lg}[i] / B_{li}} \\ \gamma^{v}_{jg}(m) &= \frac{\sum_{l=1}^{T}\sum_{i=1}^{Q_l} \mathbb{1}_{\{e_{li}=m\}} \frac{\gamma^{v-1}_{jg}(m) n_{lig}}{\sum_{g=1}^{R}\gamma^{v-1}_{jg}(m) n_{lig}} T^{v-1}_{l,j} / B_{li}}{\sum_{l=1}^{T}\sum_{i=1}^{Q_l} \mathbb{1}_{\{e_{li}=m\}} T^{v-1}_{l,j} O_{lg}[i] / B_{li}}. \end{aligned} \tag{41}$$

Notice that the coalescence rate estimates closely resemble the pairwise maximum likelihood estimate for two lineages derived in Equation 10 in Section 1.1.

#### 2.5 EM derivation: inference for admixture time and proportions

We assume a Markov model to express the likelihood for Term 1 in Equation 28. In addition to the local ancestry hidden variable ($z$), we introduce another hidden variable, $\omega_l$, which records the number of recombination events since admixture between positions $l-1$ and $l$. The distribution of $\omega_l$ depends on a user-provided recombination map, specifying the genetic distance between two grid points, denoted as $r_l$, and the time since admixture $\lambda$. The "complete-data likelihood" for this term can then be written as:

$$P(z,\omega \mid \lambda,\pi) = \prod_{l=1}^{T}\Big(\prod_{j=1}^{A}\pi_j^{I\{z_l=j,\omega_l>0\}}\Big)P(z_{l-1})^{I\{\omega_l=0\}}\frac{(\lambda r_l)^{\omega_l}e^{-\lambda r_l}}{\omega_l!}, \tag{42}$$

where the likelihood for the number of recombination events follows a Poisson counting process. Notably, a local ancestry change occurs only if there is at least one recombination event. As a boundary condition, we impose $P(\omega_l > 0) = 1$ for $l = 1$. The model parameters are $\lambda$, representing the admixture time, and $\pi$, the global cluster proportions. The corresponding log-likelihood can be expressed as:

$$\log P(z,\omega_l \mid \lambda,\pi) = \sum_{l=1}^{T}\Big(\sum_{j=1}^{A} I\{z_l = j,\omega_l > 0\}\log\pi_j\Big) + \omega_l\log(\lambda r_l) - \lambda r_l + \text{constants}. \tag{43}$$

Note in the above equation, we have removed terms (called them 'constants') which do not depend on $\lambda$ or $\pi$, as they will be discarded in the M-step.

##### 2.5.1 E-step

To perform the EM algorithm, we compute the expected log-likelihood of the complete data, where the expectation is taken over the hidden variables conditioned on the observed data and the model parameters from the previous iteration (including $\gamma$).

$$\mathbb{E}_{z,\omega_l|\gamma^{\upsilon-1},\lambda^{\upsilon-1},\pi^{\upsilon-1},t}\Big(\sum_{l=1}^{T}\Big(\sum_{j=1}^{A} I\{z_l = j,\omega_l > 0\}\log\pi_j\Big) + \omega_l\log(\lambda r_l) - \lambda r_l\Big). \tag{44}$$

We start simplifying the expected log-likelihood by taking the expectation inside the summation, and transforming the expected value of indicator random variables as probabilities.

$$\sum_{l=1}^{T}\Big(\sum_{j=1}^{A} P(z_l = j,\omega_l > 0|\gamma^{\upsilon-1},\lambda^{\upsilon-1},\pi^{\upsilon-1},t)\log\pi_j\Big) + \mathbb{E}(\omega_l)\log(\lambda r_l) - \lambda r_l \tag{45}$$

where the expectation is taken over the hidden variables conditioned on the data and previous iterations' parameter updates. We are interested in inferring three terms, (1) $P(z_l = j,\omega_l > 0|\gamma^{\upsilon-1},\lambda^{\upsilon-1},\pi^{\upsilon-1},t)$, (2) $\mathbb{E}(\omega_l|\gamma^{\upsilon-1},\lambda^{\upsilon-1},\pi^{\upsilon-1},t)$ and (3) $P(z_l = j|t,\gamma^{\upsilon-1},\lambda^{\upsilon-1},\pi^{\upsilon-1})$. We start with the first term:

$$\begin{aligned}
P(z_l = j,\omega_l > 0|P^{\upsilon-1},t) &= P(z_l = j|P^{\upsilon-1},t)P(\omega_l > 0|z_l = j,P^{\upsilon-1},t)\\
&= P(z_l = j|P^{\upsilon-1},t)\sum_{i=1}^{A}P(\omega_l > 0|z_l = j,z_{l-1} = i,P^{\upsilon-1},t)P(z_{l-1} = i|z_l = j,P^{\upsilon-1},t)\\
&= \sum_{i=1}^{A}P(z_l = j|P^{\upsilon-1},t)P(\omega_l > 0|z_l = j,z_{l-1} = i,P^{\upsilon-1},t)P(z_{l-1} = i|z_l = j,P^{\upsilon-1},t)\\
&= \sum_{i=1}^{A}P(\omega_l > 0|z_l = j,z_{l-1} = i,P^{\upsilon-1})P(z_l = j,z_{l-1} = i|P^{\upsilon-1},t)
\end{aligned} \tag{46}$$

where $P^{\upsilon-1} = (\gamma^{\upsilon-1},\lambda^{\upsilon-1},\pi^{\upsilon-1})$ is a short-hand for the tuple of all previous parameters. We denote the second term (coloured in blue) as $\epsilon_{ij}(l)$ as it denotes the probability of being in state $i$ and state $j$ at positions $l-1$ and $l$ respectively. Similar to the Baum-Welch algorithm, we can solve for $\epsilon_{ij}(l)$ through the forward-backward pass in HMMs.

$$\begin{aligned}
\epsilon_{ij}(l) &= P(z_l = j, z_{l-1} = i | P^{v-1}, t) \\
&= \frac{P(z_l = j, z_{l-1} = i, t | P^{v-1})}{P(t|P^{v-1})} \\
&= \frac{P(t_1.., t_{l-1}, z_{l-1} = i|P^{v-1})P(z_l = j|z_{l-1} = i, P^{v-1})P(t_l.., t_T|z_l = j, P^{v-1})P(t_l|z_l, P^{v-1})}{P(t|P^{v-1})} \\
&= \frac{P(t_1.., t_{l-1}, z_{l-1} = i|P^{v-1})P(z_l = j|z_{l-1} = i, P^{v-1})P(t_l.., t_T|z_l = j, P^{v-1})P(t_l|z_l, P^{v-1})}{\sum_{j=1}^{A}\sum_{i=1}^{A} P(t_1.., t_{l-1}, z_{l-1} = i|P^{v-1})P(z_l = j|z_{l-1} = i, P^{v-1})P(t_l.., t_T|z_l = j, P^{v-1})P(t_l|z_l, P^{v-1})}
\end{aligned} \tag{47}$$

where, $P(t_1.., t_{l-1}, z_{l-1} = i|P^{v-1})$ is called the forward-pass and $P(t_l.., t_T|z_l = j, P^{v-1})$ is called the backward-pass. The transition probability for the HMM is given as follows:

$$P(z_l = j|z_{l-1} = i, P^{v-1}) = \begin{cases} (1 - e^{-r_l\lambda^{v-1}})\pi_j^{v-1}, & \text{if } i \neq j \\ e^{-r_l\lambda^{v-1}} + (1 - e^{-r_l\lambda^{v-1}})\pi_j^{v-1}, & \text{if } i = j \end{cases} \tag{48}$$

whereas the emission probability for the HMM $P(t_l|z_l, P^{v-1})$ only depends on $\gamma^{v-1}$ and follows from the scaled likelihood in Equation 29:

$$P(t_l|z_l = j, P^{v-1}) = \Big(\frac{\sum_{g=1}^{R}\gamma_{jg}^{v-1}(m)n_{lig}}{\sum_{g=1}^{R} n_{lig}} e^{-\sum_{g=1}^{R}\gamma_{jg}^{v-1}\cdot O_{lg}[i]}\Big)^{\frac{1}{B_{li}}}. \tag{49}$$

The first term (highlighted in red) in Equation 46 represents the probability of at least one recombination event, conditioned on the local ancestry states $z_{l-1}$ and $z_l$. It is expressed as follows:

$$P(\omega_l > 0|z_l = j, z_{l-1} = i, P^{v-1}) = \begin{cases} 1, & \text{if } i \neq j \\ \frac{(1-e^{-r_l\lambda^{v-1}})\pi_j^{v-1}}{e^{-r_l\lambda^{v-1}} + (1-e^{-r_l\lambda^{v-1}})\pi_j^{v-1}}, & \text{if } i = j. \end{cases} \tag{50}$$

The next term to be computed in the E-step is $E(\omega_l)$, where the expectation is taken over the hidden variables $z_l$ and $\omega_l$, conditioned on the observed data and the model parameters from the previous iteration. This expectation can be simplified as follows:

$$\begin{aligned}
\mathbb{E}(\omega_l|P^{v-1}, t) &= \mathbb{E}(\omega_l|\omega_l > 0, P^{v-1}, t)P(\omega_l > 0|P^{v-1}, t) \\
&= \mathbb{E}(\omega_l|\omega_l > 0, P^{v-1})P(\omega_l > 0|P^{v-1}, t) \\
&= \frac{\lambda^{v-1}r_l}{1 - e^{-\lambda^{v-1}r_l}} P(\omega_l > 0|P^{v-1}, t).
\end{aligned} \tag{51}$$

In order to evaluate $P(\omega_l > 0|P^{v-1}, t)$ we do the following:

$$\begin{aligned}
P(\omega_l > 0|P^{v-1}, t) &= \sum_{j=1}^{A}\sum_{j=1}^{A} P(\omega_l > 0|z_l = j, z_{l-1} = i, P^{v-1}, t)P(z_l = j, z_{l-1} = i|P^{v-1}, t) \\
&= \sum_{j=1}^{A}\sum_{j=1}^{A} P(\omega_l > 0|z_l = j, z_{l-1} = i, P^{v-1})\epsilon_{ij}(l).
\end{aligned} \tag{52}$$

The terms required in Equation 52 have already been calculated in Equations 47 and 50. As a boundary condition, we impose $P(\omega_l > 0) = 1$ for $l = 1$. Finally, to compute the local ancestry estimate $T_{l,j}^{v-1} = P(z_l = j \mid t, \gamma^{v-1}, \lambda^{v-1}, \pi^{v-1})$ needed in Equation 41, we perform a forward-backward pass:

$$T_{l,j}^{v-1} = P(z_l = j|t, \gamma^{v-1}, \lambda^{v-1}, \pi^{v-1}) = \frac{P(t_1.., t_{l-1}, z_{l-1} = i|P^{v-1})P(t_l.., t_T|z_l = j, P^{v-1})}{\sum_{j=1}^{A} P(t_1.., t_{l-1}, z_{l-1} = i|P^{v-1})P(t_l.., t_T|z_l = j, P^{v-1})} \tag{53}$$

where, $P(t_1.., t_{l-1}, z_{l-1} = i|P^{v-1})$ and $P(t_l.., t_T|z_l = j, P^{v-1})$ are the forward and backward probabilities, respectively.

##### 2.5.2 M-step

Once all the necessary equations in the E-step have been determined, we proceed to the M-step, where we maximise the expected log-likelihood. Specifically, we optimise with respect to $\lambda$ and the cluster proportions $\pi$ by maximising the following equation:

$$\sum_{l=1}^{T}\left(\sum_{j=1}^{A} P(z_l = j, \omega_l > 0|\gamma^{\upsilon-1}, \lambda^{\upsilon-1}, \pi^{\upsilon-1}, t)\log\pi_j\right) + \mathbb{E}(\omega_l)\log(\lambda r_l) - \lambda r_l. \tag{54}$$

1. Differentiating with respect to $\pi_j$ and setting to zero, we get:

$$\pi_j^{\upsilon} \propto \sum_{l=1}^{T} P(z_l = j, \omega_l > 0|\gamma^{\upsilon-1}, \lambda^{\upsilon-1}, \pi^{\upsilon-1}, t)$$

$$\pi_j^{\upsilon} = \frac{\sum_{l=1}^{T} P(z_l = j, \omega_l > 0|\gamma^{\upsilon-1}, \lambda^{\upsilon-1}, \pi^{\upsilon-1}, t)}{\sum_{j=1}^{A}\sum_{l=1}^{T} P(z_l = j, \omega_l > 0|\gamma^{\upsilon-1}, \lambda^{\upsilon-1}, \pi^{\upsilon-1}, t)}. \tag{55}$$

   We get the term $P(z_l = j, \omega_l > 0|\gamma^{\upsilon-1}, \lambda^{\upsilon-1}, \pi^{\upsilon-1}, t)$ from the E-step Equation 46.

2. Differentiating with respect to $\lambda$ and setting to zero, we get:

$$\lambda^{\upsilon} = \frac{\sum_{l=1}^{T} \mathbb{E}(\omega_l|\gamma^{\upsilon-1}, \lambda^{\upsilon-1}, \pi^{\upsilon-1}, t)}{\sum_{l=1}^{T} r_l}. \tag{56}$$

   Note, we get the expected number of recombinations from Equation 51 in the E-step.

#### 2.6 Summary of the algorithm

We present a summary of the EM algorithm for GhostBuster here:

**Algorithm 1** EM Algorithm for GhostBuster

**Input:** Genealogies $\mathbf{X} = \{(t_1, n_1), (t_2, n_2), \ldots, (t_C, n_C)\}$ containing coalescence times and population counts for each coalescence event, number of components $A$, number of time epochs $E$, number of grid points $T$, total number of coalescence events $C$ and number of reference groups $R$.
**Preprocessing:** Calculate likelihood scaling term $B_c$ for each coalescence event $c = 1, 2, \ldots, C$, and $r_l$, genetic distance between two grid points
**Initialize:** Coalescence rates $\gamma_j^{(0)}$, mixing coefficients $\pi_j^{(0)}$ for each component $j = 1, 2, \ldots, A$ and admixture time $\lambda^{(0)}$
**Set:** Iteration $\upsilon = 1$
**repeat**
    **E-step:**
    **for** each component $j = 1, 2, \ldots, A$ **do**
        **for** each data point $(t_{li}, n_{li})$ **do**
▷ $t_{li}$ represent the coalescence time in $l$-th grid and $i$-th coalescence event
▷ $P^{\upsilon-1} = \{\gamma^{\upsilon-1}, \lambda^{\upsilon-1}, \pi^{\upsilon-1}\}$
            Compute the likelihood under the component:

$$P(t_l | z_l = j, P^{\upsilon-1}) = \Big( \frac{\sum_{g=1}^{R} \gamma_{jg}^{\upsilon-1}(m) n_{lig}}{\sum_{g=1}^{R} n_{lig}} e^{-\sum_{g=1}^{R} \gamma_{jg}^{\upsilon-1} \cdot O_{lg}[i]} \Big)^{\frac{1}{B_{li}}}$$

        **end for**
        Perform forward-backward pass with the HMM to get $P(z_l = j | X, P^{\upsilon-1})$
            see Equation 53
    **end for**
    **M-step:**
    **for** each component $j = 1, 2, \ldots, A$ **do**
        Update the coalescence rates:
            see Equation 41
        Update the cluster proportions:
            see Equation 55
        Update the admixture time:
            see Equation 56
    **end for**
    **Increment:** $\upsilon = \upsilon + 1$
**until** convergence
**Output:** Final parameters, $\{\gamma^{\upsilon}, \lambda^{\upsilon}, \pi^{\upsilon}\}$ and local ancestry posteriors, $P(z_l = j | X, P^{\upsilon})$ for $l = 1, 2, \ldots, T$ and $j = 1, 2, \ldots, A$

### 3 Algorithmic details for GhostBuster

#### 3.1 PCA visualisation of ancestral components

To visualise the clustering performed by GhostBuster, we perform Principal Component Analysis (PCA) on the coalescence count and opportunity matrices obtained from the genealogies. The coalescence count matrix is an $N_{\text{ref}} \times N_{\text{sites}}$ matrix, where the $(g,l)^{\text{th}}$ entry represents the total contribution of reference group $g$ to the coalescence events at genomic position $l$. We aggregate the contributions from all coalescence events involving the target individual at genomic position $l$, considering only those events that occur within the predefined start and end time epochs. These time epochs are aligned with the start and end epochs used in the GhostBuster analysis. These epochs control the time periods in which we expect to observe differences in coalescence patterns. We construct a similar matrix for the opportunity, also an $N_{\text{ref}} \times N_{\text{sites}}$ matrix, where each entry $g, l$ records the total opportunity for reference group $g$ at position $l$. The same time epochs are used as in the coalescence count matrix to ensure consistency.

The coalescence count and opportunity matrices are then stacked to form a $2N_{\text{ref}} \times N_{\text{sites}}$ matrix, which is mean-centred and standardised. We perform eigen-decomposition on this matrix to obtain the leading principal components. These principal components provide a visual interpretation of the clustering performed by GhostBuster, helping to better understand the patterns in coalescence across the genome.

#### 3.2 Coancestry curves to date admixture events

Given the local ancestry inferred using GhostBuster, we use coancestry curves to estimate the date of the admixture event. The joint probability of observing ancestry at a genetic distance $g$ under a single-pulse admixture model is given by:

$$\begin{aligned}
p_{AA}(g) &= \alpha^2 + \alpha(1-\alpha)\exp(-g\lambda) \\
p_{AB}(g) &= \alpha(1-\alpha) - \alpha(1-\alpha)\exp(-g\lambda) \\
p_{BB}(g) &= (1-\alpha)^2 + \alpha(1-\alpha)\exp(-g\lambda)
\end{aligned} \tag{57}$$

where $\alpha$ is the genome-wide proportion of ancestry 'A', $1-\alpha$ of ancestry 'B' and $\lambda$ is the admixture time in generations. We also consider admixture models with two-dates corresponding to two-pulses of gene flow. The joint probability of observing ancestry at a genetic distance $g$ under a double admixture model is given by:

$$\begin{aligned}
p_{AA}(g) &= \alpha_1^2(1-\alpha_3)^2 + \alpha_1^2\alpha_3(1-\alpha_3)\exp(-g\lambda_2) + \alpha_1\alpha_2(1-\alpha_3)\exp(-g\lambda_1) \\
p_{BB}(g) &= \alpha_2^2(1-\alpha_3)^2 + \alpha_2^2\alpha_3(1-\alpha_3)\exp(-g\lambda_2) + \alpha_1\alpha_2(1-\alpha_3)\exp(-g\lambda_1) \\
p_{AB}(g) = p_{BA}(g) &= \alpha_1\alpha_2(1-\alpha_3)^2 + \alpha_1\alpha_2\alpha_3(1-\alpha_3)\exp(-g\lambda_2) - \alpha_1\alpha_2(1-\alpha_3)\exp(-g\lambda_1) \\
p_{CA}(g) = p_{AC}(g) &= \alpha_3(1-\alpha_3)\alpha_1 - \alpha_3(1-\alpha_3)\alpha_1\exp(-g\lambda_2) \\
p_{CB}(g) = p_{BC}(g) &= \alpha_3(1-\alpha_3)\alpha_2 - \alpha_3(1-\alpha_3)\alpha_2\exp(-g\lambda_2) \\
p_{CC}(g) &= \alpha_3^2 + \alpha_3(1-\alpha_3)\exp(-g\lambda_2)
\end{aligned} \tag{58}$$

where, $\alpha_1(1-\alpha_3)$ is the genome-wide proportion of ancestry 'A', $\alpha_2(1-\alpha_3)$ of ancestry 'B', $\alpha_3$ of ancestry 'C' and $\lambda_1$, $\lambda_2$ are the corresponding admixture times. Finally, we consider admixture models corresponding to a constant migration rate and continuous admixture. In this case, the joint probability of observing ancestry at a genetic distance $g$ is given by:

$$p_{AA}(g) = \alpha^2 + \alpha(1-\alpha)\sum_{j=\lambda_e}^{\lambda_s} w_j \exp(-gj)$$
$$p_{AB}(g) = \alpha(1-\alpha) - \alpha(1-\alpha)\sum_{j=\lambda_e}^{\lambda_s} w_j \exp(-gj)$$
$$p_{BB}(g) = (1-\alpha)^2 + \alpha(1-\alpha)\sum_{j=\lambda_e}^{\lambda_s} w_j \exp(-gj) \tag{59}$$

where, $w_j = \frac{\mu(1-\mu)^{\lambda_s - j}}{1-(1-\mu)^{\lambda_s-\lambda_e+1}}$; $\lambda_s$ and $\lambda_e$ are the start and end time of the admixture, $\mu$ is the constant migration rate. Detailed derivation of the analytical coancestry curves can be found in the supplementary information of [14].

We fit the exponential distribution to all possible ancestry combinations and jointly maximise the log-likelihood (assuming a Gaussian error model on the residuals from least-squares fits) using the Nelder–Mead algorithm [14, 15]. In the case of three-way or more complex admixture, we fit coancestry curves for each pair of ancestries separately, estimating the dates for each event independently. We use 20 block-jackknife across chromosomes to estimate standard errors when dating the admixture event. Additionally, we use diploid ancestry calls (summing the posteriors of two haplotypes) to prevent phasing errors from biasing the admixture dating.

Genetic drift can cause the fixation or loss of certain alleles, introducing a downward bias in the dating estimates derived from coancestry curves. This effect is particularly pronounced in small populations or when considering ancient events, where the influence of drift is much greater. To mitigate this bias, we normalise the coancestry curves by the coancestry curves generated from randomly sampling haplotype pairs from different individuals, also known as pseudo-individuals. As previously done in Globetrotter [14], this approach corrects for the impact of genetic drift and provides an unbiased admixture date.

Another source of error in admixture dating can arise from inaccuracies in recombination maps. This is particularly relevant for dating ancient events, where errors on the scale of tens of kilo-bases in the recombination map can downward-bias the inferred date [16]. In our case, this issue does not affect the simulated examples, as we assume a perfect recombination map, but might have some impact on real data, especially the old admixture dates. However, due to the lack of suitable error estimates for the recombination map of the population being analysed, we highlight this as a cautionary note for the interpretation of results from real data.

Finally, the effects of selection and the evolution of recombination maps can influence admixture dates; however, we assume these factors have a minimal impact on our real-data examples.

#### 3.3 Calculation of node persistence

The majority of nodes in adjacent trees are shared. Accurate calculation of node persistence is critical in our likelihood estimation, as it prevents the over-counting of identical or similar coalescence events. In true genealogies, nodes can be distinguished by their coalescence times. These coalescence times remain consistent across adjacent trees unless a recombination event affects the corresponding node.

However, in inferred genealogies, the coalescence times are not precisely identical, even in the absence of recombination, as each tree is dated independently based on its structure. To identify equivalent nodes in inferred genealogies, we examine the descendant composition of these nodes. Specifically, we store a vector of length equal to the number of reference samples, representing the descendant composition of the lineage coalescing with the target lineage. In this vector, the $m$th entry is set to 1 if haplotype $m$ descends from the coalescence event, and 0 otherwise. Similar to defining equivalent branches in Relate [10], we define two nodes as equivalent if the correlation coefficient between their descendant composition vectors exceeds a predefined threshold (default = 0.5). To ensure that each node is matched with at most one other node, we rank pairs of candidate nodes in descending order of their correlation coefficient. Nodes are then associated based on their position in this sorted list, with any pair involving a node already associated with another node being removed.

In true genealogies, where we have access to actual node persistence, we observed that the approximate node persistence reasonably aligns with the true values (see Supplementary Note Figure **4** for comparison).

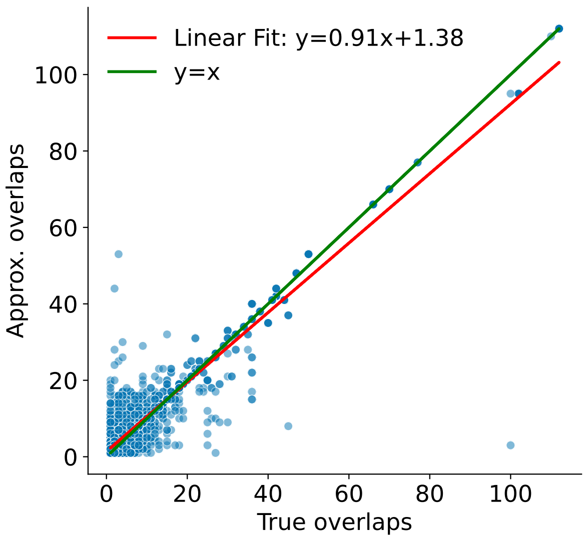

**Supplementary Note Figure 4**: **Comparison of Approximate and True Node Persistence in Simulated Trees.** True node persistence is determined by identifying the exact coalescence times across neighbouring trees, while approximate node persistence is estimated based on the correlation of descendant sets between adjacent trees. In this case, the correlation cutoff for approximate node persistence is set to 0.5, with the number of overlaps indicating the number of 10kb intervals for which the node persists.

#### 3.4 Handling uncertain coalescence events

Since the mutation and recombination rates in humans are approximately similar, multiple tree structures often exist to explain the genetic variation at a given site, making genealogy inference uncertain and potentially inaccurate. To address this uncertainty, we scale the likelihood of our mixture model based on the confidence in coalescence events, which is evaluated by the presence of mutations above the node.

Specifically, we trace the coalescence events along the target lineage backwards in time, identifying subsets of nodes (or coalescence events) that are jointly supported by at least one mutation (see Supplementary Note Figure **5**). For each such subset of events, we are not certain in which order lineages will coalesce. For instance, while the target lineage may be coalescing with each reference lineage in turn as observed in the genealogy, in truth, these reference lineages may coalesce with each other first before coalescing with the target lineage. To incorporate this uncertainty, we calculate the expected number of coalescence events (or nodes) involving the target lineage. We do this by effectively allowing coalescences between lineages to be shuffled.

We assume that clades coalescing with the target are resolved: that is, they have mutations supporting their tree structure. We further assume that unresolved coalescence events occur around the same time, simplifying their contribution to the likelihood. Given $i$ lineages, the probability that the next coalescence involves the target lineage is $\frac{i}{\binom{i+1}{2}} = \frac{2}{i+1}$. Therefore, the expected number of coalescence events, conditioned on the presence of $C$ unresolved events, can be computed as the sum of the probability of coalescing with the target for the $C$ independent coalescence events:

$$\mathcal{E}[C] = \sum_{i=1}^{C} \frac{i}{\binom{i+1}{2}} \tag{60}$$

$$= \sum_{i=1}^{C} \frac{2}{i+1}. \tag{61}$$

We scale the likelihood for each node by the expected number of coalescence events divided by the observed number of coalescence events, that is $\frac{\mathcal{E}[C]}{C}$. For cases where there are 1, 2, or 3 unresolved nodes, the corresponding expected number of coalescence events equals 1, $\frac{5}{3}$, and $\frac{13}{6}$, leading to effectively scaling the likelihood by 1, $\frac{5}{6}$, and $\frac{13}{18}$, respectively.

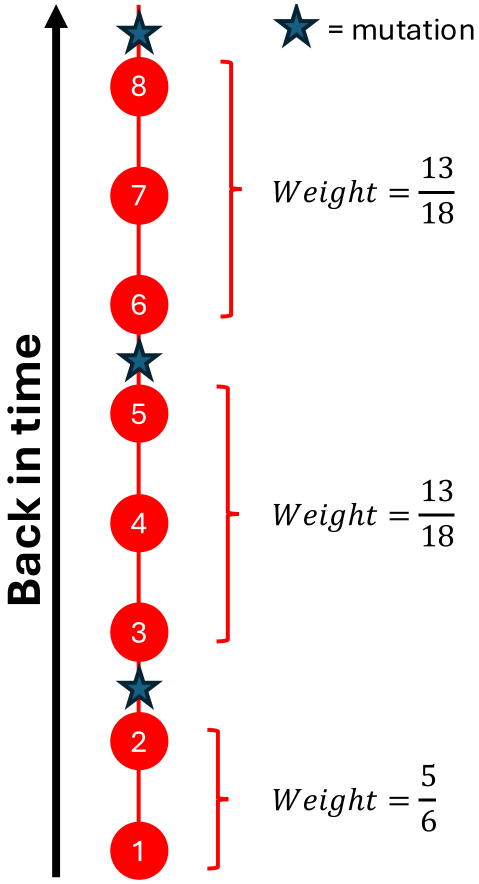

**Supplementary Note Figure 5**: **Handling uncertainty in coalescence events.** We calculate the expected number of coalescence events for subsets of nodes jointly supported by a mutation. In this example, there are three such subsets on the target lineage, with weights for each coalescence event calculated based on the number of nodes in the subset.

### 4 Additional Simulations

#### 4.1 Four-way admixture

In this scenario, we assume four reference populations (A, B, C, and D), each contributing 25% to form the focal individual 5,000 years ago. Populations A and B split 50,000 years ago, as do populations C and D, with all populations merging back together 100,000 years ago (see Supplementary Note Figure **6**). We assume a constant diploid population size of 20,000 for all populations after their split and 10,000 for the super-populations before 50,000 years ago. To assess the case without a ghost population, we sample 10 diploid individuals from each population and five focal (admixed) individuals. For the ghost population scenario, we sample 10 individuals each from populations A, B, and C, with none from population D. We used a constant mutation rate of $1.25 \times 10^{-8}$ per base per generation and the HapMap3 human recombination rate map. We use `msprime` [17] for this simulation.

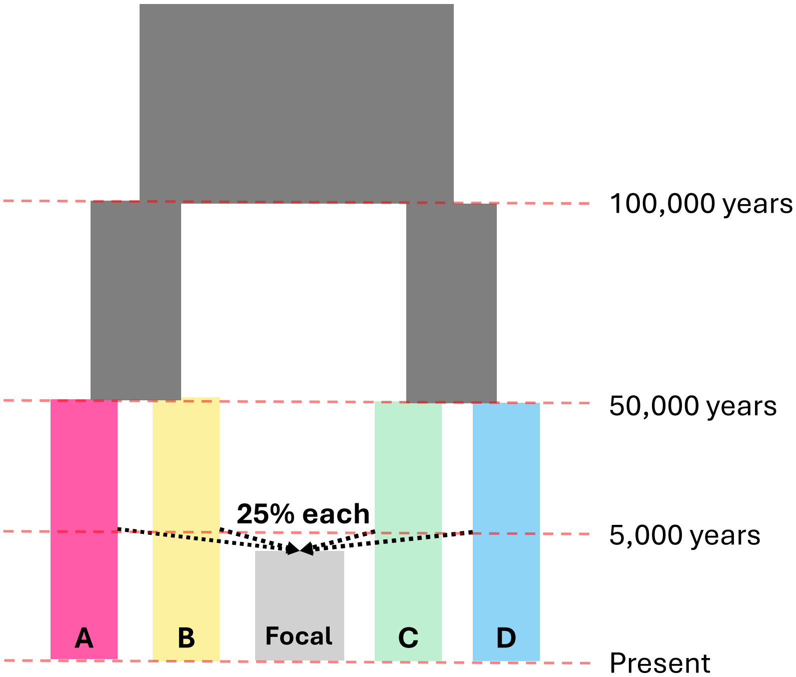

**Supplementary Note Figure 6**: **Simulation design for four-way admixture scenario.**

We applied GhostBuster to trees inferred using Relate, assuming a constant mutation rate of $1.25 \times 10^{-8}$ per base per generation, while results comparing with true trees simulated using `msprime` are also presented. Additionally, we only focused on coalescence events occurring from 0 to 50,000 years ago.

We first tested the method with access to all four reference populations and applied our approach to decompose the focal individual's ancestry into four distinct components or clusters. By design, the focal individual is symmetrically related to all four source groups, and so simple genome-wide average coalescence rates show no indication of admixture. We apply GhostBuster, starting with a random initialisation of coalescence rates and proportions and fit our model over 200 EM iterations until the log-likelihood converges. The resulting inverse coalescence rates represent the four clusters that best differentiated the coalescence events for the focal individual (see Supplementary Figure **2**a). The inverse coalescence rate (ICR) for each component reflects its affinity toward one of the four reference populations—a lower ICR value indicates faster coalescence with that specific population. Our analysis revealed that the four clusters identified by GhostBuster contributed approximately 25% to the focal individual's ancestry, with each cluster more closely related to one of the four reference populations. For example, component 1 was most similar to population D, component 2 to population A, component 3 to population C, and component 4 to population B. The clusters were clearly distinguishable using the top four principal components (PCs) of the coalescence count and opportunity matrices (see Supplementary Figure **2**b). Further evaluation based on the expected coefficient of determination and held-out log-likelihood indicated that four components provided the best fit to the data, with diminishing returns in held-out log-likelihood beyond four components (see Supplementary Figure **2**c). We also accurately dated the admixture event to 185.1 generations, or 5,182.8 years, using coancestry curves (see Supplementary Figure **2**d). Finally, we assessed the accuracy and calibration of our local ancestry inference by comparing it with the simulated ground truth, demonstrating strong correlation and low calibration error (see Supplementary Figure **2**e-g).

Next, we removed population D (without loss of generality) from the genealogies and attempted to decompose the focal individual. This scenario simulates the presence of a ghost population, as population D contributed to the focal individual's genetic makeup but is not included in the reference panel. We reran our method, fixing the number of components to four. The four components still contributed roughly 25% to the total ancestry. The inverse coalescence rates (ICRs) for each component only existed for populations A, B, and C, as no individuals from population D were sampled. Component 2's ICR was closest to population B, component 3 was closest to population A, and component 4 was closest to population C (see Supplementary Figure **3**a). However, component 1 was more distantly related to all three populations, indicating slower coalescence with each. This could be the ancestry relating to population D, which may have been tagged by its slower coalescence rates with populations A-C. Similar to the run with access to population D, we performed PCA visualisation (see Supplementary Figure **3**b), evaluation of the number of clusters (see Supplementary Figure **3**c), coancestry curve dating (see Supplementary Figure **3**d), and accuracy and calibration of local ancestry inference (see Supplementary Figure **3**e-g). When comparing local ancestry with the simulated ground truth, we found a strong correlation, with an $R^2$ value of 0.86, indicating that component 1 effectively captured the ancestry related to population D without requiring direct samples from that population.

#### 4.2 Denisovan-like Introgression into Papuans

This scenario uses the simulation model and script provided by [18] to simulate Denisovan introgression into Papuans. We assume three reference populations (resembling Papuans, Africans, and Denisovans), where Papuans and Africans split 72,036 years ago, and Denisovans split from modern humans 656,908 years ago (see Supplementary Note Figure **7**). Denisovans admix with Papuans around 43,935 years ago with a 5% admixture proportion. The population sizes are $3,899$ for modern Papuans, 27,122 for Africans, and 4,947 for Denisovans before they went extinct. We also simulate the Out-of-Africa bottleneck, where the Papuan population size drops to 136. We fit five Papuan individuals, using 25 Papuans and 25 Africans as the reference panel. Additionally, we assess the power and compare local ancestry estimates with and without an ancient Denisovan sample dated to 67,570 years ago. We used a constant mutation rate of $1.25 \times 10^{-8}$ per base per generation and the HapMap3 human recombination rate map. We again use `msprime` [17].

Similar to our previous simulation, we applied GhostBuster to trees inferred by Relate, assuming a constant mutation rate of $1.25 \times 10^{-8}$ per base per generation. We focused only on coalescence events occurring from 10,000 to 1,000,000 years for this analysis.

We begin our analysis by assuming we have access to one ancient Denisovan sample. We decompose five out of the 25 Papuan individuals, assuming two distinct components. As with the previous simulation, we randomly initialise the parameters and fit the EM algorithm for 200 iterations until the log-likelihood converges. The converged components reveal a major component, contributing around 95.85% to the local ancestry, and a minor component contributing 4.15% (see Supplementary Figure **4**a). The inverse coalescence rates (ICRs) characterising these components differ significantly. Component 1 demonstrates a slower coalescence rate with the ancient Denisovan sample and a relatively faster coalescence with the modern Mbuti samples. In contrast, component 2 exhibits a very fast coalescence rate with Denisovans but a slower coalescence with modern Mbuti and Papuan samples. It is important to note that the most striking difference between the two components is the vastly accelerated coalescence with Denisovans in the minority group, up to 1,000×

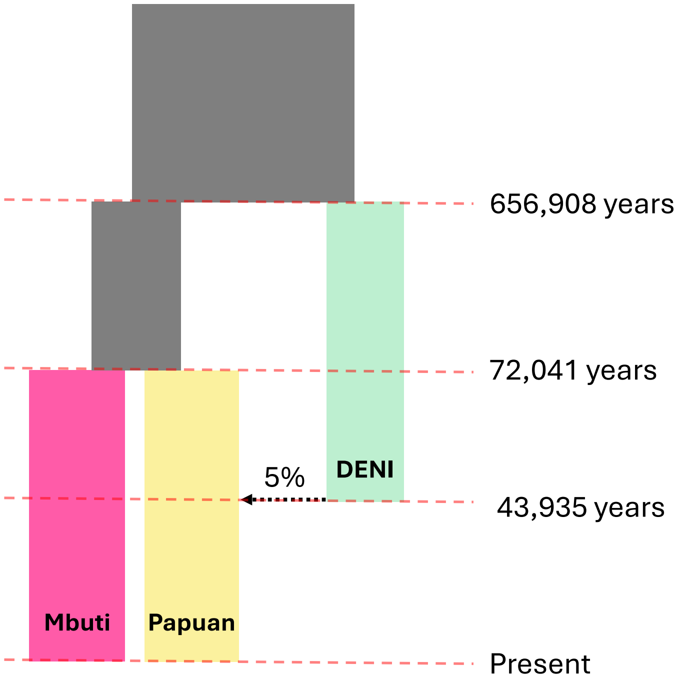

**Supplementary Note Figure 7**: **Simulation design for Denisovan-like introgression into Papuan scenario.**

faster compared to the majority group. However, there is also a substantial difference in coalescence with modern groups, which we will leverage to decompose the ancestry without access to the ancient Denisovan sample.

The clusters were distinguishable using the top four principal components (PCs) of the coalescence count and opportunity matrices (see Supplementary Figure **4**b). Further evaluation based on the expected coefficient of determination and held-out log-likelihood indicated that two components provided the best fit to the data, with diminishing returns in held-out log-likelihood beyond four components (see Supplementary Figure **4**c). We also accurately dated the admixture event to 1,546.7 generations, or 44,854.3 years, using coancestry curves (see Supplementary Figure **4**d). Finally, we assessed the accuracy and calibration of our local ancestry inference by comparing it with the simulated ground truth and HMMIX, a recent HMM-based method tailored to find archaic introgression such as this one [18], demonstrating strong correlation with the ground truth, surpassing HMMIX, and showing low calibration error (see Supplementary Figure **4**e-g).

Next, we removed the Denisovan samples from our analysis and decomposed the same Papuan individuals into two components. In this scenario, the Denisovan ancestry in Papuans can be thought of as a ghost population. Our method converged on two components, contributing approximately 95.9% and 4.1% to the local ancestry, respectively (see Supplementary Figure **5**a). The ICRs of the minor component closely matched those of the minor component from the earlier analysis that included the Denisovan sample, showing slower coalescence with Mbuti and Papuans (see Supplementary Figure **4**a). Similar to the run with access to the Denisovan genome, we performed PCA visualisation (see Supplementary Figure **5**b), evaluation of the number of clusters (see Supplementary Figure **5**c), coancestry curve dating (see Supplementary Figure **5**d), and accuracy and calibration of local ancestry inference (see Supplementary Figure **5**e-g). To verify that the minor component indeed represents Denisovan ancestry, we compared the local ancestry inferred using GhostBuster with the simulated ground truth and found a high degree of correlation ($R^2 = 0.62$), surpassing HMMIX in this case [18].

In order to test the robustness and false positive rate of our method, we attempted to decompose five Mbuti individuals in the same simulation. The Mbuti individuals are panmictic, with no simulated population structure or admixture. When we forced GhostBuster to find two components for Relate trees, it identified two components with varying coalescence rates with Papuans (see Supplementary Figure **6**a). At face value, this pattern could be interpreted as evidence for subtle heterogeneity in how different regions of the Mbuti genome relate to Papuans. However, these components were not clearly separated in the PC plots (see Supplementary Figure **6**b). In terms of held out log likelihood, the two-component solution offered only a slight improvement compared to a single component model (see Supplementary Figure **6**c). Moreover, the histogram of the local ancestry posteriors and the expected R-squared of 0.49 both suggest that the inferred local ancestry is highly uncertain (see Supplementary Figure **6**c,e). Dating this event using coancestry curves yielded an admixture date of around 2,800 generations, or 81,200 years, which is older than the simulated Mbuti and Papuan split time, raising further suspicion (see Supplementary Figure **6**d).

A key consideration is that the reference panel used in this analysis, Papuans, is itself admixed. Papuans harbour Denisovan ancestry, and this ancestry is not uniformly distributed across the genome. As a result, genomic regions in Papuans with different levels of Denisovan-related drift may exhibit different coalescence patterns with the Mbuti. GhostBuster may therefore be partitioning the Mbuti genome according to heterogeneity in the reference population rather than true structure within the Mbuti. Taken together, the weak separation in PC space, minimal improvement in held-out likelihood, low expected R-squared, and implausibly ancient dating all suggest that while a two-component model can technically be fit, it does not provide compelling evidence for admixture in Mbuti. More generally, decomposition results are relative to the chosen reference panel. It is therefore advisable to run GhostBuster across multiple targets while alternating references to distinguish genuine target structure from artefacts driven by admixture or heterogeneity in the reference population to obtain a coherent picture of demographic history.

### 5 Recent admixtures

We reconstruct Relate trees for the HGDP+1,000GP dataset, comprising 1,026 individuals and fit three cases with previously reported recent admixture [14, 19].

#### 5.1 Hazara

Hazara individuals from Central Asia exhibit clear signals of recent admixture between South and East Asian groups. Using GhostBuster, we decompose ancestry across chromosomes 1–5 for five Hazara individuals, focusing on coalescence events within the last 10,000 years and incorporating all other populations of the HGDP+1,000GP dataset as references. Our analysis reveals two components, each contributing approximately 50%. Component 1 aligns with East Asian groups, while component 2 is closer to South Asian groups (see Supplementary Figure **7**a). These components are well-separated in PC space (see Supplementary Figure **7**b), and model selection supports a two-component fit with high predictive accuracy (see Supplementary Figure **7**c). Coancestry curves estimate an admixture date of 26.2 generations or 733.6 years (see Supplementary Figure **7**d). Local ancestry inferred by GhostBuster correlates strongly with estimates from Mosaic, with a mean correlation of 0.87 (see Supplementary Figure **7**e), although minor differences may arise from Mosaic's phasing correction step.

#### 5.2 Bedouin

Bedouin individuals from the Levant have a well-documented history of European and African admixture within the last 40 generations. Using GhostBuster, we decompose ancestry across chromosomes 1–5 for five Bedouin individuals, focusing on coalescence events within the last 10,000 years and incorporating all other populations as references. Our analysis reveals two components: a major component (90%) closely aligned with Middle Eastern and European groups, and a minor component associated with sub-Saharan

African ancestry (see Supplementary Figure **8**a). These components are clearly distinguishable in PC space (see Supplementary Figure **8**b), and model selection supports this two-way structure (see Supplementary Figure **8**c). Coancestry curves estimate an admixture date of 36.6 generations, or approximately 1,024.8 years (see Supplementary Figure **8**d). Local ancestry inferred by GhostBuster is highly concordant with estimates from Mosaic, with a mean correlation of 0.86 (see Supplementary Figure **8**e).

#### 5.3 Maya

Maya individuals from Central America exhibit a well-documented history of two-way admixture between Native American and European groups, largely resulting from European colonisation. Using GhostBuster, we decompose ancestry across chromosomes 1–5 for five Maya individuals, focusing on coalescence events within the last 10,000 years and incorporating all other populations as references. Our analysis reveals two components: a major component (86%) closely aligned with other Native American groups, and a minor component associated with European ancestry (see Supplementary Figure **9**a). These components are clearly distinguishable in PC space (see Supplementary Figure **9**b), and model selection supports this two-way structure (see Supplementary Figure **9**c). Coancestry curves estimate an admixture date of 6.4 generations, or approximately 179.2 years (see Supplementary Figure **9**d). Local ancestry inferred by GhostBuster is highly concordant with estimates from Mosaic, with a mean correlation of 0.92 (see Supplementary Figure **9**e).

### 6 PRDM9-based analyses

#### 6.1 PRDM9 hotspots

We obtained genome-wide maps of PRDM9-A [20] and PRDM9-C [21] hotspots, comprising 25,440 and 19,366 autosomal hotspots, respectively. For PRDM9-C, hotspot coordinates were further refined by analysing published chromatin-immunoprecipitation sequencing of ssDNA (ChIP-SSDS) data for DMC1, generated from testes of a human male heterozygous for two C-like PRDM9 alleles (C and L4) [21, 22]. Hotspots were identified, and their intensities quantified using a peak-calling framework [23], and the most likely PRDM9-binding site within each hotspot was inferred using a motif-calling approach [24]. The midpoint of this binding site was taken as the hotspot centre.

PRDM9-A and PRDM9-C hotspots were analysed separately, and any region where both hotspot types occurred within a 5 kb interval was excluded. For GCbGC analyses, only hotspots above the median hotspot heat were retained. For hotspot-death analyses, we used the filtered PRDM9-C hotspots where we inferred hotspot centres.

#### 6.2 Testing of "hotspot death"

Both theory [25] and observed data for human [26] and mouse [26, 27] hotspots imply that mutations occurring within these motifs, and disrupting binding by a particular PRDM9 allele, are over-transmitted in offspring possessing that allele, resulting in a strong evolutionary advantage (drive) relative to neutrally evolving mutations. The primary mechanism for this "hotspot death" phenomenon is that PRDM9 will tend to bind intact motifs, causing double-strand breaks that are repaired using its homolog, which may carry disruptive mutations. This drive acts to eliminate hotspot motifs from genomes within populations where they are active, providing a complementary yet distinct signature to GCbGC acting at the same hotspots, that is highly population-specific [23] if hotspots differ between groups.

To identify hotspot death within PRDM9 binding motifs, we constructed 31-bp (PRDM9-A) and 32-bp (PRDM9-C) position-weight matrices (PWMs) based on base frequencies around the motif centre. Motif boundaries were defined using positions where the $\log_2$-entropy relative to a uniform base distribution exceeded 0.1, indicating sites where mutations may affect binding. The resulting PWMs are shown in Supplementary Figure **24**.

We identified all segregating mutations in our joint Vindija Neanderthal, Denisova and African individual (Modern Africans + archaic) variation data, which were at least 10-bp separated from the previous SNP (to avoid overcounting single motifs, or potential simultaneous mutations) and did not create or remove CG dinucleotides (where ancestral and derived allele calls might be unreliable as these sites are known to be highly mutable). Of these 11,328,164 SNPs, we then identified those overlapping a PRDM9-A or PRDM9-C binding motif and where the derived variant reduced the predicted PRDM9 binding score based on the PWM, resulting in 3,194 PRDM9-A and 1,071 PRDM9-C disrupting mutations. We also defined (genome-wide) potentially introgressed SNPs from HA into Neanderthal similar to our GCbGC analysis (see section 6.4) as SNPs whose estimated age is below 500,000YBP, with the derived allele present in both humans and Neanderthal (at least one copy) and where the SNP both maps to the nearby Relate tree and is not inferred to be allele-flipped (i.e., the chimp-inferred ancestral allele appears correct to Relate). For each of the 11.3M SNPs retained above, we obtained its upstream and downstream distance to the nearest such inferred "introgressed" SNP.

To test for evidence of drive favouring motif-disrupting mutations, we first calculated the "null" expectation and compared our observations against it. Separately for PRDM9-A and PRDM9-C, we computed this null by estimating mutation rates from nearby regions (100 bp–1 kb from each hotspot motif), accounting for base composition within motifs and the spectrum of motif-disrupting changes (i.e. separately estimating the rate of mutation from A/T and G/C bases to each other possible bases). Using these rates, we then predicted the expected ("null") number of motif-disrupting mutations across all frequency combinations $(i, j, k)$ in

Neanderthals, Denisovans, and humans ($i, j = 0, 1, 2$; $k = 0$–$362$), based on genome-wide mutation counts, noting that the majority of mutations occur outside hotspot regions. Although GC-biased gene conversion within hotspots might in principle bias motif frequencies, as a recombination-driven phenomenon we can view this as part of the signal within a motif, although because both the A- and C- alleles bind GC-rich motifs, this phenomenon is predicted to tend to act in the opposite direction to, and frequently more weakly than, motif death driven by PRDM9 binding impacts.

Based on these predictions, we compared the expected and observed numbers of PRDM9-A and PRDM9-C motif-disrupting mutations across human allele frequencies. Because hotspot death drives mutations towards fixation, it is expected to produce an excess of high-frequency derived mutations in affected populations. We therefore stratified variants by derived allele frequency (0%, 10%, ..., 100%) in humans and examined enrichment across these bins. In particular, we plotted observed divided by expected counts (Supplementary Figure **24**; human-only) and tested (1-sided) for excess mutations over expectations, using an exact Poisson $p$-value modelling null mutation counts as Poisson-distributed with the null expectation, and similarly obtaining 95% CIs for the ratio of observed vs. expected mean at each threshold. For both A- and C-alleles, this resulted in overwhelming evidence supporting motif death; > 4-fold expected mutation counts in the highest frequency bin ($p = 9.9 \times 10^{-6}$ for the C-allele and $p = 3.24 \times 10^{-27}$ for the A-allele motifs). All bins were significant for both motifs, with, as expected, an increasing enrichment with increasing derived allele frequency threshold.

Notably, the C-allele showed stronger enrichment than the A-allele at lower frequencies, with the opposite occurring at high-frequency mutations, consistent with a more recent onset of drive for the C-allele than the A-allele generating many mutations not yet nearing fixation. This supports previous work [28] suggesting the C-allele is derived, and the signal of GCbGC generating more fixed sites in hominins for the A than the C-allele (see section 6.3).

Given the very strong hotspot death signal in humans, we then refined our analysis to explore patterns in both archaic genomes, recalling that the GCbGC signal suggests activity of C-allele hotspots (but not A-allele hotspots) in a shared human-Neanderthal-Denisovan ancestral population, and of A-allele hotspots (and potentially not C-allele hotspots) in the human-related population that introgresses into Neanderthals. We note that the motif death signal is distinct from the former, which relies on SNPs outside the motif itself. Compared to the GCbGC signal, we expect a priori that (given a small mutational target), the hotspot death signal impacts a smaller number of variants, but potentially much more strongly, because of their direct impact on hotspot activity, meaning that such mutations might undergo rapid spread towards fixation, and so quite precisely reflect the timescales when particular hotspots are active.

We divided SNPs into those whose derived alleles are found only in humans, are shared (frequency> 0) between humans and one of the archaics (but not the other), or seen in both archaics and modern humans, resulting in four categories for each of A-allele and C-allele hotspot disrupting mutations (we discuss mutations seen only in Neanderthal and/or Denisova below). We abbreviate H (Human), N (Neanderthal), and D (Denisovan) so that, e.g. H-N SNPs are those where derived alleles are present in humans and the Neanderthal but not the Denisovan, etc. For each such category, we obtained observed versus expected ratios, 1-sided exact $p$-values of enrichment, and 95% confidence intervals for observed vs. expected activity, and plotted these against derived allele frequency in humans (Extended Data Figure 9). This revealed, as expected, a strong human-specific signal (with increased enrichment) for both A-allele and C-allele motifs (Extended Data Figure 9a). However, only two types of variant showed enrichment in the archaics, with results consistent across frequency thresholds: a very strong signal for H-N SNPs for A-allele motifs, but not C-allele motifs ($p = 1.3 \times 10^{-16}$ for the 90% cutoff, significant for all cutoffs, and with increasing enrichment for increased human frequencies; Extended Data Figure 9b; Supplementary Table **3**c), and enrichment for H-N-D SNPs shared among all three hominins for C-allele (Extended Data Figure 9c; $p = 0.017$ for all motifs, and $p = 0.012$ for motifs above 90% derived frequency in humans), but not A-allele motifs (which were instead depleted in this category). Thus, both motifs show drive signals in humans but distinct signals in archaics, which are fully consistent with the GCbGC signals.

As with GCbGC, the lack of any A-allele death signal shared between humans and Denisovans rules out introgression into the latter from a human-related population possessing the former. In contrast, partial sharing of hotspot death between humans and Neanderthals suggests that the latter might have acquired this signal from a shared ancestral group. Consistent with this, there is no signal for either motif for mutations not shared with humans ($p > 0.16$ for mutations seen only in N, D or N-D), implying minimal drive at these hotspots after their final split with humans. This might reflect a combination of lack of evolutionary time, activity of other hotspots, or small population sizes in the archaic groups. Similarly, Denisovans only share a significant C-allele drive signal at sites shared with both humans and Neanderthals, implying activity of the C-allele – but no evidence of A-allele activity – in the non-introgressed ancestral group "HC" (Figure 5c).

##### 6.2.1 Testing of "hotspot death" by evidence of HA introgression

In principle, two possible models might explain these patterns (and those for GCbGC).

1. A first possibility is rapid replacement of the C-allele in a shared human-Neanderthal-Denisovan ancestral population by the A-allele, and subsequent introgression into Neanderthal of the now A-allele-dominated human population. This seems a priori perhaps unlikely given the derived status of the C-allele, as well as the reasonably high frequency of the C-allele in these human populations today, but given the rapid evolution of PRDM9 in general is important to consider.
2. The second possibility is that the A-allele and C-allele drive and GCbGC instead occurred in distinct, deeply separated archaic populations HC and HA, as shown in Figure 5c and suggested by GhostBuster, with the latter group then splitting to form the main Neanderthal-Denisovan population, and the former group both admixing into Neanderthals, but also mixing with the second group to form (the ancestors of) modern-day humans as an admixed population.

In other words, the A-allele and C-allele "hotspot death" drive that has generated SNPs present in humans today either (1) occurred in a single evolving population but at different times, or (2) in different populations. The latter means that humans today are formed as an ancient admixture of these two distinct groups, allowing our species to possess both types of signal strongly, as observed.

These possibilities may be distinguished using patterns of hotspot-death enrichment stratified by admixture in Neanderthals. If the first possibility is true, haplotypes admixing into Neanderthals will carry the older signal of C-allele hotspot death from the earlier (ancestral) population in which this allele was active, so that the signal of H-N-D sharing should remain unaffected (or even become stronger, if some additional C-allele drive occurred in the descendant population) within segments introgressed from the human lineage. In contrast, if the second possibility is correct, then haplotypes introgressing into Neanderthals will carry only A-allele hotspot death, and not substantial C-allele death. Moreover, where introgression into Neanderthal occurs, the Denisovan genome will still typically inherit C-allele death mutations from the shared H-N-D ancestral population HC, and so in introgressed regions, C-allele disrupting mutations will be shared by Denisovans and Humans but not Neanderthals. Thus, non-introgressed regions will be enriched for H-N-D shared mutations, while introgressed regions show H-D mutation enrichment, at C-allele disrupting mutations.

To test this, we used a measure of potential introgression for each A-allele or C-allele motif (we note that it is not possible to use GhostBuster inferred ancestry robustly near recombination-prone hotspots, so as for our GCbGC analyses we employ a more elementary definition for this test): the distance to the nearest potential introgressed SNP (similar to GCbGC analysis described in section 6.4 and Figure 5h). These distances are expected to be reduced in introgressed regions, because they are more likely to contain such SNPs. To visualise signals, we plotted the median distance to the nearest introgressed SNP for each threshold, hotspot motif type and category of SNP used in the previous section. (Extended Data Figure 9d-f). The results suggest no particularly strong introgression signals for human-only motif-disrupting SNPs, and overall there is significant evidence of introgression into Neanderthals for only two categories of SNP: First, we see introgression at A-hotspot SNPs shared between humans and Neanderthals as expected for both hypotheses and from the GCbGC results, but second, also the Neanderthal shows a strong signal of introgression at C-hotspot SNPs shared between humans and Denisovans, but absent in Neanderthal. This observation means

Neanderthal appears to be introgressed when it does not possess C-hotspot disrupting mutations that are present in the other two hominins, a pattern expected under the second hypothesis only. Moreover, at C-hotspots, H-N-D fully shared disrupting mutations show a trend towards lacking introgression (longer median distance), again as expected only under the second hypothesis. We repeated the analysis, with an alternate definition of introgressed distance (maximum of the upstream and downstream introgressed distance for each hotspot) and obtained similar conclusions (Supplementary Table **3**d).

To formally test and obtain confidence intervals, we assessed whether the observed signals exceeded expectations under a matched null model that accounts for mutation type (e.g. human–Denisovan shared mutations absent in Neanderthals). Specifically, for each PRDM9-A or PRDM9-C motif-disrupting mutation, we randomly sampled 10 control mutations from the 5 kb surrounding regions (excluding motif-disrupting sites), matched for allele frequency in Neanderthals, Denisovans, and humans. Then, for SNPs in the categories discussed above (H-N shared SNPs for the A-allele, and H-N-D vs H-D shared SNPs for the C-allele), we examined whether SNPs in these categories showed departures relative to the control SNPs in their introgression measures (distance). To maximise power (available SNP numbers mean we ultimately test only 40 SNPs for the A-allele and 60 for the C-allele), we analysed all SNPs of frequency above that minimum human frequency giving the strongest signal of hotspot death enrichment (lowest $p$-value) in the above testing based on observed mutation counts (yielding $n > 351$ copies in Africans for the A-allele whose signal is dominated by high-frequency derived variants, and $n > 25$ copies for the C-allele). Because we condition on both the number and frequencies of mutations in the subsequent introgression testing, the choice of $n$ threshold does not impact the introgression test under the null: thresholding, though, usefully enriches for SNPs truly impacted by drive to eliminate hotspots. Finally, we simply used a non-parametric (1-sided) Wilcoxon rank sum test of whether A-allele H-N shared SNPs are nearer to introgressed SNPs than expectations from the matched controls. For the C-allele, if the second hypothesis is true then H-N-D distances should be larger than expected from null SNPs, while the opposite will hold for H-D distances. We therefore jointly tested the two impacted categories: H-N-D and H-D SNPs by first centring observed distances by subtracting the median distance in each category from all SNPs within 5kb, and sign-flipping the centred H-D values to contrast effect direction. We then again Wilcoxon-tested, with the combined vector, vs. the control SNPs processed identically and under a 1-sided positive alternative.

The results confirm that introgression drives the A-hotspot signal: $p = 0.016$ for the nearest distance, $p = 0.021$ for the maximum distance to likely introgressed SNP (Supplementary Table **3**d). Note that our test examined whether hotspot-disrupting mutations are more introgressed relative to other H-N shared SNPs. This result therefore means not only that this signal is dominated by likely introgression, but that this holds more strongly even than expected for H-N shared SNPs, likely because the A-allele hotspot death signal for SNPs in the H-N category is dominated by actual introgression, with little contribution (in contrast to the background region) from SNPs falling within this category simply due to incomplete lineage sorting. Thus, hotspot death at A-hotspots arises due to introgression from the HA population, similar to the GCbGC signal (Extended Data Figure 9).

For the C-allele, the test again yielded significant $p$-values for both tests: $p = 0.025$ for the nearest distance and $p = 0.035$ for the maximum distance, as suggested by Extended Data Figure 9e-f. Finally, control reciprocal testing (of C-allele hotspots but using the A-allele hotspot category H-N, and vice versa) yielded no significance ($p > 0.35$ for all 4 tests). Using a mutation frequency threshold of $n > 0$ instead of $n > 25/351$ also yielded similar, though slightly less significant results; Supplementary Table **3**d). Thus, hotspot death at C-hotspots specifically avoids appearing on the Neanderthal background in regions where Neanderthals are introgressed from the HA population, but remains shared between Denisovans and Humans, once again consistent with having occurred in the HC population, and with the GCbGC signal (Figure 5g). Again, we note that this pattern holds more strongly than expected for randomly selected mutations in the same categories. Therefore, this signal provides statistically significant and highly consistent (with the GCbGC signals, and with the overall pattern of motif death enrichment) evidence against the first hypothesis discussed above, and in close agreement with patterns predicted by the second hypothesis, where A- and C-allele hotspot death mainly occurs in separated populations.

##### 6.3 GCbGC signals on the human, chimp and gorilla lineages

We used the liftOver function within R to map between the human (H; hg38), chimpanzee (C; PanTro6) and Gorilla (G; Gorgor6) reference genomes. For each motif, we did this for the 5kb upstream and 5kb downstream of the motif centre, generating regions of reciprocal-best mapping and lacking indels to compare the genomes. Within the resulting 10kb regions, we called ancestral bases as a particular allele if all 3 species aligned there, and at least two of the three species agreed, after removing bases contained within a CpG dinucleotide in any species. Against these ancestral calls, we then identified, classified and counted mutations on particular lineages (H, C or G).

Separately for A-allele and C-allele hotspot motifs, and for 50bp bins tiled across the region after removing the central 50bp containing the motif itself, we counted mutations from A/T to G/C (W→S) and S→W mutations, normalising by the total number of called ancestral W or S bases within each region. We plotted the resulting signal of excess W→S mutations per weak base along the genome for each lineage (Supplementary Figure **23**), after subtracting background expectations obtained by averaging mutation rates 2.5–5kb from the motif. For both the A-allele and C-allele motif loci, this confirmed a signal of GCbGC specific to W→S mutations, and specific to the human lineage where these alleles are (and have been) active. Moreover, the A-allele signal is much stronger than the C-allele signal, consistent with it being the older/ancestral allele. No signal is apparent for the chimpanzee or gorilla lineages, or for mutations in the opposite direction (S→W).

For the central bin (corresponding to mutations 25–50bp away from the motif centre to exclude mutations within the motif, but otherwise examine nearby mutations), we obtained and plotted exact 95% Poisson confidence intervals for the excess mutations, and we also obtained and plotted (as horizontal lines) the excess non-fixed i.e. segregating variants within the H-N-D dataset, which contribute to the lineage count if the derived allele is present in the human genome sequence. For this, we weighted each central-bin SNP in our H-N-D data by its derived human frequency and again subtracted the background expectation using mutations 2.5–5kb from the motif; we upweighted the resulting estimates by the fraction of genome passing our masking filters in all 3 hominins (52%). The non-fixed mutation estimates for A-motif and C-motif hotspots are different, and we note that the C-motif estimate is similar to the total W→S excess seen on the human lineage, while the A-motif estimate is significantly below (but still ∼2/3 of) the total human-lineage W→S excess seen. Thus, all excess W→S signal for C-hotspots is statistically explained by variants not yet fixed in humans, supporting the idea that this allele arose only recently and has not had time to fix such variants, with the A-allele older, but with signals still mainly captured by mutations segregating among humans, Neanderthals and Denisovans today.

##### 6.4 Testing for GCbGC by evidence of HA introgression

To test whether hotspots nearby SNPs showing evidence of introgression from the archaic hominin population HA into Neanderthal are associated with archaic hotspot activity, we analysed the GCbGC signal at PRDM9 hotspots stratified by introgression strength around the hotspot. For the Chagysarskya Neanderthal (whose age is most similar to Denisovan) we identified all W→S and S→W mutations within 250bp of the estimated centre of each hotspot (after masking the central ±20bp region nearest the hotspot centre to avoid potential biases due to hotspot death as discussed above), which were present in at least 1 derived copy in the Neanderthal (later, we repeated this using equivalently defined Denisovan SNPs). For each such SNP, we annotated it by the distance to the nearest likely introgressed SNP, defined as the nearest SNP outside the hotspot (over 500bp from the nearest hotspot centre) and which (i) maps to the Relate genealogy and is not allele-flipped, (ii) has an estimated age below 500,000YBP, and (iii) has a derived allele present in both Neanderthal and African samples (in 3 or more copies in the latter, to remove likely back-to-Africa variants), but absent in Denisovans. Then, W→S and S→W mutations within hotspots were stratified into five quintiles using the distance annotation, so that hotspots that have introgressed from the HA population are expected to be over-represented within the lower bins, because SNPs within these bins are near a young SNP shared between humans and Neanderthals.

To construct a test, we used counts of W→S vs. S→W SNPs in each bin. Modelling these counts as binomial conditional on their total allows construction of an exact 95% confidence interval of the ratio of W→S to S→W mutations in each bin, expected to increase if ancient hotspot activity has occurred in Neanderthal ancestors, because recombination promotes W→S and helps to eliminate S→W mutations from the Neanderthal genome. In Figure 5h, we plot this ratio and CI's for both A and C hotspots. In these plots, we also show results applying the corresponding analysis for Denisovan-human shared SNPs absent in Neanderthals. We used this normalisation (slightly different from that used in Fig. 5f-g) to ensure robust testing: using a W→S vs S→W ratio rather than W→S counts avoids biases arising from changing total diversity levels (which would increase both W→S and S→W counts), preventing impacts simply from increasing/decreasing overall local diversity between bins (for example small distances might bias towards more SNPs nearby, generally). The corresponding ratios for Denisovan-human SNPs further provide a natural control for comparison among hominins at the identical hotspots, with e.g. the same base composition in each bin.

The results show that the Neanderthal-specific elevation of W→S mutations at PRDM9-A hotspots is strongly associated with introgression, and is significantly stronger in lower "introgression" bins, while not significantly greater for the last bin, which essentially reflects non-introgressed hotspots, than the average Denisovan signal (dotted line, which represents an appropriate null given that no PRDM9-A peak exists in Denisovans in Fig. 5f). No trend is apparent in Denisovan-human SNPs, consistent with their lack of signal overall, implying that introgression from the HA group fully explains the observed GCbGC signal at PRDM9-A hotspots in Neanderthals. We note that not all regions in the first bin are likely to be introgressed from HA, so this analysis is expected to underestimate the strength of the effect, but perhaps entries in the last bin are unlikely to be introgressed, and here the signal vanishes, implying that without introgression, the GCbGC signal in Neanderthals essentially vanishes.

For PRDM9-C hotspots, a higher ratio of W→S vs S→W is observed overall (Fig. 5h), consistent with a GCbGC signal observed in both Neanderthals and Denisovans (Fig. 5f-g), but this does not differ significantly from the equivalent baseline in any bin, show any clear trend across bins, or differ for any bin between Neanderthal and Denisovan. Therefore, it appears that GCbGC at PRDM9-C hotspots is similar in both Neanderthals and Denisovans, and not associated with introgression into Neanderthals.

#### 6.5 PRDM9 typing

We first downloaded data for PRDM9 alleles and human genotypes from [22]. To infer PRDM9 alleles in archaic hominins, we extracted sequence reads for the PRDM9 region using samtools for region chr5:23,526,326-23,527,898, using no mapping quality filter, because we expect such reads to map similarly well to multiple places within the ZF array. We use nine individuals: the Ust-ishim [29] human (as a positive control), Denisova [30], three high-coverage Neanderthal individuals we analyse elsewhere (Altai [31], Chagyrskaya [32] and Vindija [33]), three lower-coverage individuals [34] to add corroborating data, and finally an individual from Denisova cave (Denisova11) that has been inferred to be an F1 hybrid between Neanderthal and Denisovan hominins [35]. We then processed each individual identically, using a slightly modified (to account for the fact that in archaic genomes we do not obtain long paired-end reads) previously published approach [28] that has been applied and validated to work well in humans.

First, we took the collection of previously identified PRDM9 alleles, and identified the DNA sequence for each of 865 unique ZFs they contain (each 27 amino acids long, i.e. 84bp). Next, we selected the 65 alleles which were confirmed in [22], having been observed in other studies. Because PRDM9 alleles have arrays of often very similar or identical zinc fingers, we probabilistically sum over which particular finger the read may derive from. To compute this likelihood, we take the following steps:

For each read from a chosen hominin, we first identified the best mapping location within each of the 65 PRDM9 alleles. We did this by mapping each read to each possible site within the 84bp long ZF-array of a PRDM9 allele, scoring +1 for a matching base and −1 for a mismatching base. For each ZF-array, we chose the highest-scoring position. We retained only those reads mismatching at most 10 bases for their best-matching position and allele. We did not allow for indels because these are expected to produce frameshift errors, and in practice, reads containing indel errors are expected to be discarded.

For each retained read, we assumed that a read would always map at the same relative placement within a zinc finger and recorded the 'best' alignment to the zinc fingers by using the best-matching position for each allele relative to the start position of the corresponding zinc finger. This yielded a position between 1 and 84 for every PRDM9 allele. Next, we assumed that given a read aligned to some particular position within the array, it has a 1% probability $p_m = 0.01$ of mismatching (e.g. due to read errors) independently for each base. Suppose some read indexed $r$ has sequenced base $s_r^i$ at read positions $i = 1, 2, \ldots, l_r$ respectively. Suppose this read maps to position $x_r$ (between 0 and 83) within the ZF-array repeats.

We then obtain the likelihood for a given PRDM9 diploid genotype with alleles $I$ and $J$, which have $L_I$ and $L_J$ zinc fingers, respectively. Suppose allele $I$ has base $q_i^I$ at read positions $i = 1, 2, \ldots, L_I \times 84$ respectively. Then, assuming our read is equally likely a priori to come from any zinc finger and indexing zinc fingers by $g$, the likelihood of the observed sequence to come from allele $I$ is given by:

$$L(r, I) = \sum_{g=1}^{L_I} \prod_{i=1}^{l_r} (1 - p_m)^{\mathbf{1}_{s_r^i = q^I_{i+x_r+(g-1)\cdot 84}}} \, p_m^{\mathbf{1}_{s_r^i \neq q^I_{i+x_r+(g-1)\cdot 84}}}. \quad (62)$$

Bases mapping outside the zinc-finger array are trimmed from the read. In other words, we keep the position relative to the zinc fingers fixed, and sum over which particular finger the read may derive from to account for the fact that zinc fingers are often very similar or identical within PRDM9 alleles. Here, mismatching zinc fingers contribute little to the likelihood, but the score for an allele is increased if it contains several zinc fingers matching the read in question.

Because we analyse only reads from the array, to obtain the genotype likelihood we condition on mapping within the array – so that the probability of coming from some particular zinc finger is $1/(L_I + L_J)$ to yield:

$$L(r; I, J) = \frac{L(r, I) + L(r, J)}{L_I + L_J} \quad (63)$$

This scaling prevents, e.g. longer alleles from obtaining higher overall likelihoods. Finally, for an overall likelihood we multiply over all reads $r$ to yield

$$L(I, J) = \prod_r L(r; I, J). \quad (64)$$

We then obtain a matrix of likelihoods for all 65 × 65 (ordered) possible genotypes at PRDM9, and also obtain a posterior probability of each genotype using a uniform prior (given the allele frequencies are unclear in the past and other hominins). In practice, high-coverage individuals possessed likelihoods often dominated by one or a few allelic combinations, outweighing the impact of these prior beliefs.

We report in Supplementary Table **3**b: (i) the allele combination yielding the highest likelihood for an individual, (ii) after ordering allele combinations from highest to lowest likelihood, the set of possible alleles within the 95% highest posterior density credible set, (iii) the likelihood drop of two particular genotypes A/A and M10/M10 that we observe in several Neanderthals (A/A is also the commonest combination in humans), (iv) the expected number and probability of 1 or more A-type alleles similar to PRDM9-A (and so not PRDM9-C) using the classification published in [22, 28].

Because PRDM9 is rapidly evolving and the zinc-finger array is highly diverse, we also considered the possibility of new alleles that might occur in each hominin by performing further analyses of reads. First, using the $x_r$ values for each read, we asked whether the segments of each read overlapping distinct zinc fingers showed a strongest match (fewest mismatches) to known zinc finger(s) found in some PRDM9 allele, but not found within the ZF-array of the most likely allelic combination for this individual. We then flagged cases where at least 3 separate reads indicated an identical unexpected zinc finger, as evidence for such a zinc finger being incorporated into the array and indicating a new type. For each read, we next conservatively repeated the same analysis, but now ignored differences with expected ZFs due to "C" to "T" transitions, which might simply result from deamination of ancient DNA. We highlight resulting unexpected ZFs that cannot be explained by such deamination.

Finally, we identified mutations that might produce potential new ZFs (not observed in human surveys). For each read, we can calculate from each term in equation 62 and 63 the probability that the read comes from a particular zinc finger and allele within the most-likely genotype for the individual. Summing over reads, we obtained the expected number of sequenced bases of each type (A, C, G or T) within each position of each ZF. After filtering to positions where the expected read counts are at least 4, and where the same is true for a position at least 10bp upstream and 10bp downstream of that base (resulting in well-covered parts of the array), we identified those positions where the most-likely base differed from that expected for the allele as potential mutations, and report (Supplementary Table **3**b) such identified mutations, and whether they correspond to synonymous or non-synonymous changes.

We first compare results to those of a previous study [36] that analysed two of our samples: the Altai Neanderthal and Denisovan, using a different approach aiming to simply identify pairs of zinc-fingers (e.g. I-J) found in the corresponding individual. Our results agree closely for both cases. We find the Altai Neanderthal to be likely homozygous for PRDM9-A, which was identified in the previous study to be the most similar allele. However, unlike the previous study, we do not see clear evidence of an unexpected zinc-finger (S) in the Neanderthal, because although a small number (3) of reads match this allele, these can be readily explained by deamination and otherwise match expected zinc fingers within PRDM9-A. Because [36] also found a smaller number of reads supporting this zinc finger than others, we suspect the Altai Neanderthal is in fact indeed homozygous for PRDM9-A rather than carrying a new type.

For Denisovan, we infer (Supplementary Table **3**b) a combination of two distinct alleles, one of the PRDM9-C type, for example, the "Baudat_I" PRDM9 allele, and the other unclear but perhaps similar to PRDM9-B. However, we also observe synonymous changes (in almost all mapped reads) in the second and penultimate zinc fingers, and the presence of an unexpected zinc-finger type (R) most likely in multiple copies, not explained by deamination and flagged by six separate reads. Therefore, we believe both copies might be distinct from published human PRDM9 alleles, with some PRDM9-C like properties. The implied pairs of zinc fingers from [36] are all found within our predicted alleles, and the two non-synonymous mutations, as well as the "R" ZF-type, were also found in that work, so our results agree essentially perfectly for this individual. We are therefore able to conclude that the Denisovan possesses at least one "C-type" PRDM9 allele.

For other individuals, our results for the Ust-Ishim human suggest two different PRDM9-A type alleles carried, perhaps PRDM9-A itself and a distinct allele. For the Vindija Neanderthal, we inferred with high confidence a different PRDM9-A type allele, called M10, on both haplotypes. M10 differs from PRDM9-A by a single, synonymous change that is not expected to impact its binding properties but introduces the "Q" zinc-finger in place of the penultimate "I" zinc-finger. There is overwhelming evidence that this Q zinc-finger is present in this and also in other Neanderthals, with the Les-Cottes Neanderthal also most likely to be homozygous for M10. The Chagyrskaya Neanderthal potentially is homozygous for the same alleles (Supplementary Table **3**b) and almost certainly has at least one allele carrying a Q zinc-finger. The low-coverage Goyet and Mezmaiskaya individuals also have one of their most likely alleles carrying this finger. Finally, the likely F1 Neanderthal/Denisovan individual Denisova11 is inferred to carry the Q-type L24 PRDM9 allele and an unknown second allele that may be mutated relative to humans (Supplementary Table **3**b). Plausibly, these alleles might therefore come from the respective Neanderthal and Denisovan parents.

Overall, our data imply that all Neanderthals examined, other than the Altai, most likely possess at least one Q zinc-finger carrying allele.

##### 6.5.1 PRDM9 typing and Neanderthal-to-human introgression

We next examined the genotypes of modern humans with introgression from Neanderthals at PRDM9. Because we lack samples overlapping those previously typed at PRDM9, we downloaded genotypes for two SNPs, rs35502065 and rs79704982, from the Ensembl website (ensembl.org), which tag GhostBuster-inferred Neanderthal introgression at PRDM9. One SNP lies upstream, and the other downstream of the PRDM9 gene, and both showed strong linkage ($R^2 > 0.8$) with Neanderthal ancestry in the analysed non-African populations (Supplementary Figure **22**).

To investigate the relationship between the Q-type zinc finger (ZF) and Neanderthal-to-human introgression, we analysed modern individuals from the 1000 Genomes Project who were genotyped at two loci tagging Neanderthal ancestry and also had PRDM9 typing available. We examined non-African and African populations separately.

Among non-Africans (PJL, TSI, FIN, CHB), 25 individuals carried at least one copy of rs79704982. Of these, 24 also carried at least one Q-type zinc finger (ZF), accounting for 24 of the 27 Q-type carriers in these populations (OR = 2750, $p < 10^{-35}$, Fisher's exact test). This indicates an almost complete association between the introgressed haplotype and Q-type *PRDM9* alleles. A similar but slightly weaker association was observed at the other SNP (OR = 82, $p < 10^{-19}$). Q-type carriers were observed in PJL, TSI, and FIN, but were absent in CHB, and predominantly carried the L20 ($n = 10$), L24 ($n = 8$), and M10 ($n = 7$) alleles. Notably, L24 and especially M10 are also observed in Neanderthals based on direct typing.

In African populations (YRI and LWK), the association between the introgressed SNPs and Q-type ZF was weaker. Among Q-type carriers, 2 individuals carried rs79704982, while 8 individuals did not, corresponding to a weaker but still significant association (OR = 42, $p = 0.0069$), likely reflecting back-to-Africa migration. The two individuals carrying both rs79704982 and a Q-type ZF possessed the M10 and L23 alleles, respectively. In contrast, 7 of the 8 Q-type carriers without rs79704982 carried the L19 allele, a C-type allele in which the Q-type ZF is inserted near the end of the array; the remaining individual carried M10.

These results suggest that the Q-type zinc finger likely arose in a population ancestral to both humans and Neanderthals, but that specific alleles carrying this motif, particularly M10, entered modern humans via Neanderthal-to-human introgression. The near-complete association between rs79704982 and Q-type alleles in non-Africans indicates that this introgressed haplotype has been maintained and expanded outside Africa, while its weaker and more heterogeneous presence in Africa is consistent with more recent back-migration.

Functionally, this introgression has consequences for recombination in humans. Previous work [37] has shown that introgressed PRDM9 alleles can alter hotspot usage. While M10 is likely functionally similar to the A allele, differing only at a synonymous position, L20 and L24 carry nonsynonymous changes in their ZF arrays. In particular, L24 (which we think is a subsequent diversification of M10) activates hotspots at MSTM1A and MSTM1B that are not active in A-allele carriers, whereas L20 (and the related Q-type allele L13) suppresses A-allele hotspots and activates MSTM1B but not MSTM1A. Thus, these Neanderthal-derived PRDM9 alleles not only modify the human recombination landscape but also generate distinct hotspot profiles relative to one another.

#### 6.6 Testing for large introgressed regions genome-wide

We tested whether large regions of the genome might have evidence of excessive introgression from the human-related introgressing HA population, relative to the genome as a whole. This is expected to occur if introgressed alleles were advantageous, because such alleles would rapidly spread towards or to fixation in Neanderthals, creating a selective "sweep" signal carrying introgression over its span, and thereby generating an unusually large region within the Neanderthal genome almost fully introgressed. Thus, identified large introgressed regions represent candidates for selection favouring introgression.

To identify such regions, we considered three measures, based on the signal of introgression of recent coalescence between humans and Neanderthals (but not Denisovans), producing mutations shared between the human and Neanderthal but not with Denisovans. First, we used the inferred probability of introgression from our GhostBuster inference for each tree, weighted by tree span. Second, we constructed a more basic measure: the mean coalescence time inferred by Relate between humans and Neanderthal (for most of the genome, both Neanderthal alleles coalesce simultaneously with humans, but otherwise we use the lower time). Finally, we considered a measure independent of our tree inference: simply the fraction of segregating SNPs that are shared (some individual carries a derived allele) between humans and Neanderthals but absent in the Denisovan. We averaged each such measure across 500kb intervals tiled across the genome and centred at 500kb positions $(0, 500, 1000, \ldots$ kilobases). We filtered out regions with high levels of masking: where the total distance spanned by Relate trees was $< 50\%$ of the interval, or where the total SNP density was in the bottom 5% of all regions, yielding unfiltered regions spanning 2.389 gigabases of the genome. We obtained each measure for each Neanderthal separately (so there is an identical sample size $n = 1$ Neanderthal and Denisovan, and $n = 181$ African individuals) and then averaged across the three Neanderthals to provide a vector of each statistic. Then, we ranked and standardised (subtracting its mean and dividing by its standard deviation) these statistics. We obtained a single "overall" statistic by simply averaging the standardised statistics, after sign-flipping the standardised mean coalescence time because we expect this to decrease in the presence of admixture, while the others should increase. This equally weights each statistic; the resulting statistic has mean zero, but because the measures are correlated, non-unit variance, so we rescaled to generate a single measure with mean zero and variance 1 along the genome, and plotted this to generate Figure 5e, alongside its overall distribution. An overall $p$-value is given by the empirical rank $p$-value of the resulting 4778 regions. Although we only use these empirical $p$-values, we note that approximately 95% of this distribution lies between 2 standard errors of the mean, and the statistic shows a reasonable fit to a normal distribution (Figure 5e), as might be expected given each 500kb region contains many SNPs and often distinct coalescent trees too.

We list the top 20 identified regions (Supplementary Table **3**a) and their ranks and values for each statistic, as well as their ranking if instead of the above we used an alternative measure of the mean ranks of our 3 statistics.

If regions are truly introgressed, we expect, based on our other analyses, that alongside a decrease in the mean human-Neanderthal ancestral coalescence time, we ought to see an increase in the mean Neanderthal-Denisovan coalescence time, and also in the overall fraction of SNPs that differ between Neanderthal and Denisovan. We therefore calculated and checked these statistics across the same 500kb regions as above, calculating them as described above (and again averaging across Neanderthals, etc.).

By either ranking (combined statistic, and average ranks of the three separate statistics), the same region showed the strongest evidence of introgression: Chromosome 5, midpoint 23.5Mb. This 500kb region contains only a single gene, PRDM9. Based on the GhostBuster results alone, this region is the 4th highest scoring in the genome, and is the 6th highest and 18th highest using the fraction of shared SNPs and Neanderthal-human ancestor coalescence time (the ranking varies slightly by statistic used, though overall concordance is strong; Supplementary Table **3**a). This region and several other high-scoring regions also show evidence of an increase in Neanderthal-Denisovan coalescence time ($p = 0.02$) while the Denisovan-human ancestor coalescence time is not unusual ($p = 0.54$), highly consistent with this truly representing admixture.

#### 6.7 Alternate approach for estimating human-like introgression fraction in Neanderthals

For an alternate approach to estimate the introgression fraction, we use all three Neanderthals and leverage the fact that there is strong admixture at PRDM9. We concatenated the three Neanderthals, then averaged, for the genome as a whole and for the likely introgressed region around PRDM9 spanning 23.5Mb$\pm$250kb, the mean TMRCA between Human-Neanderthal (H-N), Human-Denisovan (H-D) and Neanderthal-Denisovan (N-D). Genome-wide means for H-N, H-D and N-D times were respectively 538KYA, 567KYA and 895KYA, versus values around PRDM9 of 305KYA, 603KYA and 1.403MYA. Thus, nearby PRDM9, the H-N coalescence time is markedly reduced while the N-D coalescence time increases, and the H-D coalescence time changes little, all as expected from our genome-wide introgression findings. We can use this to estimate the genome-wide introgression fraction, assuming, from our other analyses, that this region nearby PRDM9 is in fact 100% introgressed (otherwise our estimate would be increased) in all the Neanderthals. If the mean H-N coalescence time in introgressed regions is reduced by this estimated factor $305/603 = 0.505$, while outside introgressed regions it is equal to the mean of the H-D time, then the admixture fraction $\lambda$ satisfies the equation:

$$538 = [\lambda \times 0.505 + (1 - \lambda)] \times 567$$

Solving this yields the estimate $\hat{\lambda} = 0.104$, again suggesting an introgression fraction of around 10%. Substituting in this value further estimates that the mean Neanderthal-Denisovan coalescence times in non-introgressed and introgressed regions are equal to 837KYA and 1.403MY, respectively, a 68% ($\sim$570KYA) increase in divergence time.

### 7 Polygenic score portability analysis

We applied the ANCHOR analysis pipeline to quantify polygenic score (PGS) portability conditional on local ancestry. Specifically, we focused on two ancestry components identified using GhostBuster: OOA-like and non-OOA-like.

To assign OOA-like and non-OOA-like local ancestry to African-ancestry samples in the UK Biobank, we first identified their closest genetic matches within the HGDP + 1000 GP dataset and transferred the corresponding local ancestry calls. UK Biobank SNP array data were phased using SHAPEIT4 [38]. We retained 4,557 African-ancestry individuals with less than 10% recent European admixture, as inferred using `hapmix` [39, 40]. Phased haplotypes were lifted over to the hg38 genome build.

We applied an optimised implementation of ChromoPainter [41], using African samples from HGDP and 1,000 Genomes as the reference panel. Parameters were optimised using an initial fitting procedure on a subset of the data, yielding a mutation probability per SNP of $\mu = 0.0011$ and a recombination scaling factor $\rho = 368.43$. For each genomic position, ChromoPainter assigned posterior copying probabilities from reference haplotypes. For each site, we retained the top 20 reference individuals with the highest posterior probabilities and renormalised these probabilities to sum to one.

We estimated OOA-like and non-OOA-like local ancestry in UK Biobank samples by applying GhostBuster to the HGDP + 1,000 Genomes reference panel. Local ancestry at each site was computed as a weighted average of reference local ancestry posteriors, with weights given by ChromoPainter copying probabilities. This yielded local ancestry estimates for both ancestral components across 4,557 phased African-ancestry individuals.

#### 7.1 Polygenic score construction

Posterior mean effect size estimates were obtained using Quickdraws [42] for 32 quantitative traits in the UK Biobank (25 independent traits based on phenotypic $R^2 < 0.2$), trained on up to $\sim$ 405,000 white British individuals. Traits were selected based on predictive performance in Europeans ($R^2 > 0.1$) and high phenotyping rate ($> 0.9$) in self-identified Africans (Supplementary Figure **18**). Across several traits, predictive performance in African-ancestry individuals was approximately 20% of that observed in Europeans (Supplementary Figure **18**).

To assess ancestry-specific predictive ability, we decomposed the PGS of each African individual into OOA-like and non-OOA-like components following the ANCHOR framework [39].

Let $H_{ij}$ denote the phased haplotype of individual $i$ at variant $j$. Let $L_{ij}$ denote the local ancestry at that position. For ancestry $c \in \{\text{OOA}, \text{nonOOA}\}$, the expected haplotype-specific allele count is:

$$H_{ij}^c = H_{ij} \cdot P(L_{ij} = c), \tag{65}$$

and the expected diploid genotype is:

$$G_{ij}^c = H_{i_1 j}^c + H_{i_2 j}^c. \tag{66}$$

Following [39], genotypes were mean-centred before constructing ancestry-specific PGS. Define

$$P_{i,j}^{c_1,c_2} = P(L_{i_1 j} = c_1) \cdot P(L_{i_2 j} = c_2), \tag{67}$$

and let $f_j^{\text{OOA}}$ denote the allele frequency in the OOA-like background. This frequency was estimated via OLS regression of observed genotypes:

$$G_{ij} = I_j + S_j \left(2P_{ij}^{\text{OOA,OOA}} + P_{ij}^{\text{OOA,nonOOA}}\right) + \varepsilon_i, \tag{68}$$

with

$$f_j^{\text{OOA}} = \frac{I_j}{2}, \qquad f_j^{\text{nonOOA}} = S_j + \frac{I_j}{2}. \tag{69}$$

Mean-centred genotypes were then defined as:

$$\tilde{G}_{ij}^{\text{OOA}} = G_{ij}^{\text{OOA}} - 2f_j^{\text{OOA}} \left(P_{ij}^{\text{OOA,OOA}} + P_{ij}^{\text{OOA,nonOOA}}\right), \tag{70}$$

$$\tilde{G}_{ij}^{\text{nonOOA}} = G_{ij}^{\text{nonOOA}} - 2f_j^{\text{nonOOA}} \left(P_{ij}^{\text{nonOOA,nonOOA}} + P_{ij}^{\text{OOA,nonOOA}}\right). \tag{71}$$

Ancestry-specific PGS values were computed as:

$$\text{PGS}_i^{\text{OOA}} = \sum_j \beta_j \tilde{G}_{ij}^{\text{OOA}}, \qquad \text{PGS}_i^{\text{nonOOA}} = \sum_j \beta_j \tilde{G}_{ij}^{\text{nonOOA}}, \tag{72}$$

where $\beta_j$ denotes the posterior mean effect size for variant $j$.

To estimate ancestry-specific predictive effects, we fit the linear model:

$$Y_i = \alpha + \gamma_{\text{OOA}}\,\text{PGS}_i^{\text{OOA}} + \gamma_{\text{nonOOA}}\,\text{PGS}_i^{\text{nonOOA}} + \mathbf{C}_i^{\top}\boldsymbol{\theta} + \varepsilon_i, \tag{73}$$

where $\mathbf{C}_i$ denotes covariates, which include Age, Sex, Age$^2$, Sex$\times$Age, Sex$\times$Age$^2$, Smoking status, Site and PC$1-20$. The coefficients were estimated using ordinary least squares, with confidence intervals obtained from 1,000 bootstrap resamples. Effect sizes were normalised relative to those estimated in non-British self-identified European individuals, providing a measure of predictive performance relative to Europeans.

### Supplementary Figures

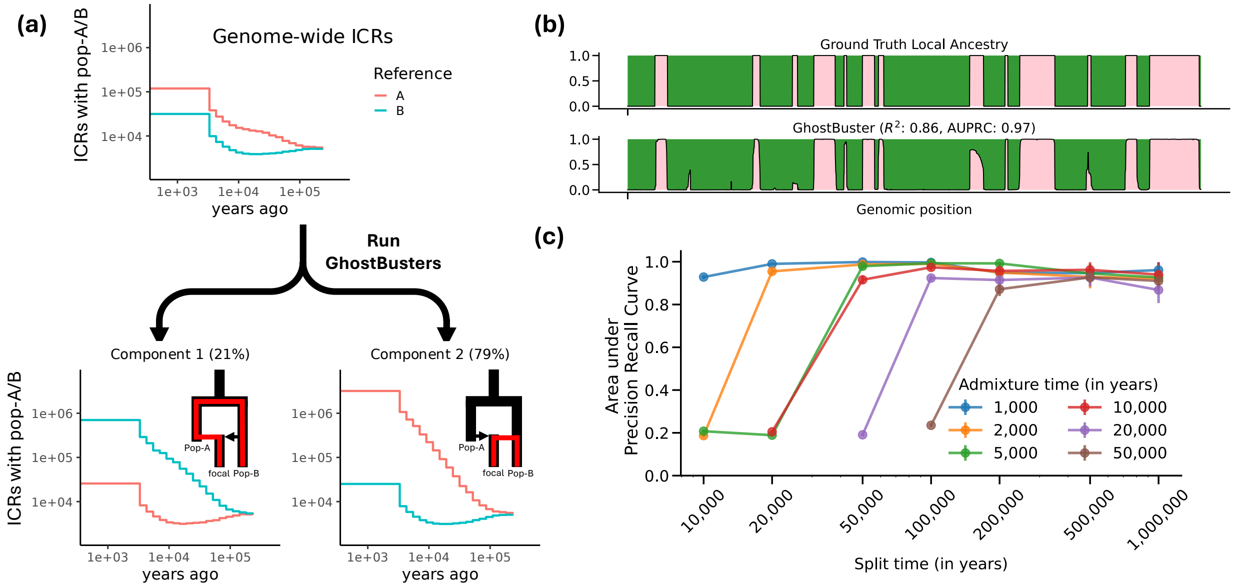

**Supplementary Figure 1**: **Performance of GhostBuster in the two-way admixture simulations with access to both source populations.** (a) Inferred genome-wide coalescence rates (ICRs) and admixture proportions for each component. Coalescence rates are between the focal individual and reference populations (Pop-A and Pop-B). (b) Comparison between ground-truth and inferred local ancestry along a simulated chromosome, demonstrating accurate recovery of ancestry segments ($R^2 = 0.86$, AUPRC = 0.97). (c) Performance across simulations with varying split and admixture times, quantified by area under the precision–recall curve (AUPRC). Error bars represent mean ± 95% confidence intervals calculated from 100 jackknife replicates along the genome.

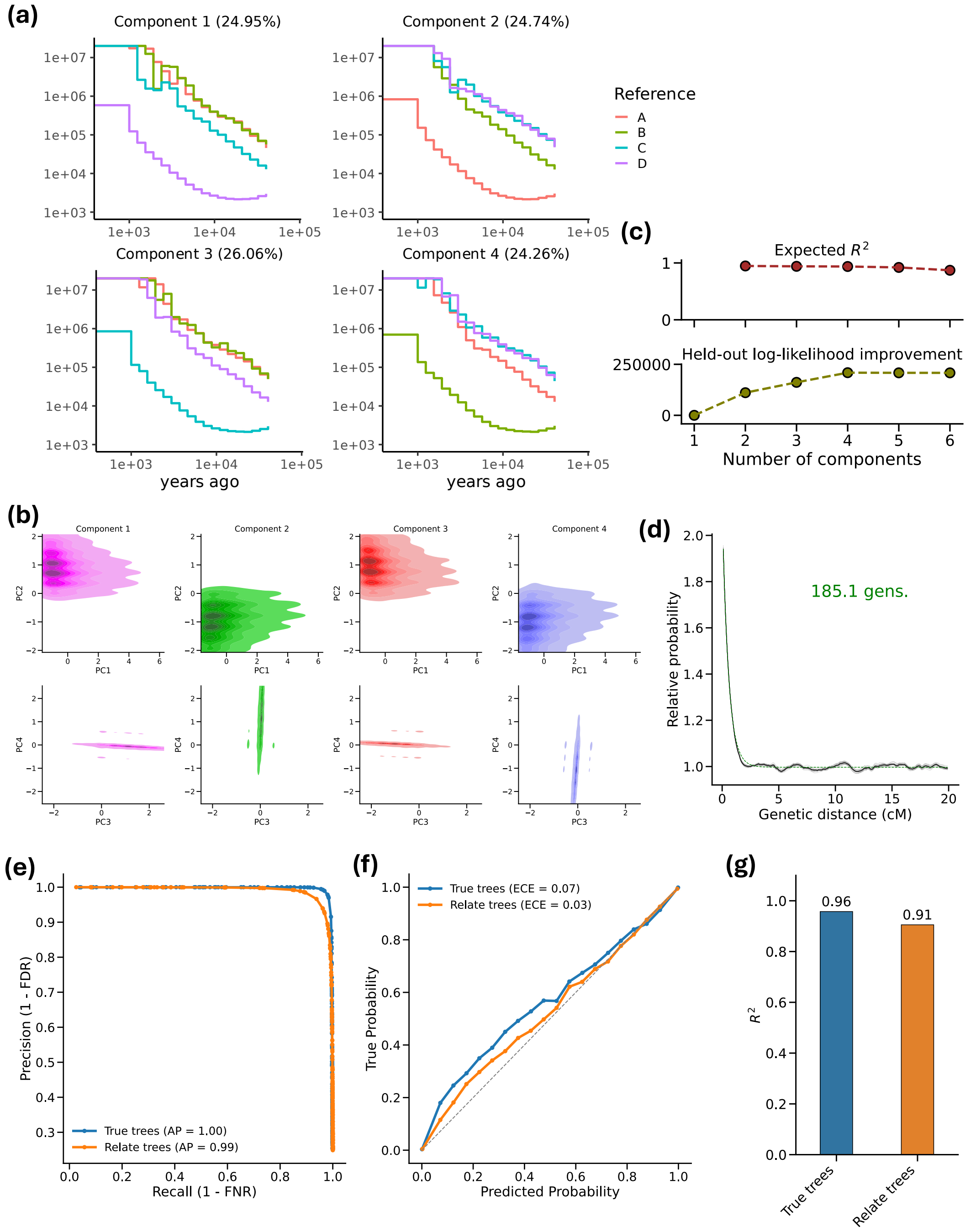

**Supplementary Figure 2**: **Decomposing focal individuals in the four-way admixture simulation with the presence of all source populations.** (a) Inferred inverse coalescence rates with a reference population and proportions. (b) PCA visualisation of the coalescence count and opportunity matrix derived from the genealogies plotted separately for each component. (c) Expected coefficient of determination and held-out log-likelihood improvement with varying number of components. (d) Coancestry curve for normalised joint probability of component 1 and the inferred admixture dates (in generations). (e) Precision-recall curves, (f) Local ancestry posterior calibration curves, and (g) prediction $R^2$ for inferred local ancestry compared to the simulated ground truth, where inference was done using true trees or Relate trees. The PCA visualisation in (b) is based on a KDE plot with a threshold of 0.05, and binary local ancestry estimates are obtained by thresholding the inferred posteriors at 0.5.

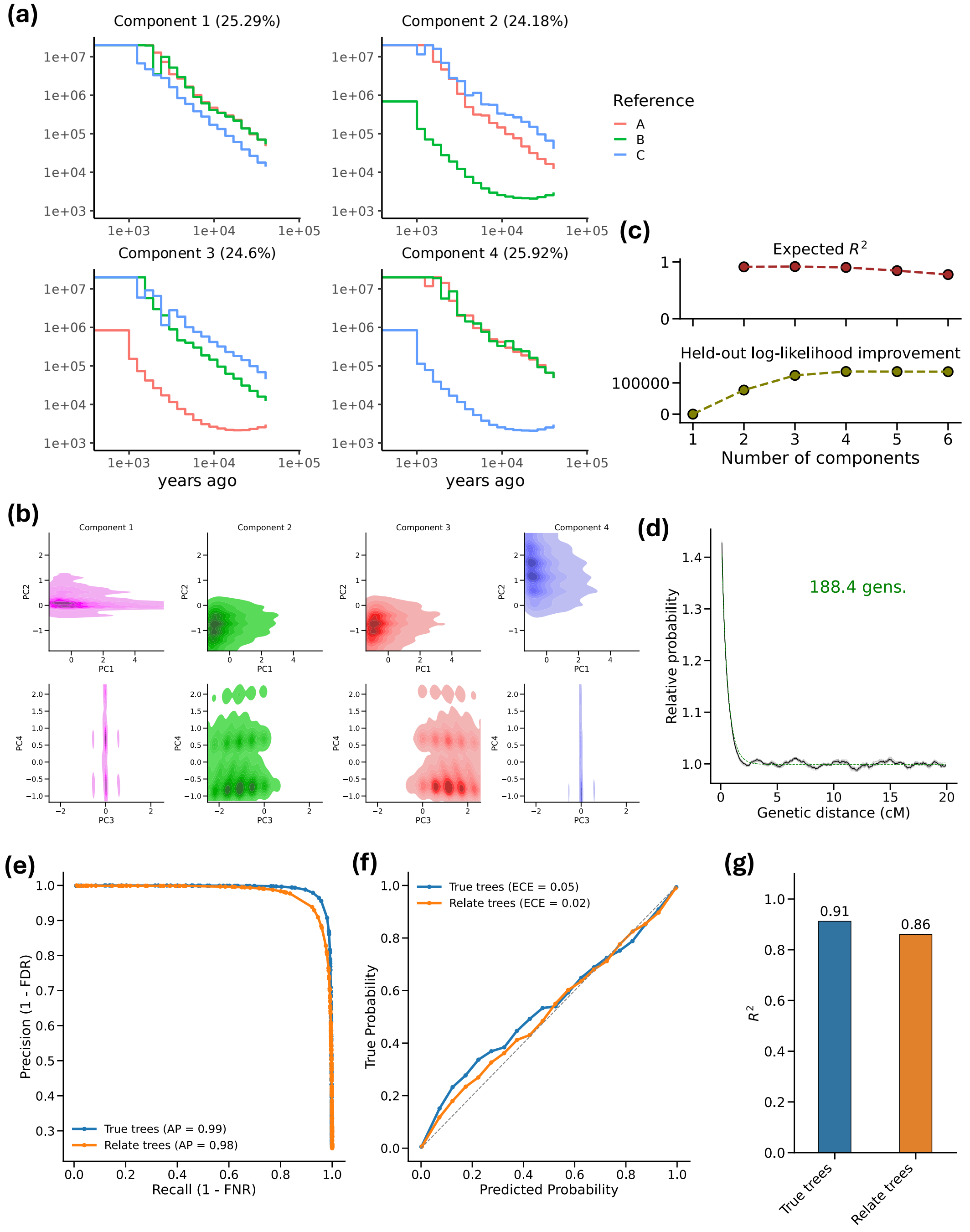

**Supplementary Figure 3**: **Decomposing focal individuals in the four-way admixture simulation with absence of one source population.** Analogous to Supplementary Figure **2**, we show (a) inferred inverse coalescence rates, (b) PCA visualisation, (c) expected coefficient of determination and held-out log-likelihood improvement with varying number of components, (d) coancestry curve, (e) precision-recall curves, (f) local ancestry posterior calibration curves, and (g) prediction $R^2$ for inferred local ancestry.

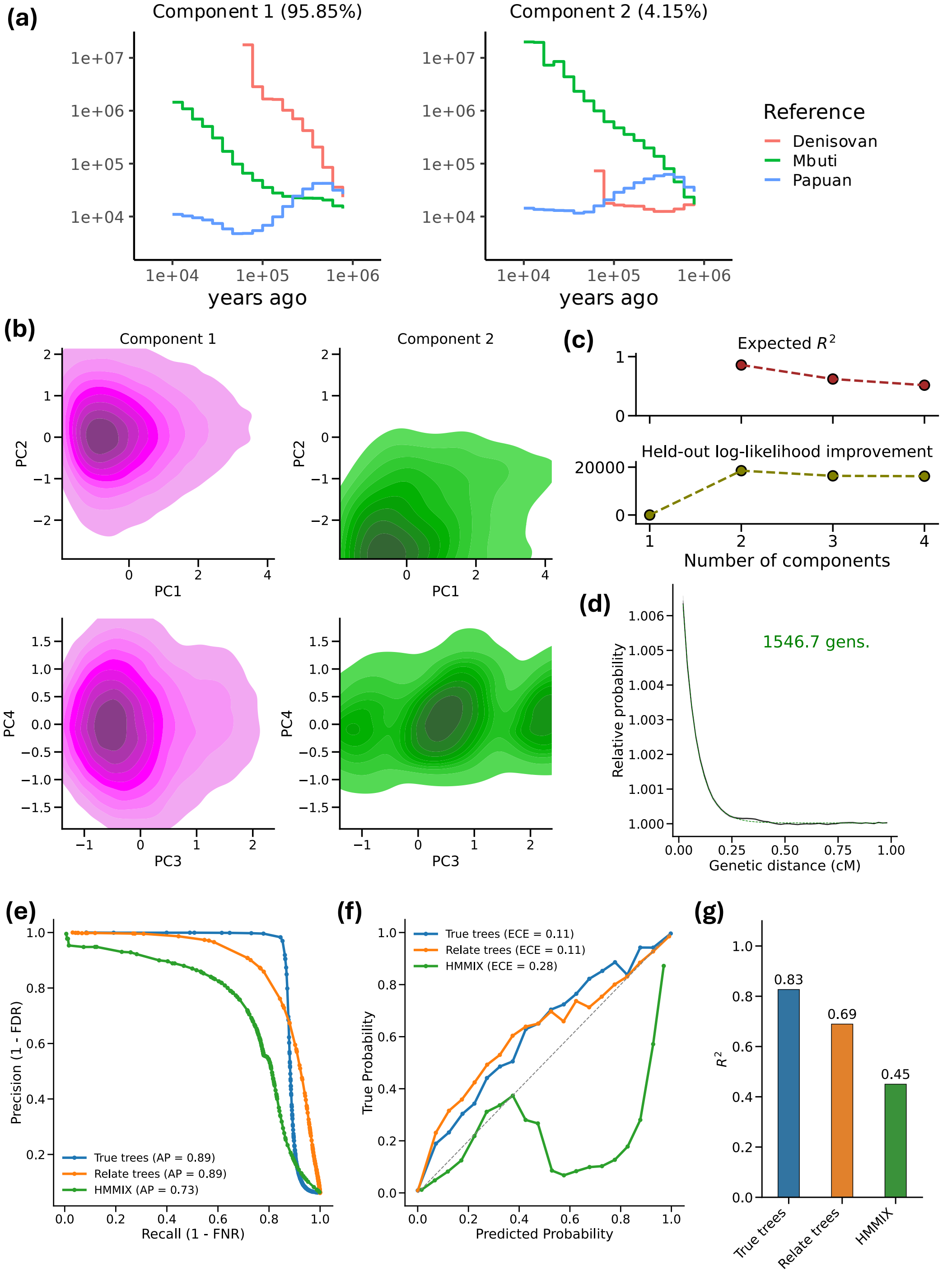

**Supplementary Figure 4**: **Decomposing Papuan-like individuals in simulations with Mbuti-like and Denisovan-like populations as additional references.** Analogous to Supplementary Figure **2**, we show (a) inferred inverse coalescence rates, (b) PCA visualisation, (c) expected coefficient of determination and held-out log-likelihood improvement with varying number of components, (d) coancestry curve, (e) precision-recall curves, (f) local ancestry posterior calibration curves, and (g) prediction $R^2$ for inferred local ancestry. We additionally compare to HMMIX [18].

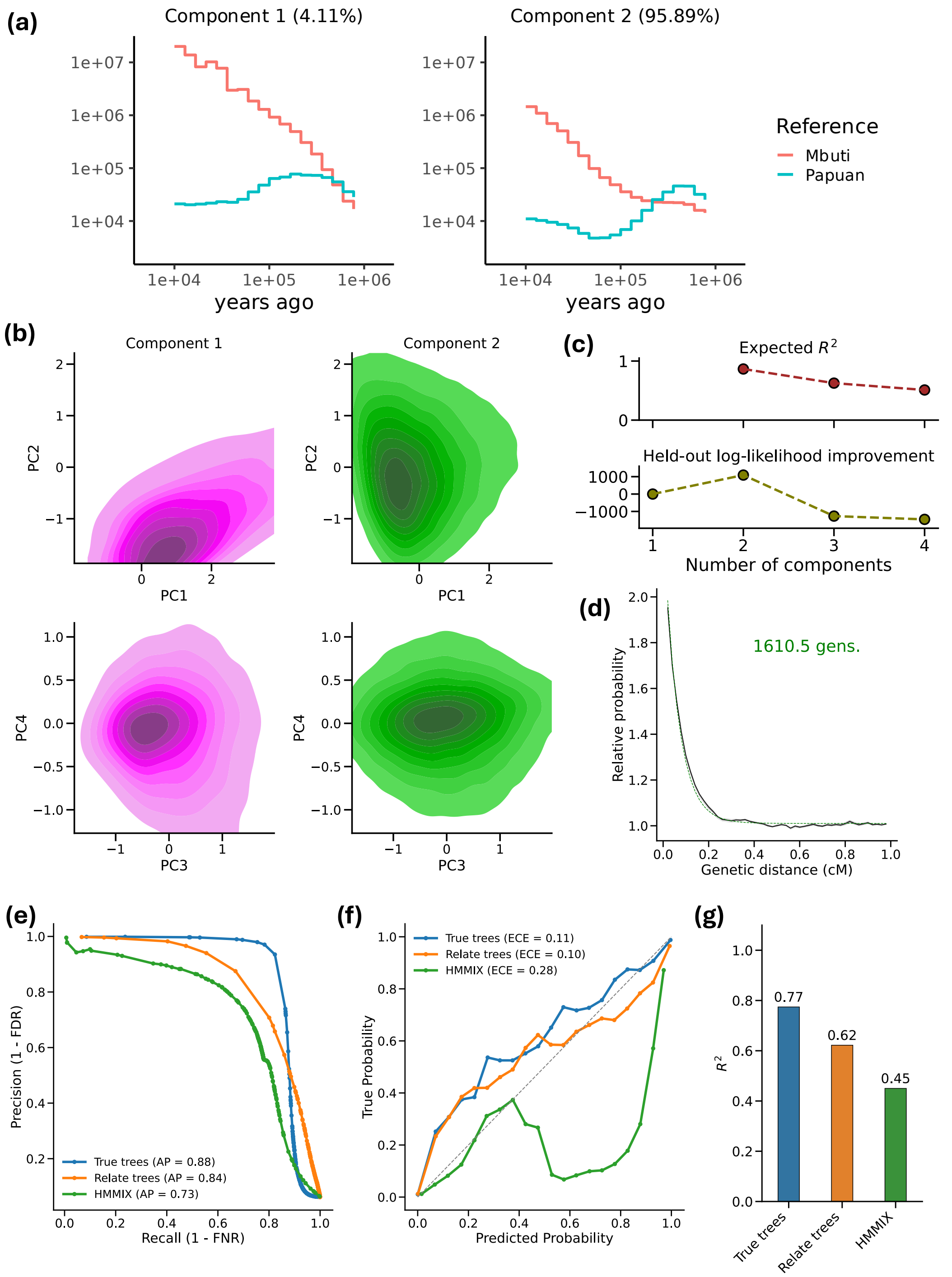

**Supplementary Figure 5**: **Decomposing Papuan-like individuals in simulations with only an Mbuti-like population and no Deniosvan-like population as additional references.** Analogous to Supplementary Figure **2**, we show (a) inferred inverse coalescence rates, (b) PCA visualisation, (c) expected coefficient of determination and held-out log-likelihood improvement with varying number of components, (d) coancestry curve, (e) precision-recall curves, (f) local ancestry posterior calibration curves, and (g) prediction $R^2$ for inferred local ancestry. We additionally compare to HMMIX [18].

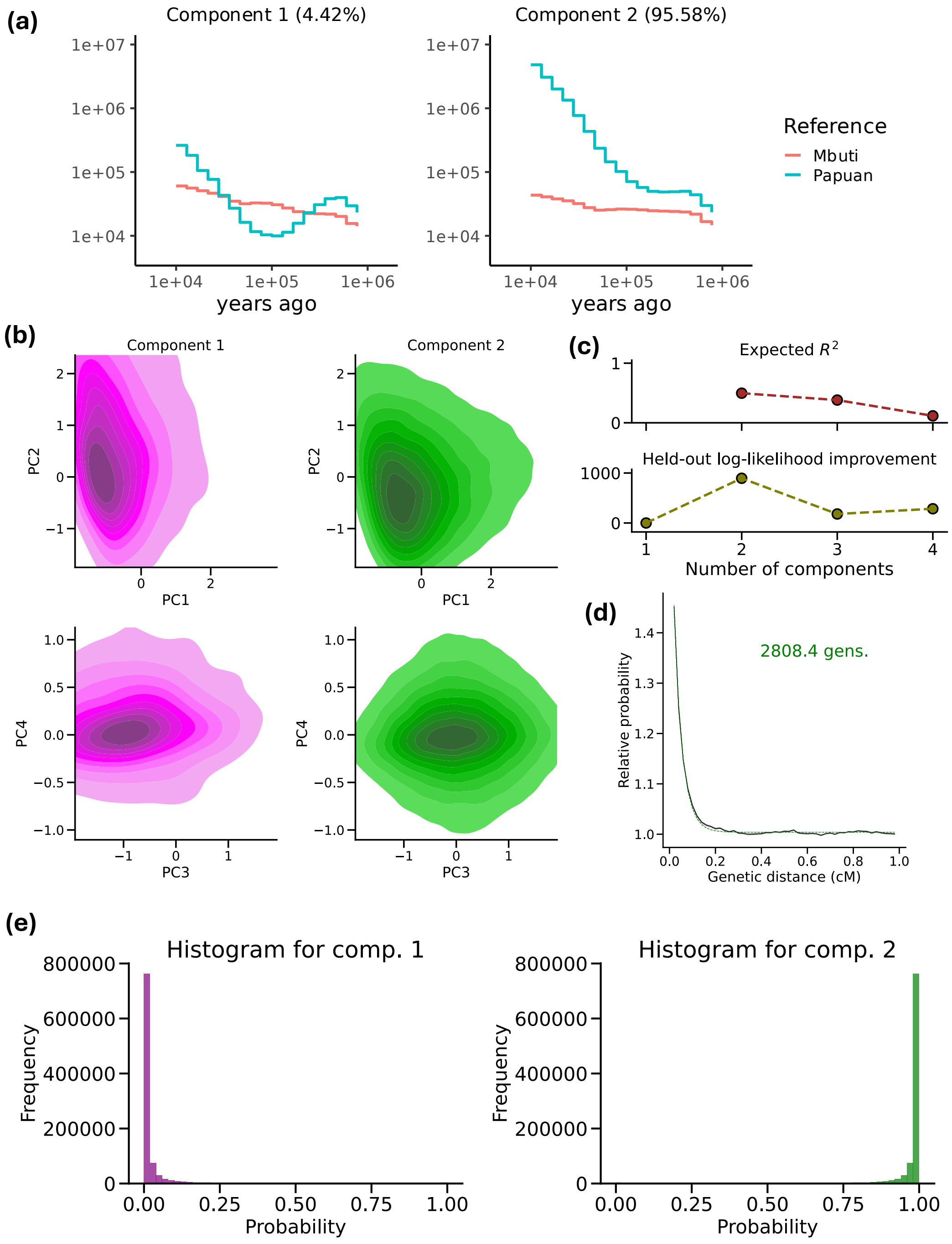

**Supplementary Figure 6**: **Decomposing unadmixed Mbuti-like individuals in simulations with Papuan-like as additional references.** Analogous to Supplementary Figure **2**, we show (a) inferred inverse coalescence rates, (b) PCA visualisation, (c) expected coefficient of determination and held-out log-likelihood improvement with varying number of components, and (d) coancestry curve. (e) Histogram of the inferred local ancestry posteriors. A low expected $R^2$, absence of clear clustering in PCA space, and unrealistically old inferred admixture dates jointly suggest that the fitted model is capturing noise rather than a genuine admixture signal.

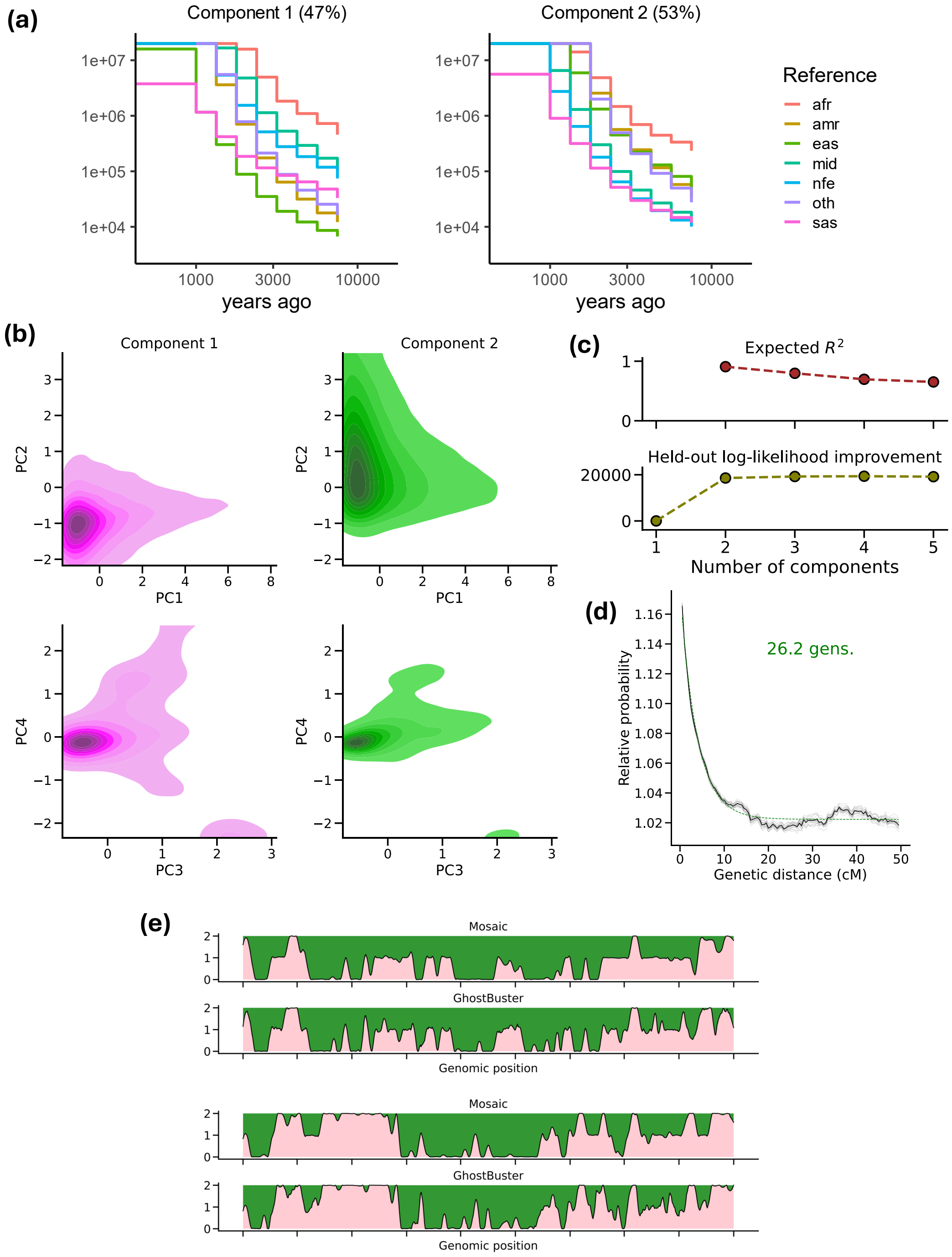

**Supplementary Figure 7**: **Decomposing Hazara individuals in Human Genome Diversity Project.** Analogous to Supplementary Figure **2**, we show (a) inferred inverse coalescence rates, (b) PCA visualisation, (c) expected coefficient of determination and held-out log-likelihood improvement with varying number of components, and (d) coancestry curve. (e) Local ancestry inference for GhostBuster vs. Mosaic. The diploid local ancestry in (e) is based on chromosome 1 for HGDP samples HGDP00102 and HGDP00103.

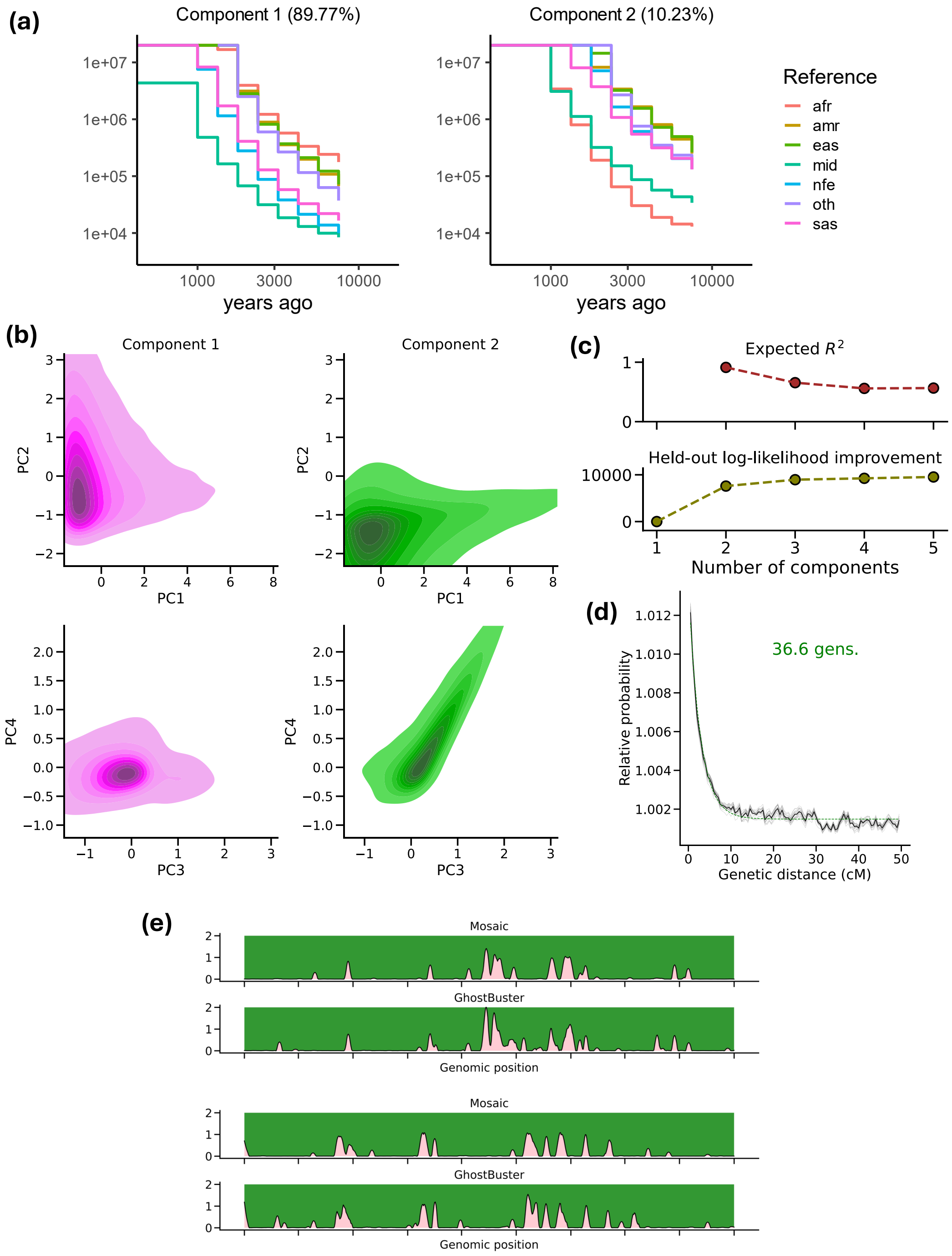

**Supplementary Figure 8**: **Decomposing Bedouin individuals in Human Genome Diversity Project.** Analogous to Supplementary Figure **2**, we show (a) inferred inverse coalescence rates, (b) PCA visualisation, (c) expected coefficient of determination and held-out log-likelihood improvement with varying number of components, and (d) coancestry curve. (e) Local ancestry inference for GhostBuster vs. Mosaic. The diploid local ancestry in (e) is based on chromosome 1 for HGDP samples HGDP00607 and HGDP00608.

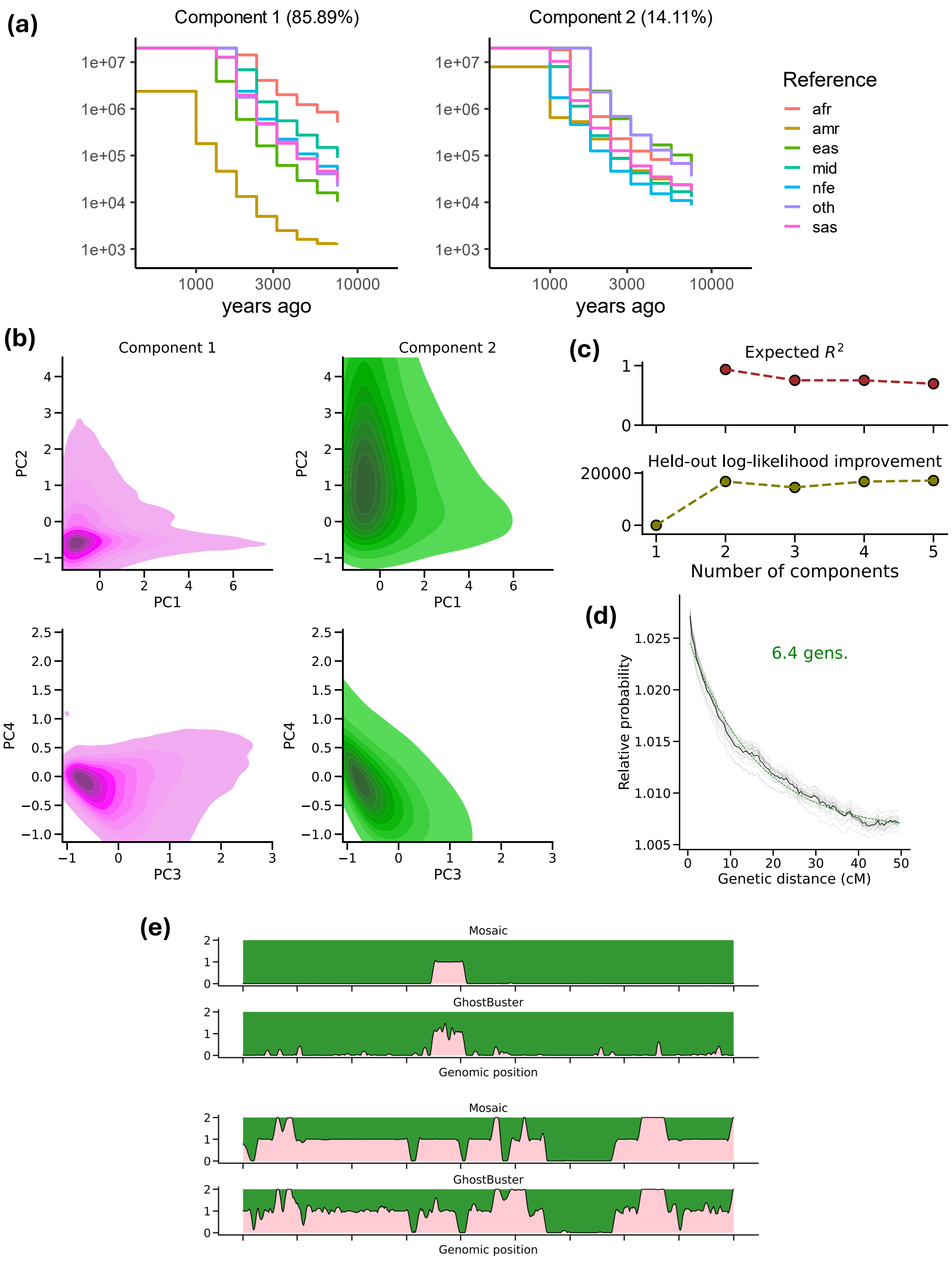

**Supplementary Figure 9**: **Decomposing Maya individuals in Human Genome Diversity Project.** Analogous to Supplementary Figure **2**, we show (a) inferred inverse coalescence rates, (b) PCA visualisation, (c) expected coefficient of determination and held-out log-likelihood improvement with varying number of components, and (d) coancestry curve. (e) Local ancestry inference for GhostBuster vs. Mosaic. The diploid local ancestry in (e) is based on chromosome 1 for HGDP samples HGDP00856 and HGDP00860.

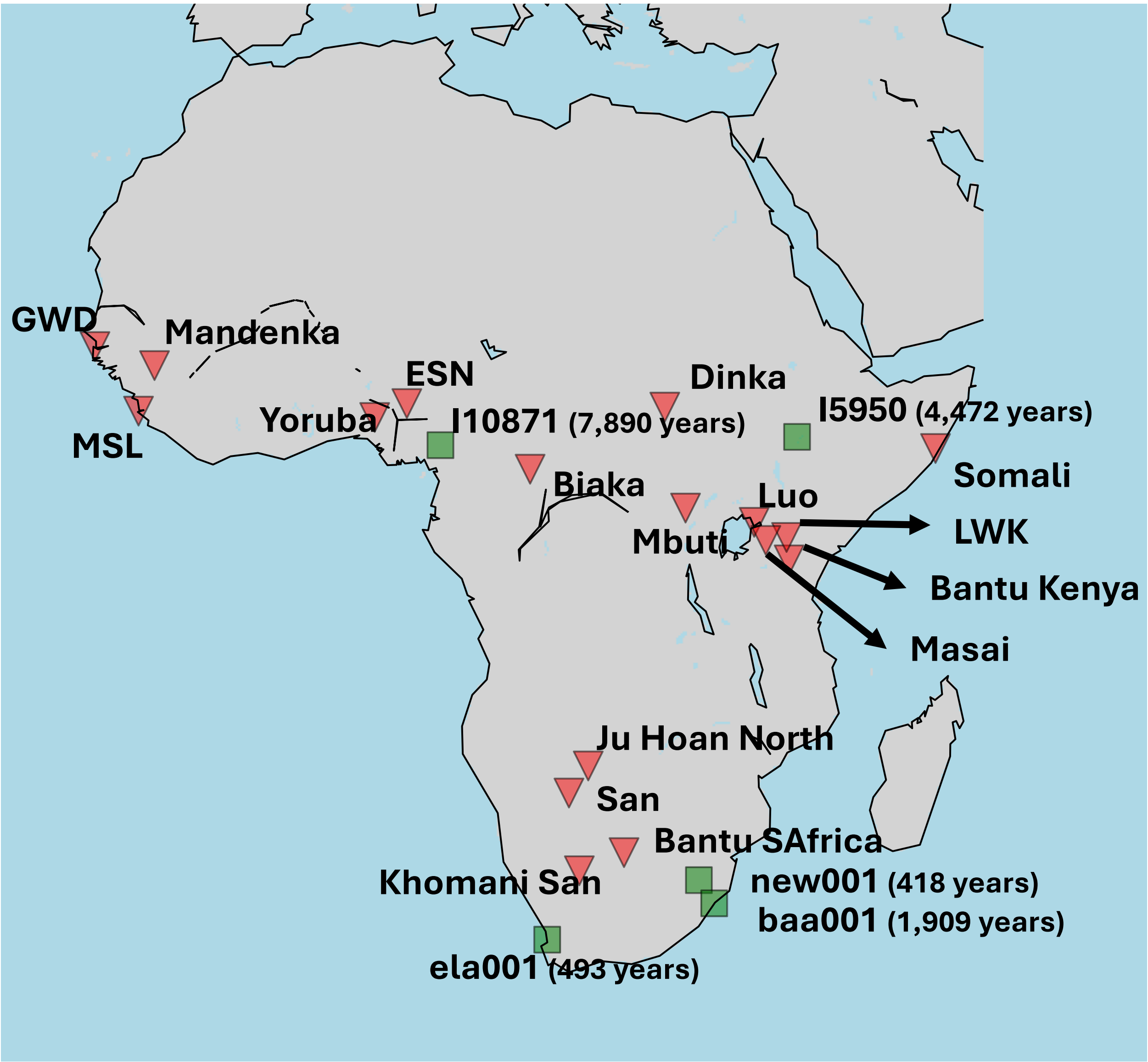

**Supplementary Figure 10**: **Modern and ancient African samples included in this study.** Geographic distribution of African populations analysed. Red triangles denote present-day populations, green squares indicate ancient individuals, with sample labels and radiocarbon dates (years before present) shown.

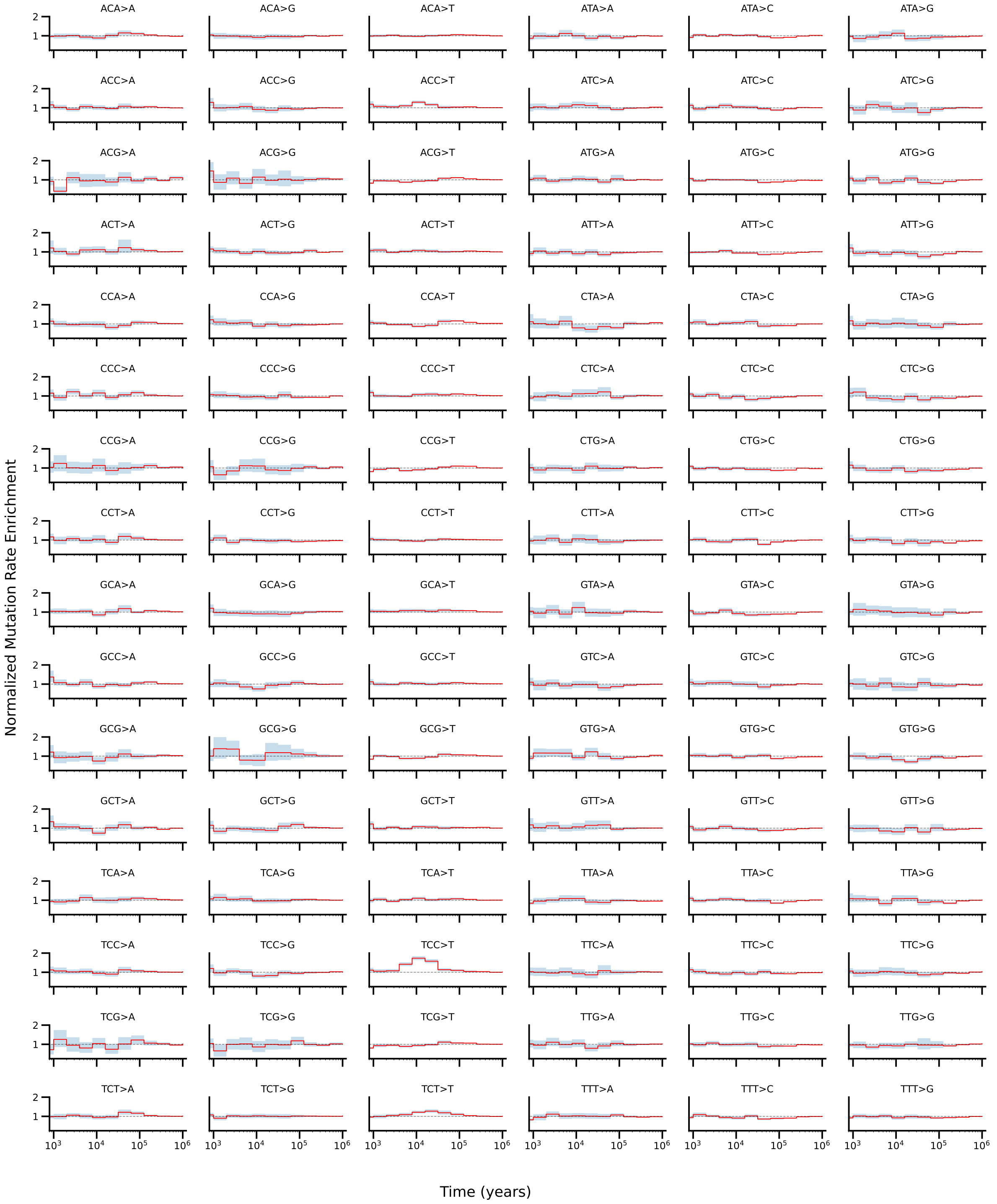

**Supplementary Figure 11**: **Normalised mutation rate enrichment corresponding to the Back-to-Africa event.** Normalised mutation rate enrichment across all 96 trinucleotide mutation types for Eurasian ancestry in Africans. Each panel corresponds to a specific mutation type, with enrichment values normalised relative to genome-wide expectations and shown as a function of allele age.

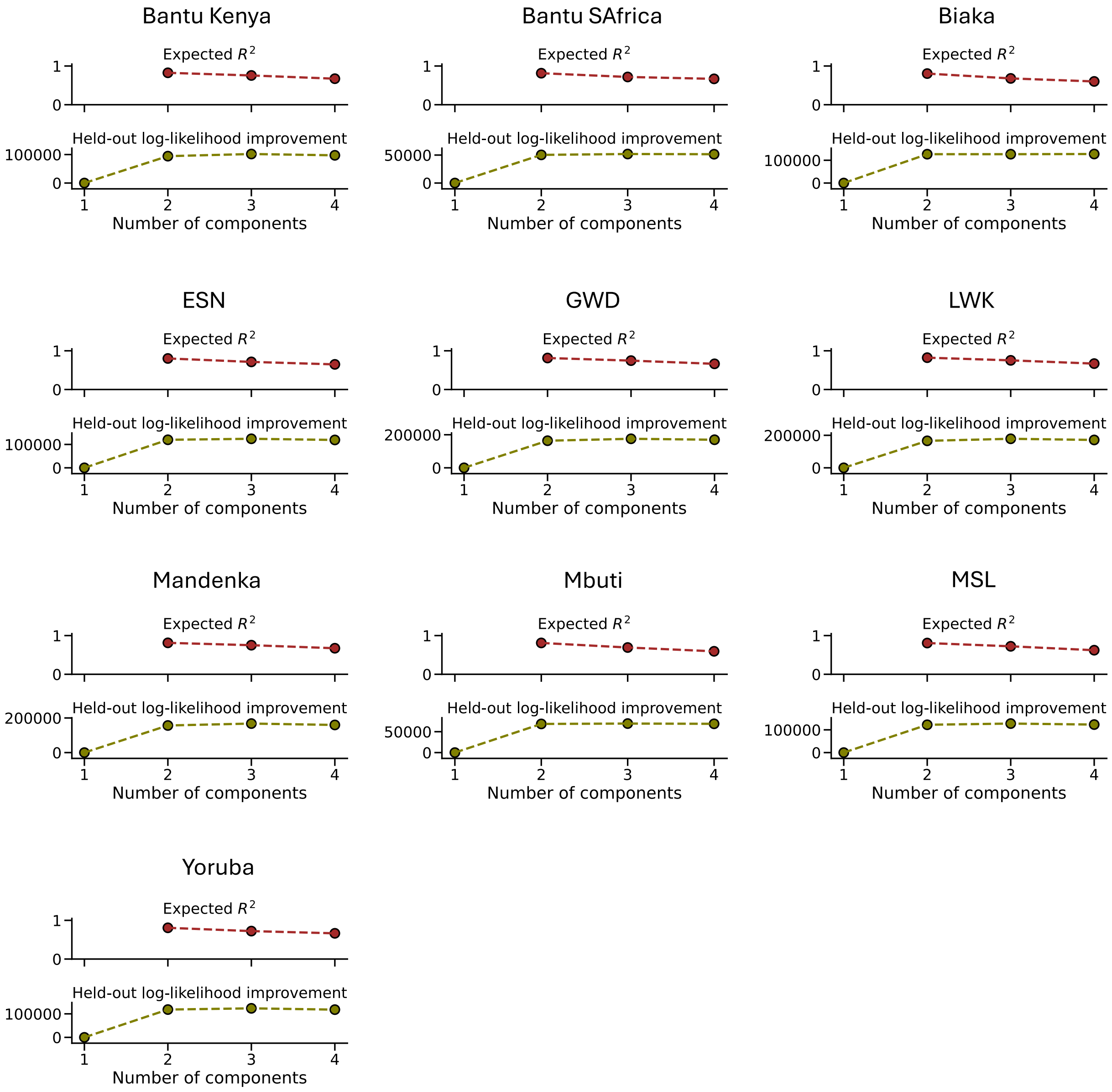

**Supplementary Figure 12**: **Model fit for the deep OOA/non-OOA admixture model across African populations.** Expected $R^2$ (top) and held-out log-likelihood improvement (bottom) for models with $k = 1 - 4$ admixture components across African populations. Increasing the number of components leads to modest changes in $R^2$ but substantial improvements in held-out log-likelihood from $k = 1$ to $k = 2$, with diminishing returns for additional components, supporting a two-component model as a parsimonious fit across populations.

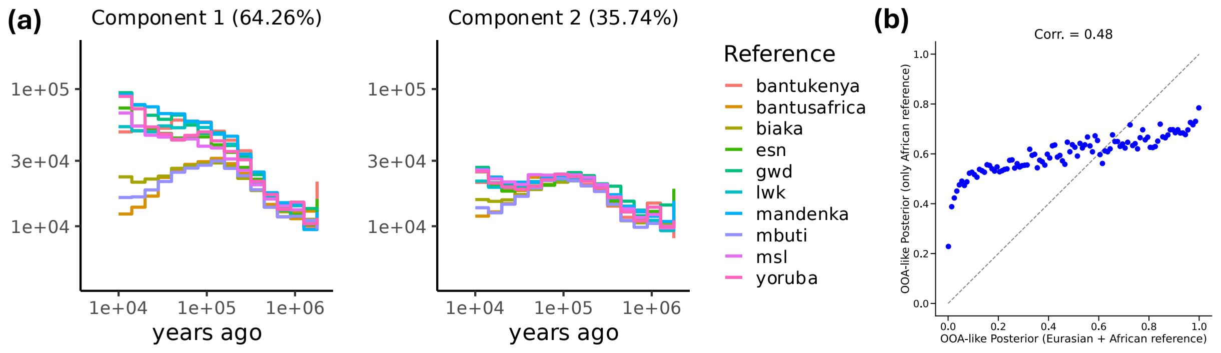

**Supplementary Figure 13**: **Comparison of GhostBuster runs with and without Eurasian reference samples.** Chromosome 1 of San individuals from the HGDP+1,000GP dataset was analysed, fixing the number of components to 2. The Eurasian samples were excluded from the reference panel before ancestry decomposition. (a) shows the inverse coalescence rates when only African populations were used as the reference, and (b) compares the correlation in local ancestry inferred with and without Eurasian samples in the reference panel. Plot (b) is based on binning the posterior probabilities into 500 equally spaced bins and calculating the average posterior within each bin. Pearson correlation coefficients (shown in panels) indicate the correlation between the raw ancestry calls.

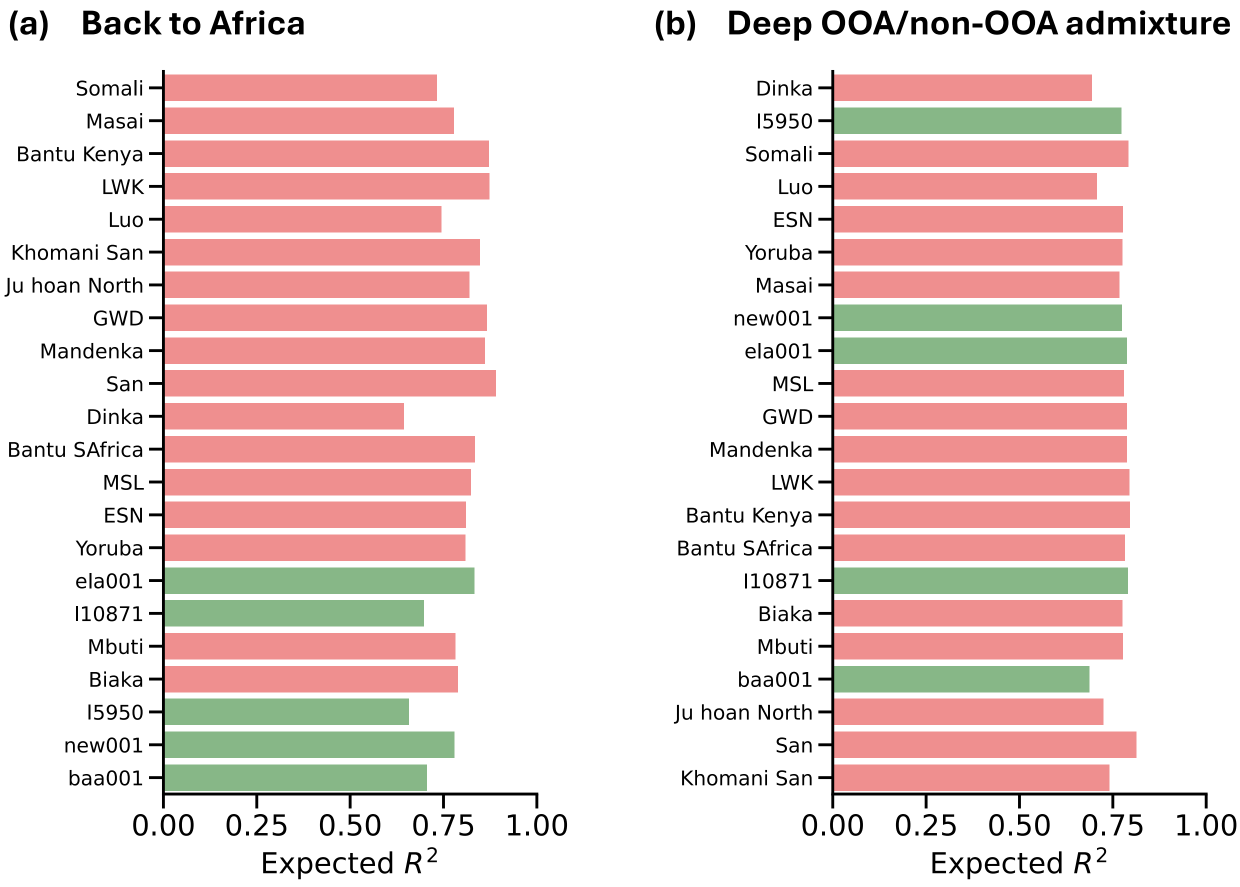

**Supplementary Figure 14**: **Expected coefficient of determination $R^2$ (defined in [19]) across African populations.** (a) Expected $R^2$ for the Back-to-Africa admixture event. (b) Expected $R^2$ for the deep OOA/non-OOA admixture event.

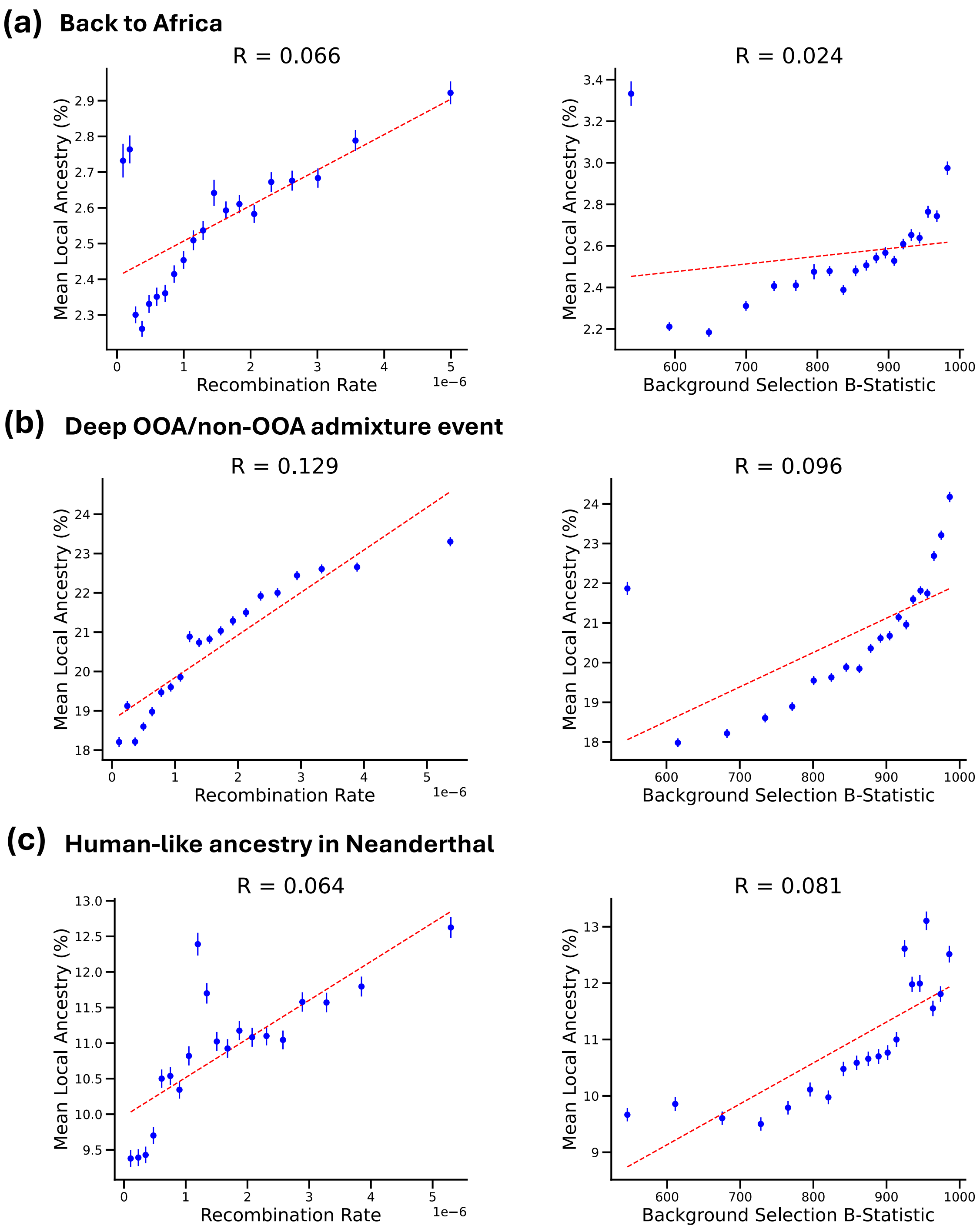

**Supplementary Figure 15**: **Correlation between GhostBuster-inferred local ancestry and genomic features.** Relationship between mean local ancestry and local (±100kb) recombination rate (left) and background selection (B-statistic [43]; right) for (a) Eurasian ancestry in Africans, (b) OOA-like ancestry in Africans, and (c) HA ancestry in Neanderthals. Points represent mean ± 95% confidence intervals across quintile genomic bins stratified by recombination rate or background selection, and dashed lines denote linear fits. Pearson correlation coefficients (shown in panels) indicate the correlation between the raw ancestry calls. Across all ancestry components, local ancestry shows little to no correlation with, and only a small change in mean values across, the genome-wide variation in recombination rate or background selection statistics, indicating that inferred ancestry patterns are not driven by variation in recombination rate or linked selection.

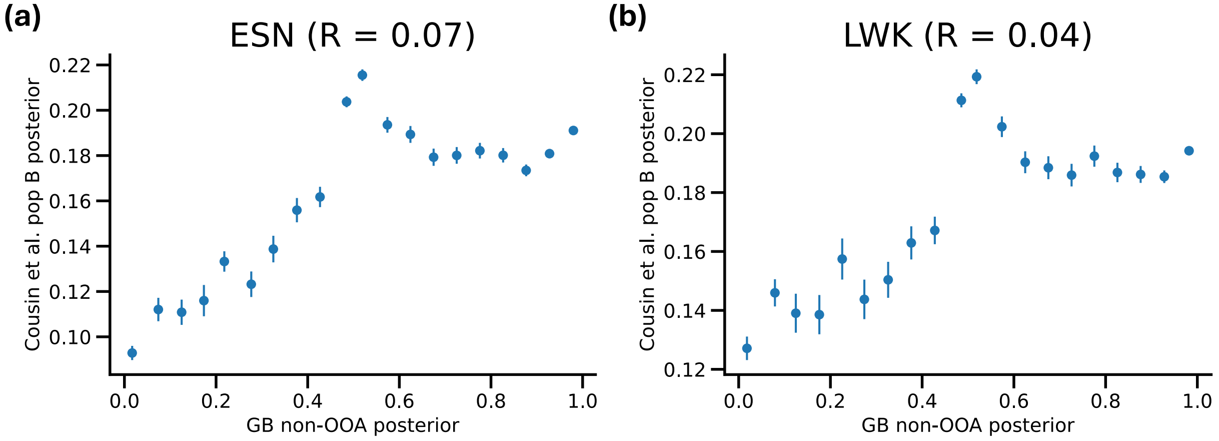

**Supplementary Figure 16**: **Correlation between GhostBuster-inferred non-OOA-like local ancestry and Cousins et al. [44] population B ancestry.** Comparison for two overlapping samples: (a) HG03515 (ESN) and (b) NA19017 (LWK). Points represent mean ± s.e. across genomic bins, and Pearson correlation coefficients (shown in panels) indicate the correlation between the raw ancestry calls.

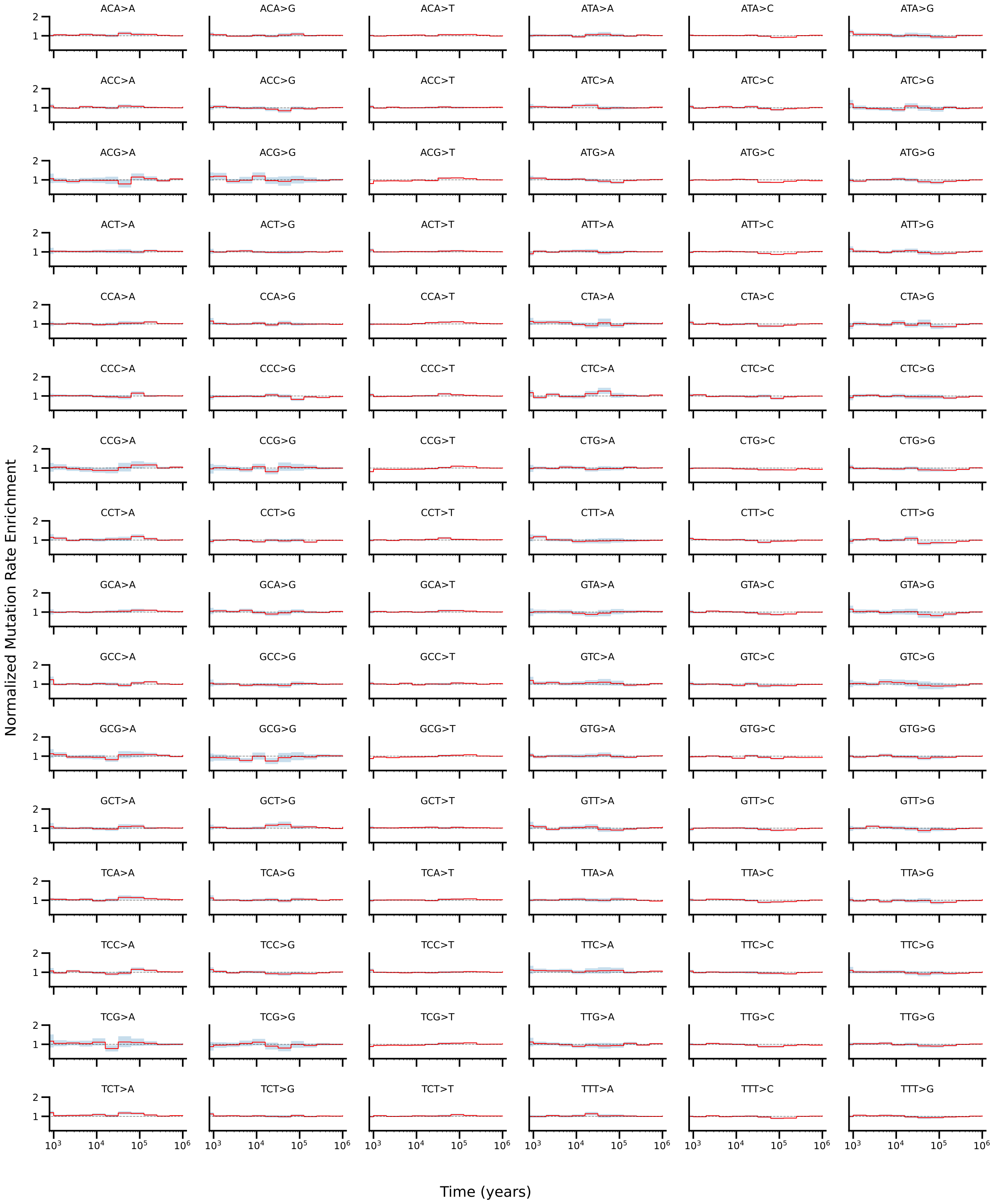

**Supplementary Figure 17**: **Normalised mutation rate enrichment corresponding to the deep OOA/non-OOA admixture event.** Normalised mutation rate enrichment across all 96 trinucleotide mutation types for OOA-like ancestry in Africans. Each panel corresponds to a specific mutation type, with enrichment values normalised relative to genome-wide expectations and shown as a function of allele age.

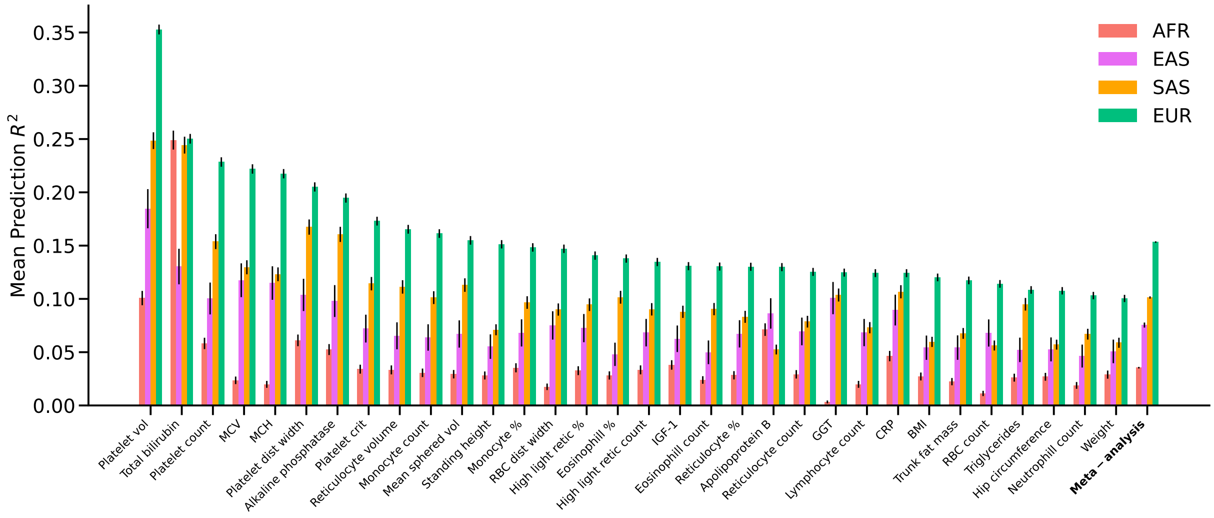

**Supplementary Figure 18**: **Predictive performance of polygenic scores across populations.** Proportion of variance explained ($R^2$) by polygenic scores across traits and populations. Bars represent the mean prediction $R^2$ with error bars indicating standard errors. Populations are grouped as African (AFR), East Asian (EAS), South Asian (SAS), and European (EUR). A meta-analysis across traits is shown on the right.

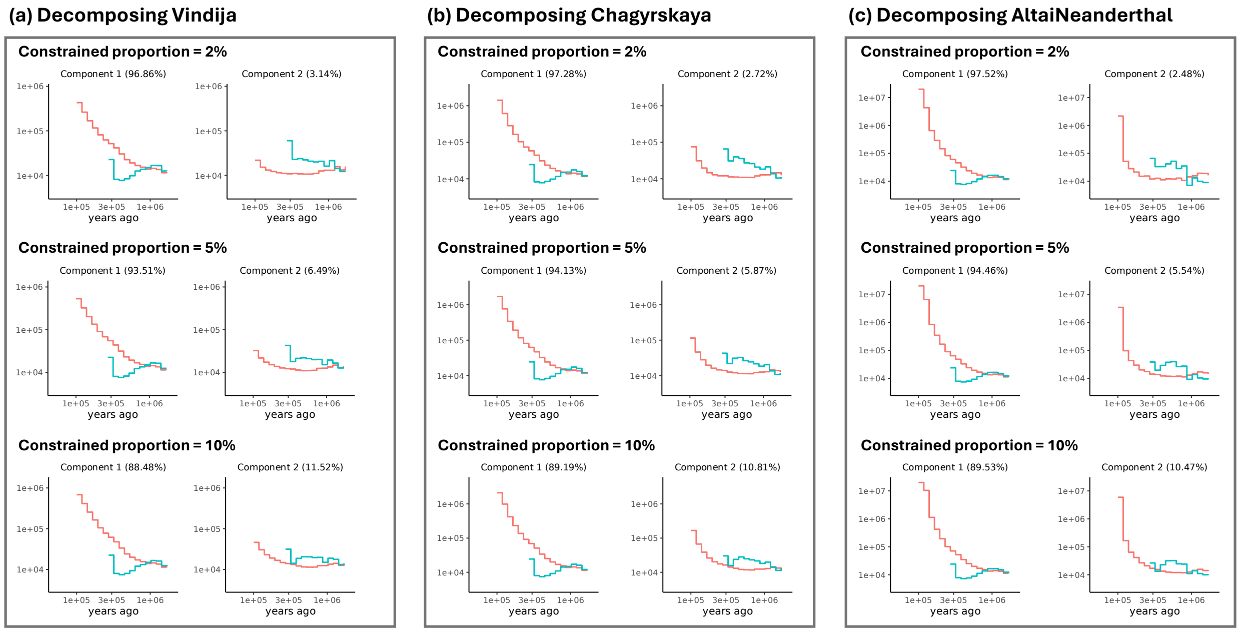

**Supplementary Figure 19**: **Decomposition of Neanderthals using GhostBuster.** Inverse coalescence rate trajectories inferred by GhostBuster for (a) Vindija, (b) Chagyrskaya, and (c) Altai Neanderthal. For each individual, results are shown under constraints fixing the proportion of the minor component (component 2) to 2%, 5%, and 10%. Corresponding component-specific trajectories and inferred mixture proportions are indicated within each panel.

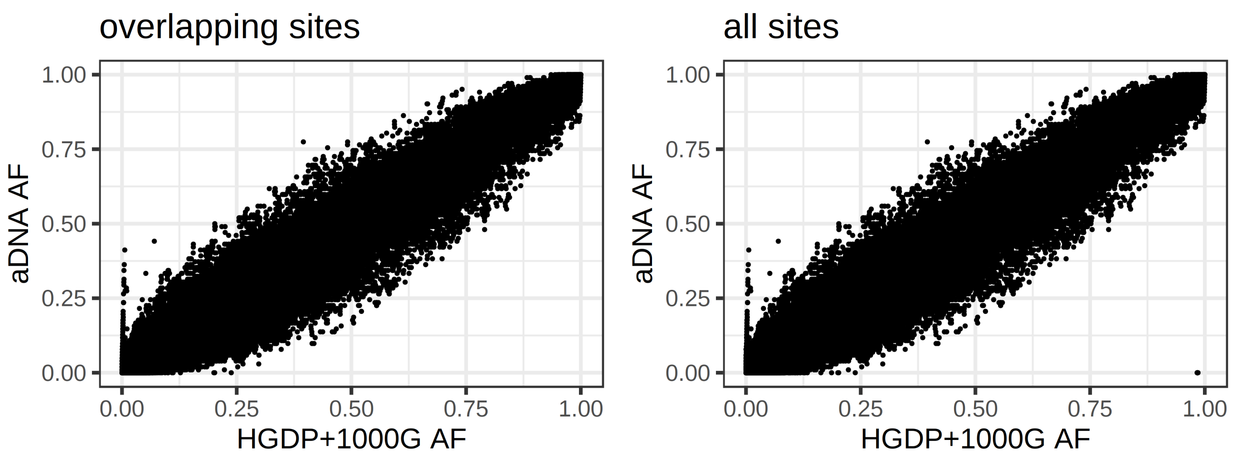

**Supplementary Figure 20**: **Comparison of allele frequencies between HGDP+**$1{,}000$**GP and ancient samples.** We analyse bi-allelic SNPs on chromosome 20 within the masked regions. The comparison includes "overlapping sites" between the two datasets, as well as a union ("all sites"), where we treat missing sites in either dataset as fixed for the reference allele.

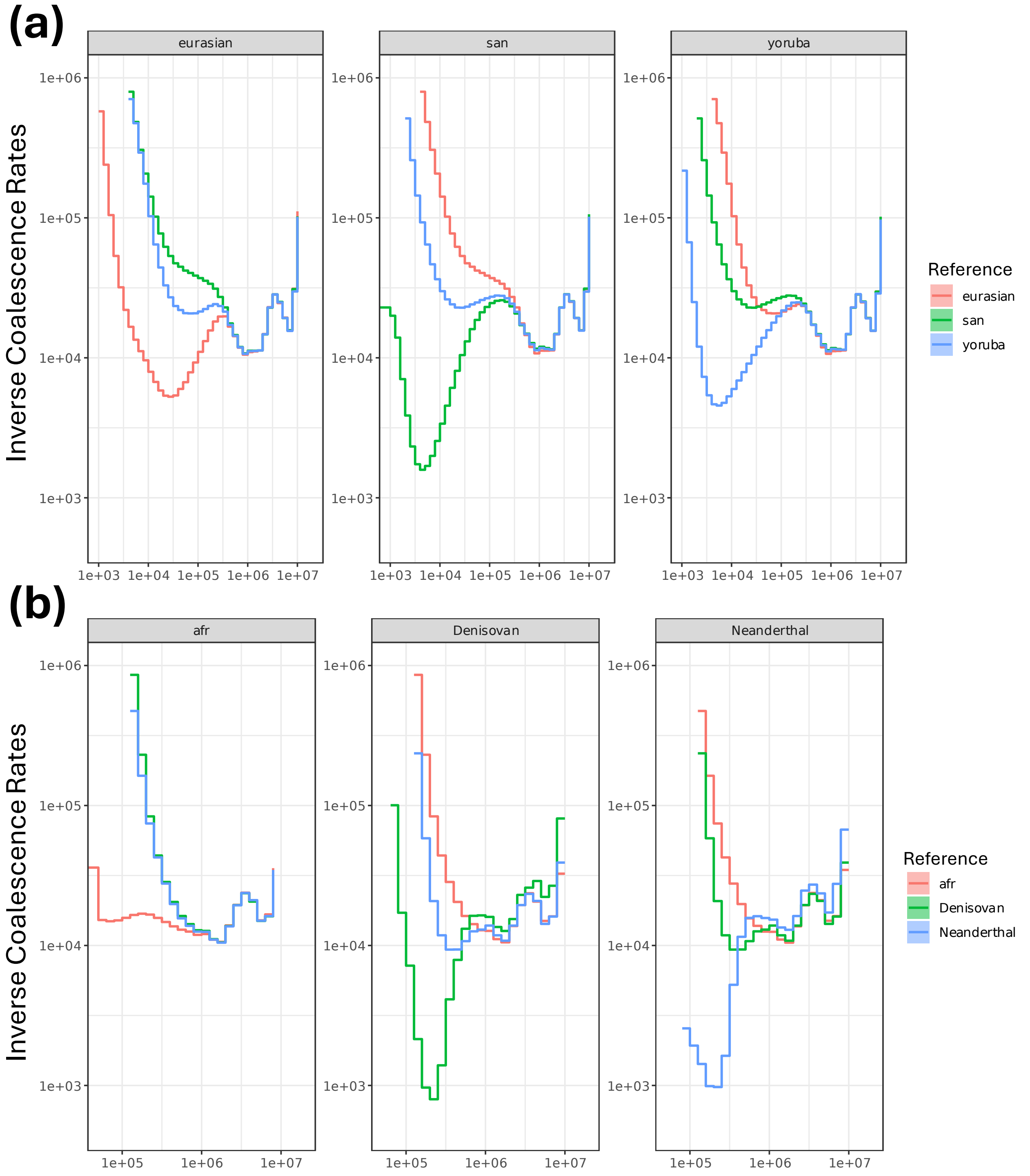

**Supplementary Figure 21**: **Genome-wide coalescence rate trajectories.** Inverse coalescence rate trajectories across time for different population groups. (a) Between modern human populations (Eurasian, San, Yoruba). (b) Between African populations (AFR) and archaic hominins (Denisovan, Neanderthal). Within each panel, curves represent inverse coalescence rates between the focal population (indicated in the panel title) and several reference groups.

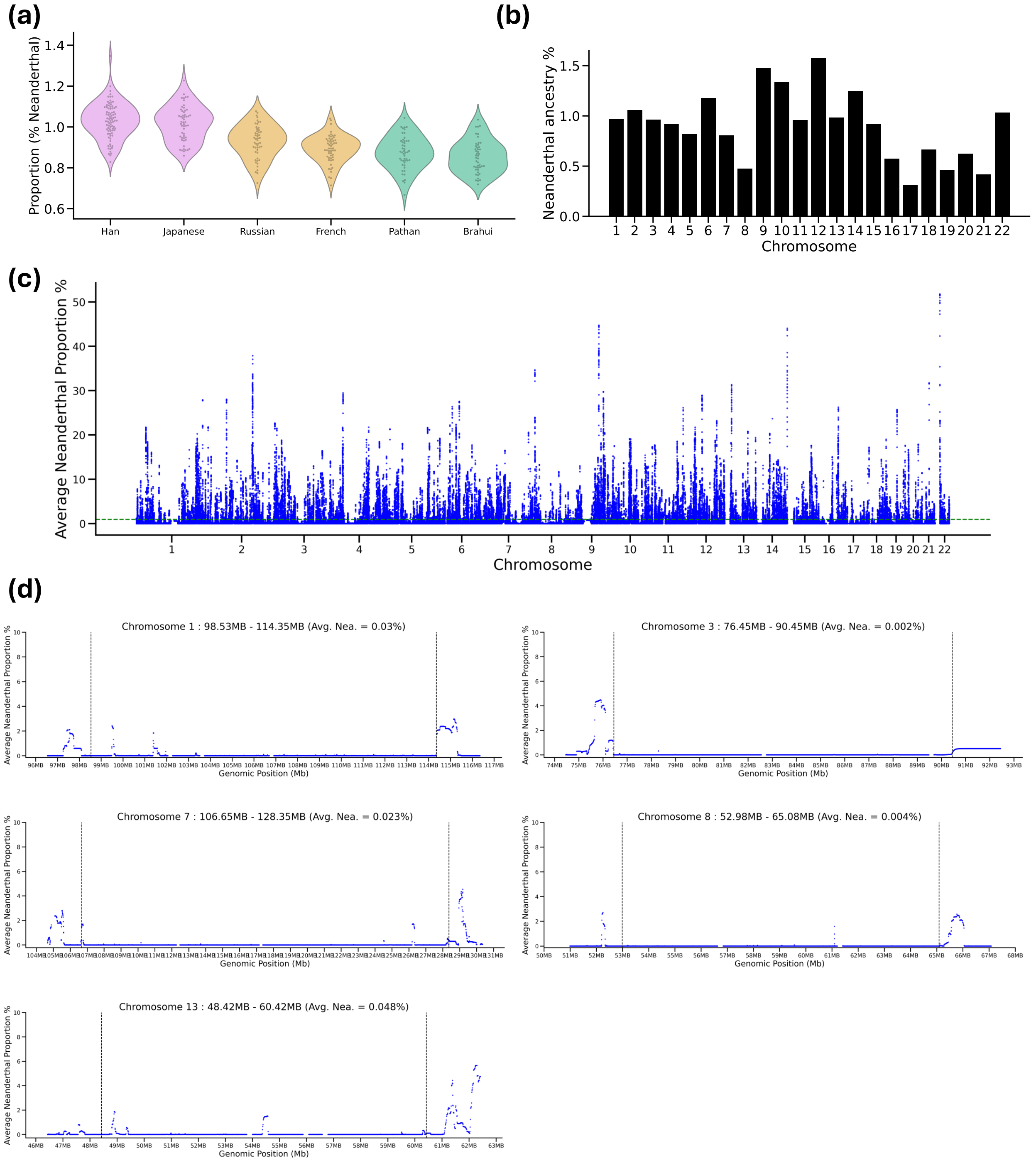

**Supplementary Figure 22**: **Neanderthal ancestry in non-African populations.** (a) Distribution of Neanderthal ancestry proportions across individuals from East Asian (Han, Japanese), European (Russian, French), and South Asian (Pathan, Brahui) populations. (b) Chromosome-wide estimates of Neanderthal ancestry, showing variation across autosomes. (c) Genome-wide distribution of Neanderthal ancestry proportions across genomic windows, with the dashed line indicating the genome-wide average. (d) Inferred Neanderthal ancestry across representative genomic regions corresponding to previously identified Neanderthal deserts [45, 46], showing depletion of introgressed ancestry within these intervals.

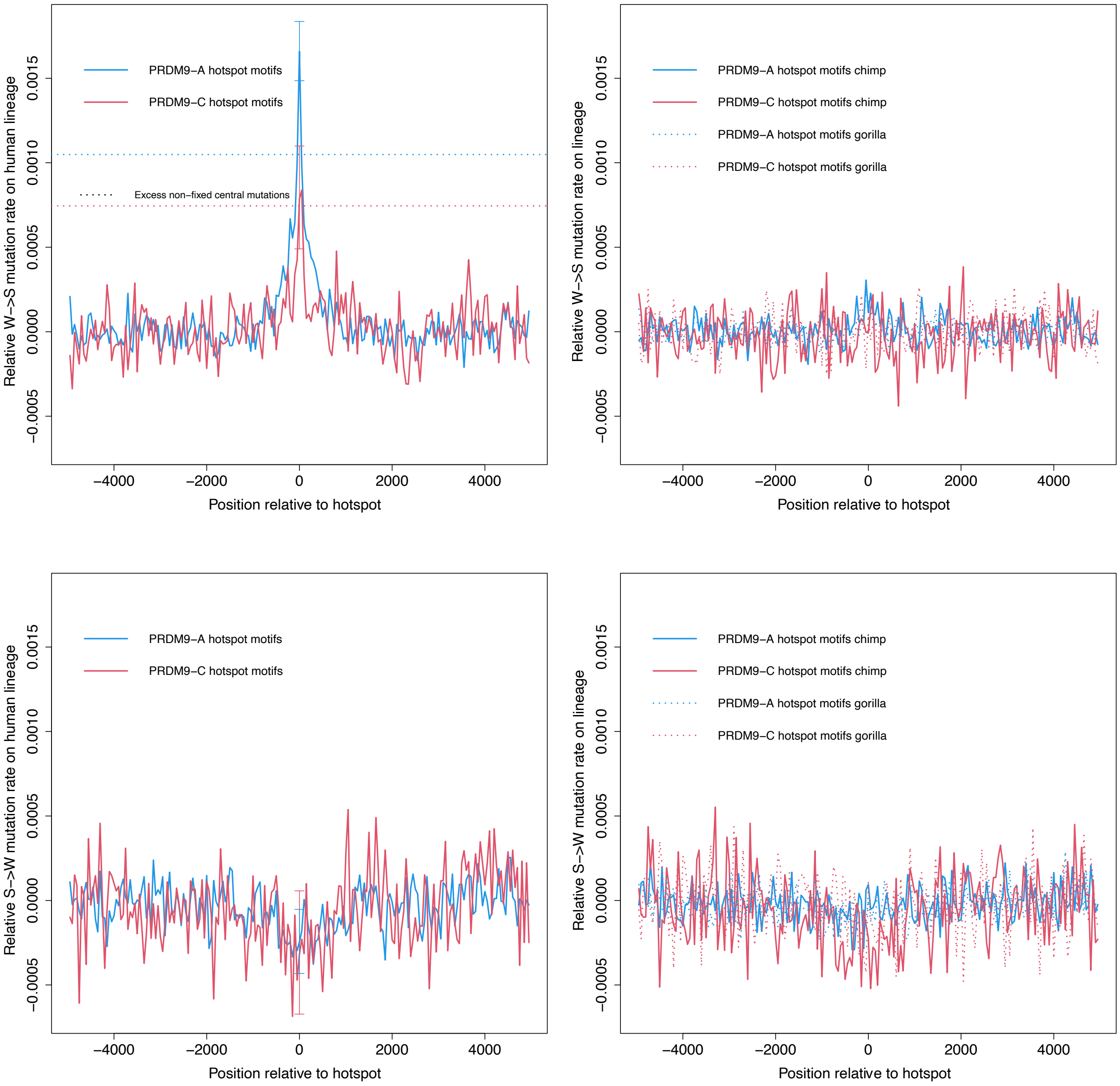

**Supplementary Figure 23**: **GCbGC signals around PRDM9 hotspots across human, chimp and gorilla lineages.** Relative mutation rates around PRDM9-A (blue) and PRDM9-C (red) binding motifs as a function of distance from the hotspot centre. Top panels show weak-to-strong (W→S) mutations, and bottom panels show strong-to-weak (S→W) mutations. Left panels correspond to the human lineage, and right panels to the chimpanzee and gorilla lineages. A pronounced enrichment of W→S mutations at hotspot centres is observed for PRDM9-A in humans, whereas PRDM9-C shows weaker signals. Dashed lines indicate baseline GCbGC estimated from mutations segregating in humans and archaics. The elevated signal at PRDM9-A relative to this baseline, but not PRDM9-C, suggests that the hominin population ancestral to both modern humans and archaic hominins primarily exhibited PRDM9-A activity with little or no PRDM9-C activity.

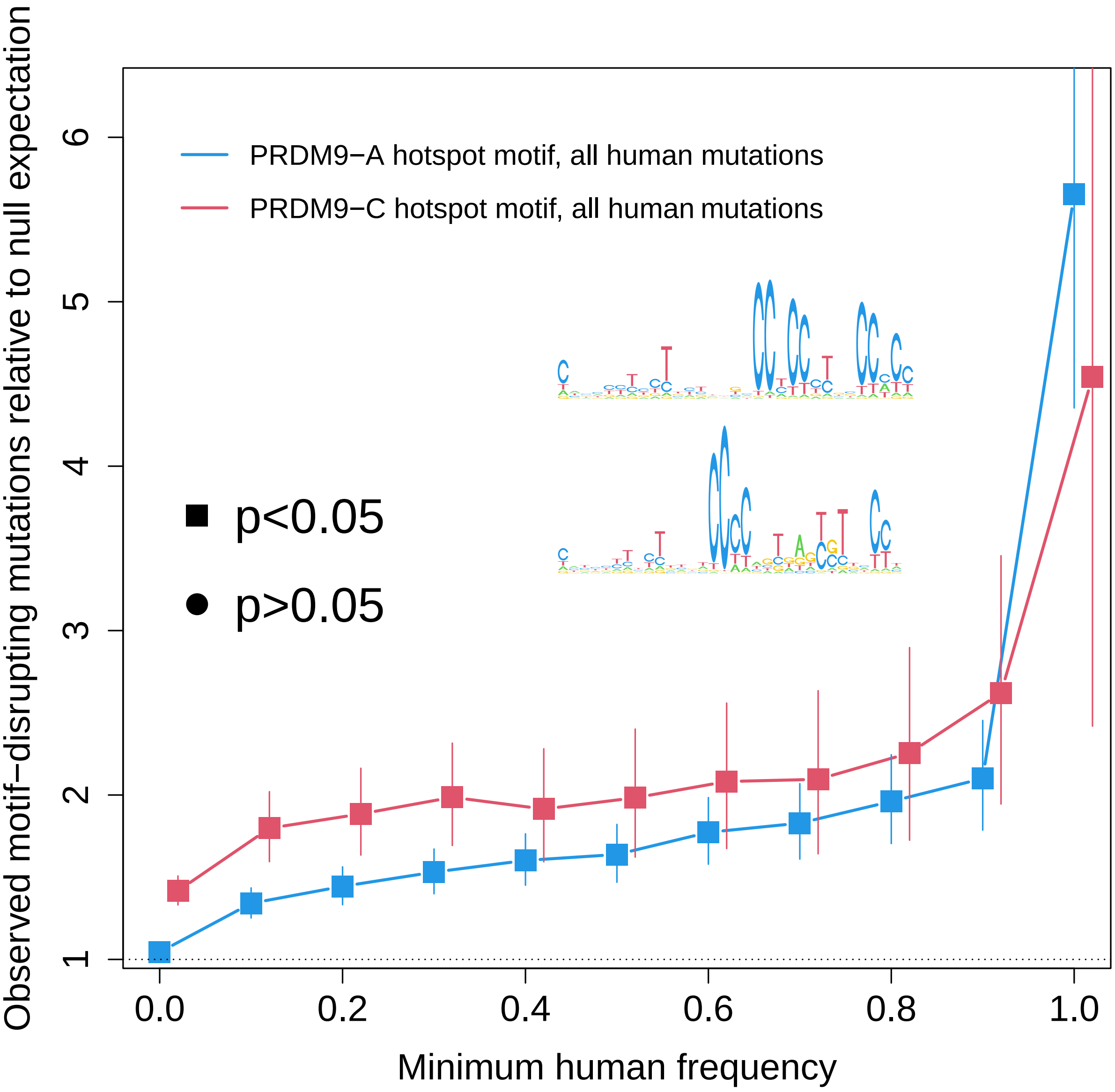

**Supplementary Figure 24**: **Evidence for hotspot death at PRDM9 binding motifs in humans.** Enrichment of motif-disrupting mutations relative to a local mutation rate null as a function of derived allele frequency in humans, shown separately for PRDM9-A (blue) and PRDM9-C (red) hotspot motifs. Points represent observed-to-expected ratios with error bars indicating 95% confidence intervals; filled squares denote nominal significance ($P < 0.05$). Enrichment increases with allele frequency, consistent with selection favouring mutations that disrupt PRDM9 binding sites. Insets show position weight matrices for PRDM9-A and PRDM9-C motifs.

### Supplementary Tables

**Supplementary Table 1**: **Ancient samples built in the genealogy.** Ancient sample names, relevant publication, sampling age (in years), country of origin and genomic coverage

| Sample Name | Publication | Sample Age | Country | Coverage |
|---|---|---|---|---|
| AltaiNeandertal | Prüfer et al. 2014, Nature | 110,450 | Russia | 52.0 |
| Vindija | Prüfer et al. 2017, Science | 41,950 | Croatia | 30.0 |
| Chagyrskaya | Mafessoni et al. 2020, PNAS | 80,000 | Russia | 27.0 |
| Denisova | Meyer et al. 2012, Science | 63,900 | Russia | 31.0 |
| WC1 | Broushaki et al. 2016, Science | 9,219 | Iran | 10.4 |
| PB675 | Cassidy et al. 2020, Nature | 5,416 | Ireland | 15.9 |
| JP14 | Cassidy et al. 2020, Nature | 5,497 | Ireland | 16.6 |
| SRA62 | Cassidy et al. 2020, Nature | 5,979 | Ireland | 13.2 |
| Yamnaya | Damgaard et al. 2018, Science | 4,890 | Kazakhstan | 25.2 |
| Ust˙Ishim | Fu et al. 2014, Nature | 45,020 | Russia | 42.0 |
| SF12 | Gunther et al. 2018, PLoS Biology | 8,895 | Sweden | 65.2 |
| KK1 | Jones et al. 2015, Nature Communications | 9,678 | Georgia | 11.3 |
| I0018 | Lazaridis et al. 2014, Nature | 7,140 | Germany | 19.0 |
| I0001 | Lazaridis et al. 2014, Nature | 8,025 | Luxembourg | 20.1 |
| I10871 | Lipson et al. 2022, Nature | 7,890 | Cameroon | 18.5 |
| I5950 | Llorente et al. 2015, Science; Lipson et al. 2022, Nature | 4,472 | Ethiopia | 11.3 |
| Klein7 | Marchi et al. 2022, Cell | 7,122 | Austria | 11.3 |
| Asp6 | Marchi et al. 2022, Cell | 7,524.5 | Austria | 12.1 |
| Ess7 | Marchi et al. 2022, Cell | 6,975 | Germany | 12.3 |
| Herx | Marchi et al. 2022, Cell | 7,078.5 | Germany | 11.5 |
| Dil16 | Marchi et al. 2022, Cell | 7,116.5 | Germany | 10.6 |
| Nea2 | Marchi et al. 2022, Cell | 8,098 | Greece | 12.5 |
| Nea3 | Marchi et al. 2022, Cell | 8,183.5 | Greece | 11.6 |
| VC3-2 | Marchi et al. 2022, Cell | 7,495.5 | Serbia | 11.2 |
| STAR1 | Marchi et al. 2022, Cell | 7,532.5 | Serbia | 10.6 |
| LEPE52 | Marchi et al. 2022, Cell | 7,812 | Serbia | 12.4 |
| LEPE48 | Marchi et al. 2022, Cell | 7,939.5 | Serbia | 10.9 |
| Bar25 | Marchi et al. 2022, Cell | 8,294.5 | Turkey | 12.7 |
| AKT16 | Marchi et al. 2022, Cell | 8,547.5 | Turkey | 12.3 |
| new001 | Schlebusch et al. 2017, Science | 418 | South Africa | 11.1 |
| ela001 | Schlebusch et al. 2017, Science | 493 | South Africa | 10.5 |
| baa001 | Schlebusch et al. 2017, Science | 1,909 | South Africa | 12.9 |
| Yana1 | Sikora et al. 2019, Nature | 31,950 | Russia | 24.3 |
| Sunghir3 | Sikora et al. 2017, Science | 34,093 | Russia | 12.0 |

**Supplementary Table 2**: **Population-level estimates of back-to-Africa (BTA) and out-of-Africa (OOA) ancestry proportions across African populations.** Populations are ordered by decreasing OOA proportion. Latitude and longitude indicate the approximate geographic sampling location for each population, and the number of samples corresponds to the individuals included in the analysis. BTA prop. represents the estimated proportion of ancestry attributable to back-to-Africa gene flow, while OOA prop. represents the proportion of ancestry tracing to the out-of-Africa-like source. Eurasian gene flow was removed to calculate OOA-like proportions. Proportions are reported as percentages and rounded to one decimal place.

| **Population** | **Latitude** | **Longitude** | **# samples** | **BTA prop. (%)** | **OOA prop. (%)** |
|---|---|---|---|---|---|
| Somali | 5.600000 | 48.300000 | 2 | 23.5 | 33.3 |
| I5950 | 6.498896 | 37.612088 | 1 | 0.8 | 30.1 |
| Masai | -1.500000 | 35.200000 | 2 | 15.2 | 27.6 |
| Dinka | 8.794444 | 27.400000 | 3 | 5.4 | 24.8 |
| Luo | -0.100000 | 34.300000 | 2 | 5.9 | 21.3 |
| new001 | -27.722953 | 30.031450 | 1 | 1.6 | 21.2 |
| ela001 | -32.317362 | 18.318878 | 1 | 2.4 | 19.8 |
| I10871 | 5.858669 | 10.077854 | 1 | 1.1 | 17.7 |
| Yoruba | 7.995095 | 5.000000 | 21 | 2.4 | 17.7 |
| ESN | 9.066667 | 7.483333 | 20 | 2.5 | 17.6 |
| BantuKenya | -3.000000 | 37.000000 | 11 | 4.7 | 17.5 |
| LWK | -1.266667 | 36.800000 | 20 | 4.6 | 17.4 |
| Mandenka | 12.000000 | -12.000000 | 21 | 3.7 | 17.4 |
| GWD | 13.466667 | -16.600000 | 20 | 3.8 | 17.2 |
| MSL | 8.484450 | -13.234450 | 20 | 2.7 | 17.0 |
| BantuSAfrica | -25.600000 | 24.250000 | 8 | 2.8 | 16.8 |
| Biaka | 4.000000 | 17.000000 | 22 | 1.8 | 12.8 |
| Mbuti | 1.000000 | 29.000000 | 12 | 1.8 | 10.5 |
| San | -21.000000 | 20.000000 | 6 | 3.3 | 7.2 |
| KhomaniSan | -27.000000 | 20.800000 | 2 | 4.6 | 6.8 |
| JuHoanNorth | -18.909314 | 21.488112 | 4 | 4.7 | 6.6 |
| baa001 | -29.509996 | 31.214876 | 1 | 0.5 | 6.3 |

**Supplementary Table 3**: **(See Excel sheet.) Summary of key statistics and results for PRDM9-based analyses.** (a) Adaptive introgression candidates; (b) summary of PRDM9 allele inference; (c) testing for hotspot death; (d) hotspot death and introgression analyses.
